# Using adaptive evolutionary signatures to screen candidate regulatory genes and cis-elements associated with Kranz development in monocots

**DOI:** 10.1101/2024.02.22.581542

**Authors:** Angeo Saji, Gopikrishnan Bijukumar, Vivek Thakur

## Abstract

C_4_ plants exhibit higher photosynthetic efficiency under high temperatures and low water availability due to their unique anatomical and biochemical adaptations, but the evolutionary mechanisms and regulatory components underlying their Kranz development remain partly known. If C_4_ traits evolved through adaptive changes, its developmental regulators should show detectable evolutionary signatures in C_4_ lineages. We investigated the adaptive evolutionary changes in both protein sequences and *cis*-motifs upstream of candidate genes differentially expressed during Kranz anatomy establishment in C_4_ grasses. We identified signatures of convergent evolution in candidate genes and their upstream regions by comparing C_4_ orthologs with their C_3_ counterparts, and enriched non-syntenic motifs present upstream of C_4_ orthologs through phylogeny-aware analysis. Out of 191 candidate genes analyzed, ten orthogroups containing maize orthologs of *EREB160, DOF11, bHLH116, MADS9, bHLH33/bHLH105, SCRO1/SCR2, CADTFR3, THX8, C3H28*, and *SHR1/SHR2* showed positive selection pressure specific to C_4_ orthologs. Among them, *DOF11, SCRO1/SCR2,* and *SHR1/SHR2* have previously been implicated in Kranz regulation by previous studies. Additionally, we identified a conserved position mutation present in *bHLH116* across C_4_ lineages, which may influence its DNA-binding domain. Moreover, we identified 39 annotated non-syntenic/C_4_-shifted DNA motifs, upstream of 28 gene orthologs, enriched in C_4_ species. Notably, genes with these motifs did not intersect with the genes under positive selection, suggesting these pathways evolved separately without much crosstalk, nevertheless, a few putative regulatory-network-motifs were observed. These motifs are potential additions to the limited Kranz-specific motifs and can aid in predicting new putative regulators after experimental validation.

**Significance statement:** C_4_ plants evolved as an adaptive response through anatomical and biochemical changes in their C_3_ relatives. Unlike the biochemical changes, the regulators/drivers of anatomical changes and the nature of underlying changes remains only partially understood. Using signatures of adaptive evolution, the current study screened candidate regulatory factors and sequences, including a few previously known and many novel candidates. These findings complement the existing experimental knowledge and advance the discovery of the gene regulatory network underlying the anatomical changes associated with C_4_ evolution.

## Introduction

C_4_ photosynthesis is one of the best-known examples of convergent evolution in plants and has evolved more than 60 times independently across angiosperms (Sage et al., 2011; Sage, 2016). It evolved as an adaptation to declining atmospheric CO₂ and increased photorespiration during the Oligocene and Miocene, allowing plants to concentrate CO₂ at the site of Rubisco and maintain photosynthetic efficiency (Ehleringer JR, et.al., 1991). The most commonly seen C_4_ strategy revolves around the spatial separation of the metabolic reactions into two different cell types, most commonly organized into a characteristic wreath-like structure around the veins known as Kranz anatomy (Brown, 1975). Because Kranz anatomy provides the anatomical framework required for C_4_ metabolism, understanding how this structure is established and regulated is central to explaining repeated C_4_ evolution and is important for efforts to engineer C_4_ traits into C_3_ crops.

Two structural features, consistently associated with all variations of Kranz anatomy, are radial patterning of mesophyll cells around the bundle sheath cells and the establishment of a much denser vein network in the leaf, both established during early leaf development (Russel et al., 1985; Nelson, 2011). In monocots, C_4_ species uniquely form rank-2 intermediate veins, whereas C_3_ monocots lack these additional vascular ranks (Külahoglu et al., 2014), suggesting that C_4_ evolution involves modification of existing developmental programs controlling vein differentiation. Radial patterning in C_4_ leaves has been linked to pathways that also regulate root patterning, particularly the *SHORTROOT–SCARECROW (SHR/SCR)* pathway. Mutant analyses and comparative studies implicate *SHR/SCR* and associated regulators such as *AtJACKDAW*, *ZmRAVEN1*, and *ZmIDD14* as candidate contributors to Kranz development (Fouracre et al., 2014; Coelho et al., 2018; Kumar and Kellogg, 2019). Large-scale transcriptomic comparisons between C_3_ and C_4_ maize leaves have identified many genes differentially expressed during early development (Wang et al., 2013; Liu et al., 2022), but these studies have not resolved which regulators are required for establishing Kranz anatomy (Wang et al., 2017). A recent spatial transcript abundance profiling study reported the spatio-temporal regulation of a few transcription factors, such as *TOO MANY LEAVES 1* (*TML1)* and *SHR1*, in minor (rank-1) intermediate veins (Perico et al., 2024); however, there was no clarity on the regulation of rank-2 intermediate veins, as well as how the expression pattern differs from C_3_ monocots. As a result, the regulatory hierarchy underlying the Kranz formation remains partly unclear.

Besides the transcription factors, *cis* regulatory elements are another unexplored layer of Kranz anatomy regulation. These noncoding regions regulate when and where genes are expressed and are known to drive cell-type-specific expression of several C_4_ metabolic enzymes. For example, upstream regulatory regions of the C_4_ phosphoenolpyruvate carboxylase gene are sufficient to confer mesophyll-specific expression and show conservation across C_4_ grasses (Akyildiz et al., 2007; Gupta et al., 2020). While metabolic specialization in C_4_ plants has been linked to regulatory evolution, much less is known about how *cis*-regulatory architecture contributes to the development of Kranz anatomy itself. Though, there have been multiple studies on regulatory element discovery for (Bundle-sheath or Mesophyll) cell-specific expression, which indicated that such motifs were often co-opted from existing ones from the ancestral C3 genomes rather than being acquired *de novo* (Williams et al., 2016; Reyna-Llorens et al., 2018; Swift et al., 2024).

It is now established that C_4_ photosynthesis evolved largely through recruitment and modification of pre-existing C_3_ genes rather than the appearance of new enzymes (Doebley & Lukens, 1998; Christin et al., 2013). Gene duplication followed by biased paralog recruitment created constrained evolutionary pathways, and strong evidence of parallel adaptation has been documented for enzymes such as phosphoenolpyruvate carboxylase (Christin et al., 2007). If Kranz anatomy also evolved through constrained routes, its developmental regulators should show detectable evolutionary signatures in C_4_ lineages, *i.e.*, accumulated nonsynonymous mutations, or motifs enriched in upstream regions of C_4_ orthologs of these regulators. However, most evolutionary studies have focused on metabolic enzymes, as a result research on discovery of transcription factors and regulatory motifs is lagging behind.

In the current study, we aim to identify whether the transcription factors, which are active during the early leaf development when vein patterning is being established, acquired any changes either in its protein sequence or in its *cis*-regulatory elements. We obtained ortholog clusters of such transcription factors along with a control gene set, and compared C_4_ and C_3_ orthologs for lineage-specific adaptive evolution. Evidence of positive selection in C_4_ lineages was found in a limited number of TFs (13 candidates from 10 orthogroups), including *SHR/SCR* and *DOF11,* which are known regulators of Kranz anatomy. The remaining 8 candidates are novel putative candidates for Kranz regulation. In contrast, upstream regulatory regions of the candidate genes showed a large number of motifs enriched in both C_4_ orthologs and control genes. After filtering these motifs in a phylogeny-aware analysis, we found that motifs showing non-syntenic behaviour were enriched in candidate transcription factors while being disproportionately low in the control geneset, suggesting that this set contains motifs possibly involved in C_4_ anatomy establishment. This observation supports the hypothesis that motifs which are recruited for C_4_ anatomy establishment have also been co-opted from existing *cis*-motifs, rather than being completely novel in nature. Together, these observations support a model in which the evolution of Kranz anatomy is driven by both targeted protein-level adaptations and widespread changes in *cis*-regulatory architecture, with limited overlap between these mechanisms.

## Results

### 191 Transcription factors differentially expressed during *Kranz*-anatomy development were selected as candidates

We wanted to examine the regulatory genes, showing differential expression during Kranz anatomy development, for evidence of C_4_-specific positive selection or motif acquisition/modification. The workflow followed is given in Fig. 1. The putative candidate genes were selected mainly from two studies in maize, namely Wang et al. (2013) and Liu et al. (2022); the former contributed putative positive and negative candidate regulators of Kranz development based on differentially expressed genes (DEGs) between husk and foliar leaf primordia, and the latter contributed DEGs between pre-Bundle-sheath cells and pre-Mesophyll cells. We started with 191 candidate genes from both of these studies, among which 9 candidates were present in both studies, as a test-set. Besides, a control set consisted of 50 genes, that had no known association with Kranz anatomy, and had a neutral or same expression profile in both husk and foliar leaf primordia in maize (Supp. Table S1, S2, S3). Among the 191 genes of the test-set, were a few transcription factors strongly implicated in Kranz development regulation, namely *SCARECROW, SHORTROOT, EREB114, TML(WIP6)*, and *DOF11*. Coincidentally, *TML* was implicated in both differential expression between husk and foliar leaf primordia and between pre-Bundle-sheath cells and pre-Mesophyll cells, but the other four were selected because of differential expression between foliar and husk primordial tissues in maize.

**Fig. 1:**
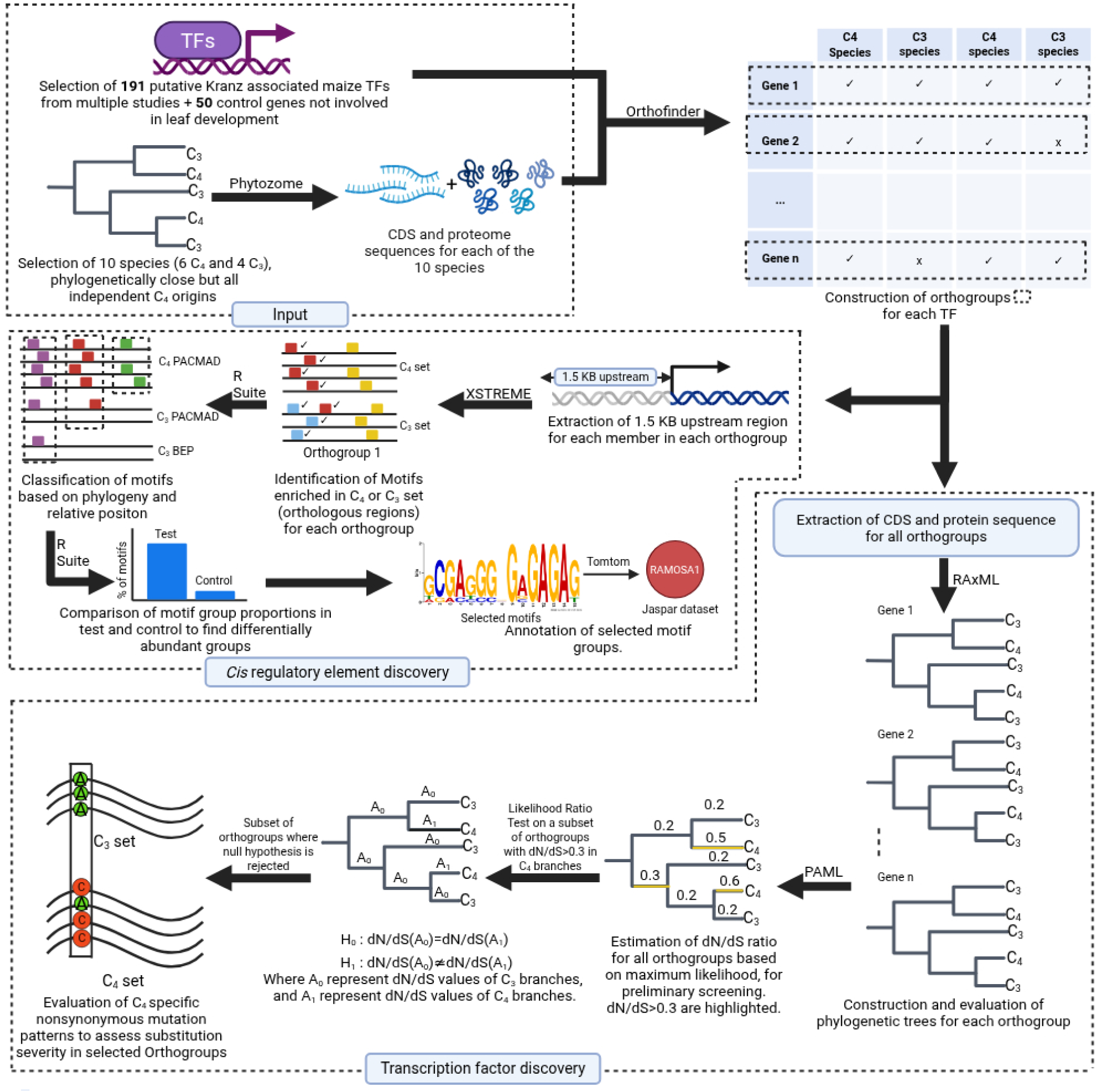
Flowchart explaining the workflow for the discovery of key candidate regulators and *Cis-*regulatory elements acquired for the establishment of Kranz anatomy. Created in BioRender.

### Candidate regulatory genes were largely conserved across six independently originated C_4_ grass species and four closely related C_3_ relatives

The choice of grass species for orthologs was narrowed down to six independently originated C_4_ plants (*Zea mays, Paspalum vaginatum, Panicum halli, Urochloa fusca, Setaria viridis, Eleusine coracana*) (Christin et. al., 2009; Grass Phylogeny Working Group II. 2012) and four C_3_ plants which were closely related to the former set (*Dichanthelium oligosanthes, Chasmanthium laxum, Oryza sativa, Brachypodium distachyon*). While all the C_4_ species belong to the PACMAD clade, the two C_3_ plants, *Chasmanthium laxum* and *Dichanthelium oligosanthes*, also come from the PACMAD clade, thus eliminating clade-specific bias. The remaining two C_3_ species, rice and *Brachypodium distachyon,* belong to the BEP clade. Compared to the previous work of cross-species selection scans (Huang et. al., 2016), this study has six independent C_4_ origins compared to their two, which offers a less biased, more robust view into the evolution of C_4_ components, which can be applied to most of the PACMAD clade C_4_ species. The phylogenetic relationship of these species to each other is shown in Supp. Fig. S1. With the candidate gene set and species finalized, the ortholog clusters of CDS and protein sequences from the selected species were generated for each (putative regulatory) gene selected for the analysis. The Ortholog cluster could be formed for 183 genes, as the others did not have a complete ortholog cluster from 10 species, so we only considered these genes for further analysis. This showed that the candidate regulatory genes selected for the evolutionary analysis are largely conserved across C_3_ and C_4_ grasses.

### Automated construction of gene-trees showed majority of genes of both test-and control-set differing from their species-tree

Since the tool for finding genes subjected to adaptive evolution (namely, PAML) requires the gene tree to be similar to the species tree, 26 ortholog clusters were initially removed as these orthogroups had a too complex a gene-tree to be fixed using partially constrained guide trees (Supp. table S4, Supp. Fig. S2, link in GitHub repository). The dropped cases had unusually high gene duplication, mostly in C_4_ species compared to C_3_ species, except in one case where there was a very large gene duplication in *Chasmanthum* species. Out of these 157 genes, only twenty genes had gene-trees that resembled their species phylogeny (link in GitHub repository). This is an expected pitfall of automated cross-species selection scan as C_4_-related genes share convergent amino acid changes across independent lineages, resulting in gene trees differing from the species tree (Casola et al., 2022). This might result in faulty calculations of adaptive evolution in the ancestral sites in the phylogenetic tree, hence the approach of providing partially constrained guide trees to avoid this pitfall. In the remaining cases, we salvaged the species phylogeny by providing partially constrained guide trees (Supp. Fig. S4, link in GitHub repository). The same procedure with the control gene set resulted in 8 trees that did not qualify for further analysis due to complexity (Supp. table S4, Supp. Fig. S3), and in the rest of the gene-trees, majority (N=33) required partially constrained guide trees to sync with the species phylogeny, showing that even in genes with no known association with kranz development, their gene-trees can differ from the species phylogeny.

### Ten orthogroups from the candidate gene-set and none from the control set showed positive selection pressure selectively on C_4_ orthologs

157 ortholog clusters were examined to find instances of positive selection across the C**_4_** orthologs, wherein the ratio of nonsynonymous substitution rate to synonymous substitution rate (dN/dS or omega) for all branches of a gene-tree was estimated using the maximum likelihood approach. We first identified 37 orthogroups after filtering out the candidates using the following criterion: modest dN/dS ratios (>0.3) in at least 2 C_4_ orthologs belonging to different species. To identify orthogroups where C_4_ species showed higher selection pressure, likelihood ratio tests (LRT) were performed on this subset of orthogroups, comparing models with and without C_4_-specific rate shifts. We observed 10 orthogroups that showed a significant likelihood ratio test statistic (2ΔlnL), involving 13 maize candidate genes (Fig. 2 & Table 1). Out of the ten orthogroups, only four had orthologs from all four C_3_ species, making them higher confidence targets compared to the other six (Table 1). To test the robustness of positive selection results, the same analysis was performed on the control gene set (N=33). While 17 orthogroups from control set had dN/dS (>0.3) ratios in at least 2 C_4_ orthologs belonging to different species, but no orthogroup was found to have a significant likelihood ratio test p-value, ruling out the possibility of chance occurrence of significant cases in the test-set (Supp. Table S5).

**Fig. 2:**
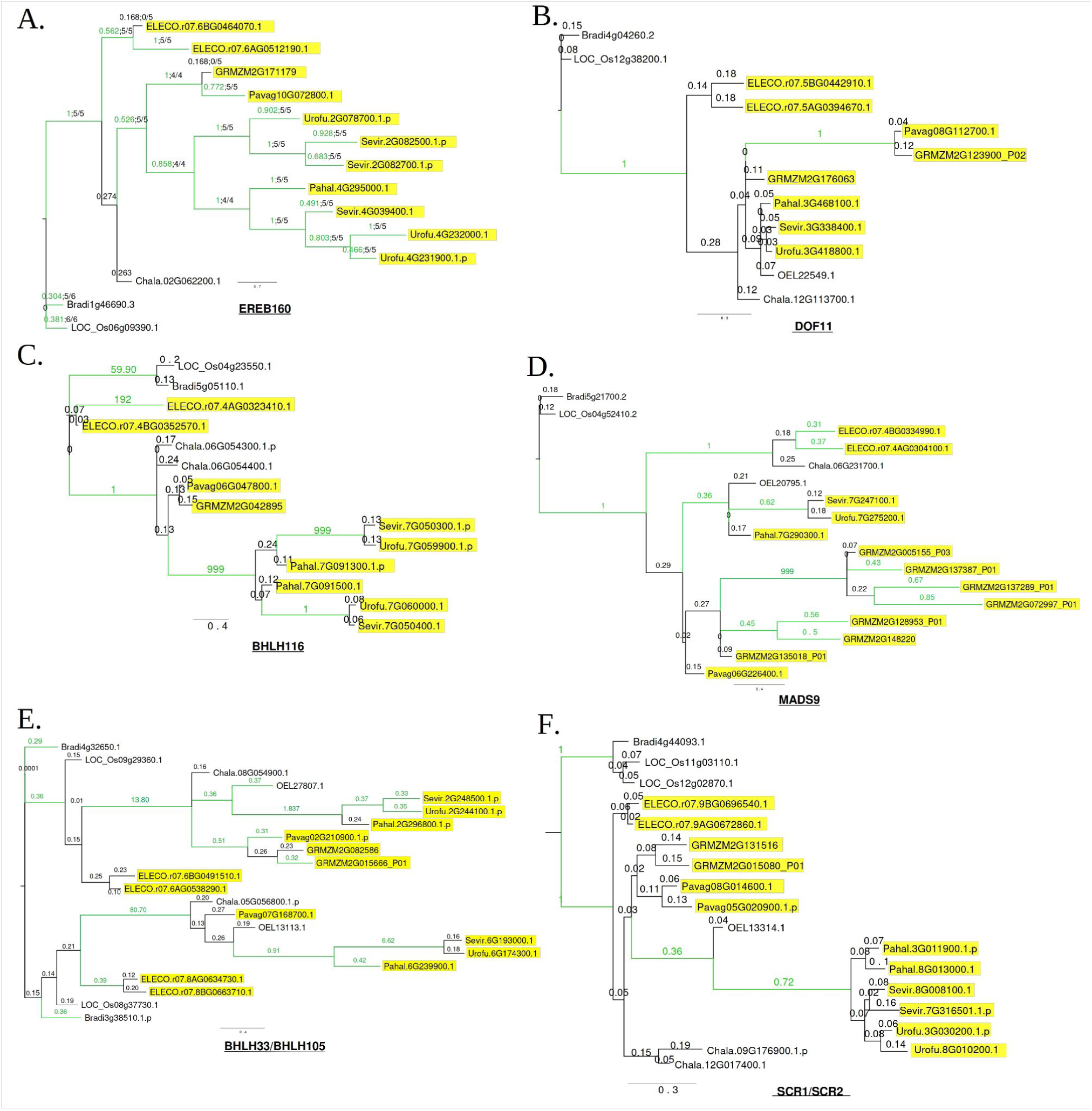
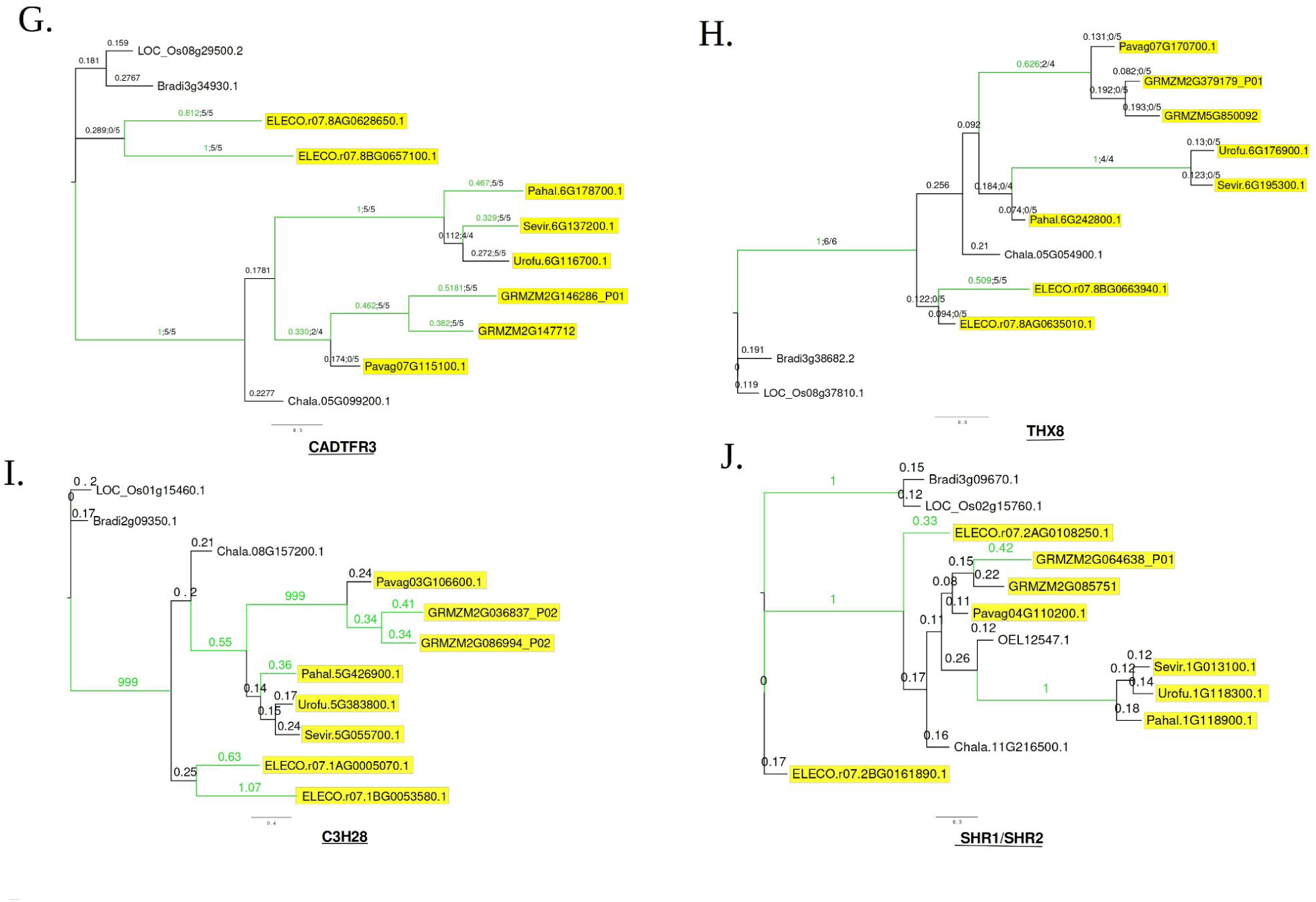
Phylogenetic trees of the orthogroups passing LRT test, with dN/dS values (displayed above the branches) for ten orthogroups: *EREB160* (A), *DOF11* (B), *bHLH116* (C), *MADS9* (D), *bHLH33/bHLH105* (E), *SCRO1/SCR2* (F), *CADTFR3* (G), *THX8* (H), *C3H28* (I), *SHR1/SHR2* (J). Genes with prefixes ‘Bradi’ indicate *Brachypodium distachyon, ‘*LOC’ indicates *Oryza sativa, ‘*Chala’ indicates *Chasmanthium laxum,* ‘ELECO’ indicates *Eleusine coracana*, *‘*OEL’ indicates *Dichanthelium oligosanthes*, *‘*Pavag’ indicates *Paspalum vaginatum, ‘*GRMZM’ indicates *Zea mays, ‘*Pahal’ indicates *Panicum halli, ‘*Urofu’ indicates *Urochloa fusca,* and *‘*Sevir’ indicates *Setaria viridis.* The branches with relatively higher dN/dS (>0.3) values are highlighted in green. The C_4_ orthologs are highlighted in yellow. Bootstrap values are not being shown as after partial constraint tree are provided, the bootstrap values for all species are returned as 90-100.

**Table 1:**
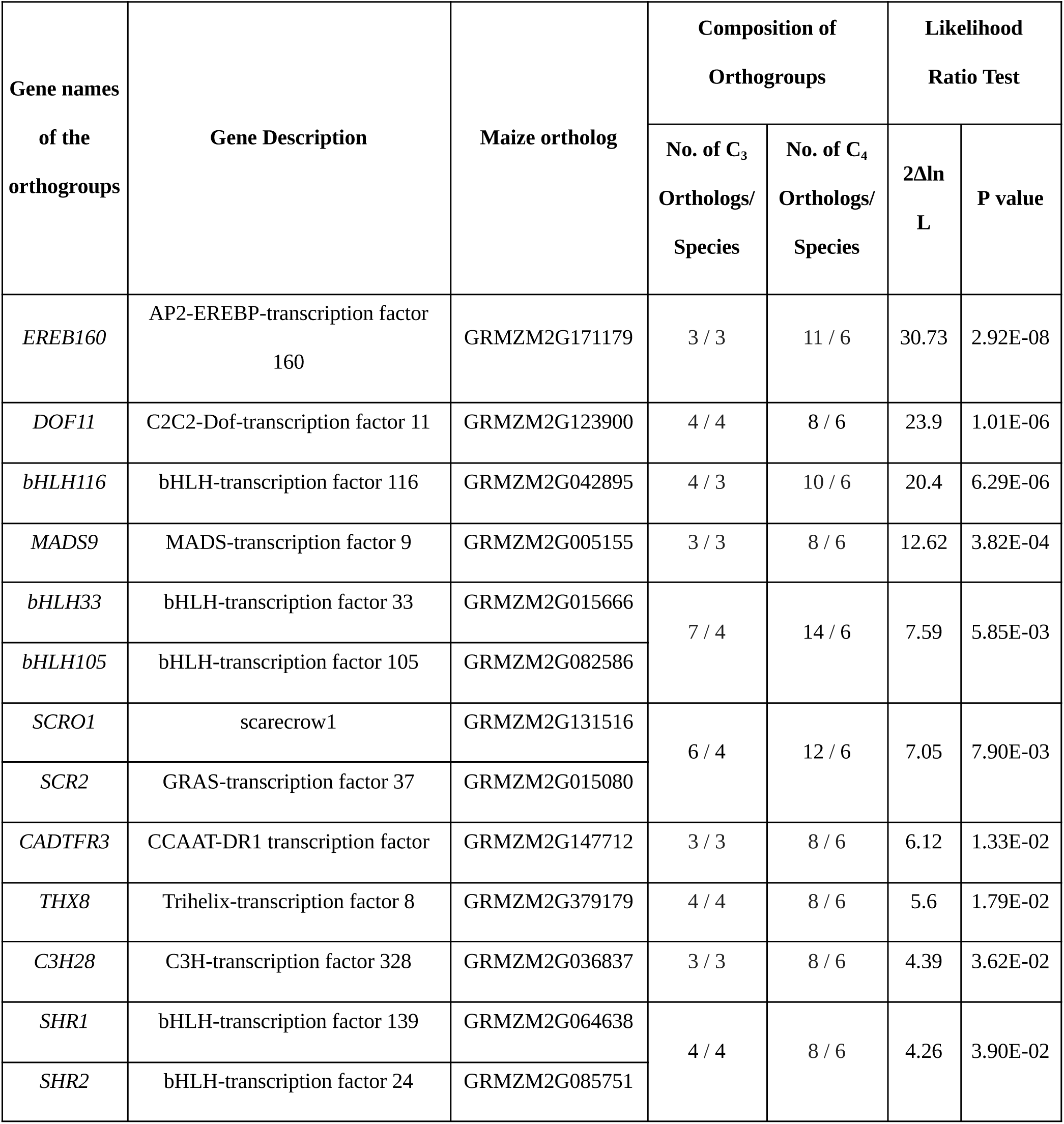
Orthogroups that passed the Likelihood Ratio Test (LRT) for C_4_ branches having higher dN/dS ratios than the C_3_ branches. The test statistic (2ΔlnL) denotes twice the difference in log-likelihoods between alternative and null codon models and is used to evaluate model fit in likelihood ratio tests.

Out of these 10 orthogroups, three have been directly or indirectly implicated in Kranz regulation-*SCARECROW*, *SHORTROOT,* and *DOF11*. The *bHLH33/bHLH105* genes, which were common to both input lists of transcription factors, were earlier overexpressed in rice, but they did not yield any alteration in leaf architecture (Wang et al., 2018), suggesting they might have duplicates in the genome that can take over their function, making a double or triple mutant necessary for observing the changes in other plants. In addition, *EREB160*, while not annotated very well, has also been nominated as a candidate gene in a previous study (Huang et al., 2018), making it an interesting candidate regulator.

The remaining ones are poorly characterized, and so far have shown association with stress and environmental response pathways in *Arabidopsis* and other orthologs (Supp. Table S6): *bHLH33* linked to stress and senescence responses in apples and sweet potatoes (Yu et al, 2021; Yu et al, 2022); *bHLH105* associated with iron homeostasis and manganese stress in *Arabidopsis* (Gao et al, 2020); *EREB160* implicated in drought and salinity responses in *Arabidopsis* (Liu et al, 2020); *C3H28* showed stress-responsive expression patterns (Wang et al, 2025); *bHLH116* described downstream of NF-Y in stress contexts (Wang et al, 2014); and *MADS9* associated with juvenile phase transitions and developmental timing (Beydler et al, 2016).

Alternatively, their functional activities can also be gauged based on any signature expression in a number of transcriptome data, such as “maize gene atlas” (Sekhon et al., 2013), “atlas of the vegetative maize shoot apex” (Knauer et al., 2019), “maize embryonic leaf development” (Liu at al., 2022), “maize early leaf and husk development” (Wang et al., 2013), “comparative leaf developmental gradient of rice and maize” (Wang et al., 2014), and “rice leaf photosynthetic development” (van-Campen et al. 2016). Some of these transcription factors showed distinct expression biases or signature expression during early leaf development (Supp. Fig. S5-S13). The *bHLH33* and *bHLH105* not only showed SAM/leaf-promidia specific expression (Supp. Fig. S5 C&D; Supp. Fig. S6 C&D), but their expression increased from P0 to P3 stage and declined later (Supp. Fig. S14 A&B, middle panel). Moreover, their expression was also higher in the “median ground meristem” (mGM) and the “three–contiguous cell stage” (3C) that initiates vein development, and declined in further stages (Supp. Fig. S5 A&B; Supp. Fig. S6 A&B). On the other hand, *bHLH116*, predicted as a negative regulator of Kranz development (Wang et al., 2013), also showed a similar signature expression as the other two bHLH proteins (Supp. Fig. S12 C&D), but exhibited negligible expression in mGM and 3C cells (Supp. Fig. S12 A&B). The remaining TFs largely showed non-specific expression profiles (Supp. Fig. S5-S14).

### Amino-acid substitutions in these C_4_ orthologs were largely dispersed within the protein sequences

To understand whether the non-synonymous substitutions in the above orthogroups (showing positive selection pressure in C_4_ orthologs) are localised, or domain specific, or spread across the gene, we examined the non-synonymous substitution frequency in the C_4_ subset of the orthogroups selected above. Besides frequency, we examined if the substitution-scores of the substituted amino acids indicate conservation or divergence of their physicochemical properties with respect to the ancestral residue (see methods). The substitutions were not uniformly distributed across the genes, but had distinct gene-specific patterns (Fig 3). Although very few substitutions were present with frequency in C_4_ orthologs ≥0.75 (one in *EREB160* and another in *bHLH116*), there were many cases of mean substitution scores in negative (<-0.5), showing substitutions very different in physicochemical properties from their C_3_ counterparts (Fig 3).

**Fig. 3:**
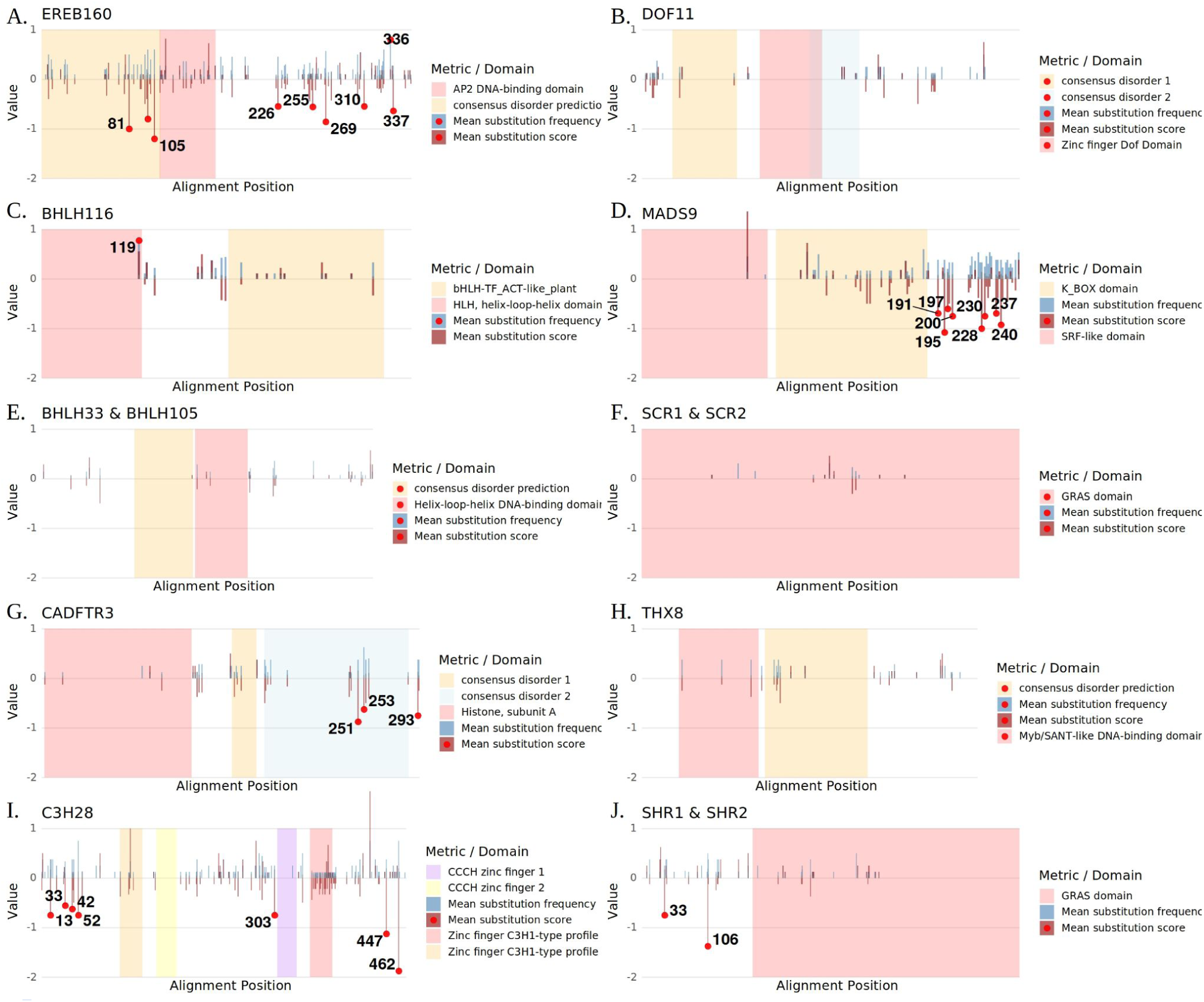
Mean frequency of non-synonymous substitutions in the C_4_ orthologs, and their mean substitution scores for the selected orthogroups. The multiple alignment positions are plotted along the X-axis, and both substitution frequency (blue lines) and substitution scores (red lines) are plotted along the Y-axis. The substitution frequency >0.75 and mean substitution score <-0.5 are highlighted with dots.

Unlike *EREB160*, in the case of *bHLH116*, the position having a substitution frequency >0.75 in C_4_ orthologs was in a conserved domain, i.e., a helix-loop-helix domain. On examining the substitution trend in an expanded orthogroup involving orthologs from the entire *Poeceae* family, the alignment showed substitution of Methionine/Leucine (ancestral residues) to other hydrophobic amino acids with smaller side-chains (Valine/Isoleucine/Alanine); the substitutions were largely C_4_-specific rather than being clade-specific (Supp. Fig. S13).

On the other hand, five orthogroups (*EREB160, MADS9, CADFTR3, C2H28* and *SHR1/2*) had substitutions, often in clusters, involving change in the physico-chemical properties of the amino acids (mean score <-0.5) (Fig. 4). Such substitutions were observed either in the intrinsically disordered regions or in unknown domains, and their frequency in C_4_ orthologs was lower than the chosen cut-off of <0.75, indicating random or unfixed substitutions. The remaining four orthogroups did not exhibit any of the above-mentioned substitution trends, namely *DOF11, bHLH33/115, SCR1/2*, and *THX8* (Fig. 3).

**Fig. 4.**
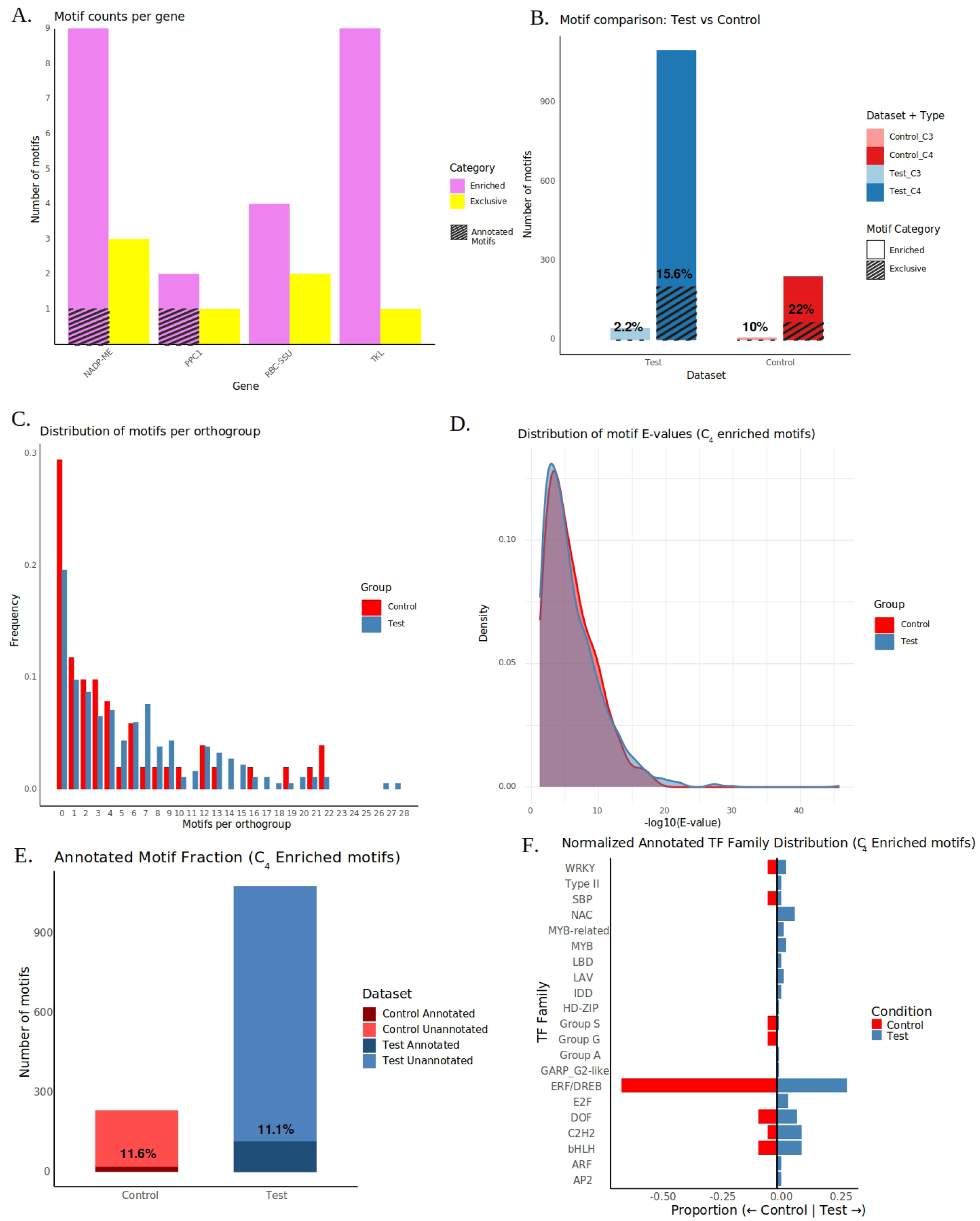
Motif enrichment and distribution in C_4_-associated and control gene sets. (A) Number of enriched and C_4_-exclusive motifs per representative gene. (B) Summary of motif enrichment across datasets, showing higher enrichment and a greater proportion of exclusive motifs in the C_4_ (test) set relative to the control. (C) Distribution of motif counts per orthogroup in test and control datasets. (D) Density distribution of motif E-values (−log10) in test and control datasets. (E) Annotation of C_4_ enriched motifs from both test and control sets. (F) Comparison of proportion of annotated motif groups (i.e., the corresponding TF family that binds to them) between test and control sets.

### Limited adaptive changes in the coding sequences prompted exploration of changes in its regulatory sequences

After exploring changes in the coding sequences of 182 genes and identifying only a limited number of cases showing species-wide variation, we turned to alternative regulatory mechanisms that could explain differential activity in C_4_ plants. To identify common transcription factor binding sites that may drive C_4_-specific expression patterns, we focused only on the upstream regulatory regions of these orthologs, and the gene body was excluded given the high sequence conservation (which might yield too many false positives). By examining motif composition and enrichment within proximal promoter regions, we sought to uncover parallel evolutionary changes in *cis*-regulatory elements that could modulate gene expression without requiring alterations to the TF coding sequences themselves. In doing so, we considered two complementary approaches: identifying motifs that are consistently enriched in C_4_ orthologs relative to their C_3_ counterparts, and those that are exclusively present in C_4_ orthologs while being absent in C_3_ counterparts.

### Previously identified motifs driving C_4_-specific expression were found enriched in the upstream of C_4_ orthologs

We first verified the two approaches on a few known motifs driving C_4_ specific expression in a few C_4_ biochemical enzymes, such as *PPC1, RBC-SSU1,* and *NADP-ME* (Gupta et al., 2020; Burgess et al., 2019; Xu et al., 2001; Giuliano et al., 1988). We could partly recover the previously identified motifs only from enriched motifs, with none of them coming from the exclusive set of motifs (Fig. 4A; Supp. Table S7). Out of five Conserved Nucleotide Sequences (CNS) previously characterized for PPC1 gene we found only one CNS region, and missed the TATA box and the other 3 CNS regions. With transketolase, we did not find the DNA Hybridization Sites (DHS) that were present not just in the promoter but in the introns and UTR regions of the genes (Burgess et al., 2019). These results indicated that the enrichment approach has better potential than the exclusive approach in predicting the C_4_ specific motifs from the upstream sequences.

### A large number of motifs, enriched in the C_4_ set, were likely due to ancestry

For the 184 orthogroups generated from the test-set, 1.5 kb upstream of the transcription start site (TSS) of each gene was extracted from the genomes. We obtained 1,101 motifs enriched in C_4_ grasses across 149 orthogroups, and only 46 motifs enriched in C_3_ grasses across 26 orthogroups (Figure 4B). Out of these motifs, 203 were exclusive to C_4_ species (Fig 5B). We repeated this protocol with 51 control genes, and we could identify 315 motifs across 37 orthogroups enriched across all C_4_ grasses, and 10 motifs from 6 orthogroups enriched in C_3_ grasses (Fig 4B). To examine the difference in the motif occurrence between test-and control-set, we compared the distribution of motifs per orthogroup for control and test set, which seemed to have small differences (Fig. 4C), indicating no uniqueness in distribution of C_4_ enriched motifs. To confirm if the observations are not due to imbalance in size of the test and control set, we then randomly sampled 55 orthogroups from the test set ten times, and then compared the motif distribution to the control set which again showed negligible difference in the distribution (Supp. Fig. S15). We also checked the E-value distribution of motifs in Test and control sets (Fig. 4D), and in all of these cases, there were no differences in the motif distribution between Test and Control sets.

**Fig. 5:**
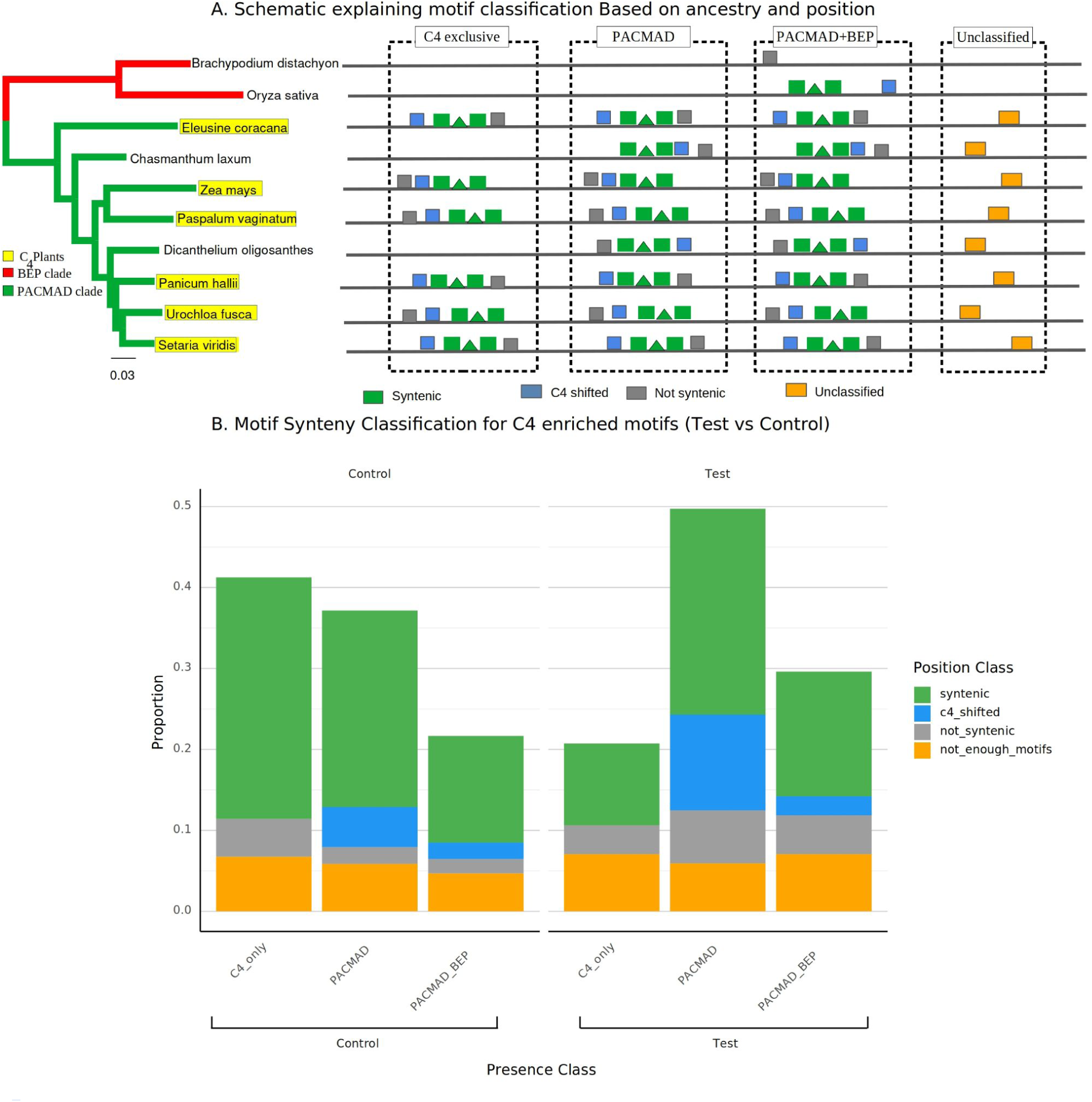
A. Schematic diagram explaining motif classification based on ancestry and relative position. B. Proportion of different motif classes in C_4_-enriched motifs of Test and Control set.

If these motifs are indeed not actual TF binding sites but rather relics of ancestry, we hypothesized that the real motifs that are not artifacts would be annotatable to the known motifs. To test this hypothesis, we annotated all the motifs present in control and test sets (see Methods). Although a large number of motifs were found to be enriched, only a small portion of these motifs were annotated in both test and control sets (Fig. 4E; Supp. Table S8). This pattern is consistent across both sets of orthogroups, including those with very high motif counts, indicating that having a lot of motifs does not mean these are annotated ones (Supp. Fig. S16). Notably, both datasets exhibit a similar distribution for annotated motifs as well, but instead of the trend where a few orthogroups harbor a large number of motifs while the majority contain relatively few, annotated motifs seem to be present in similar numbers across all (1-5 in test and 1-3 in control).

While motif distributions remain largely consistent between the two datasets, aggregating annotated motifs at the level of transcription factor (TF) families revealed distinct regulatory signatures (Fig. 4F). As demonstrated in the TF family enrichment test, the test dataset exhibits a significantly larger number of members across multiple families, suggesting a shift in regulatory composition. Notably, the *E2F, MYB, MYB-related, C2H2, bHLH, NAC, IDD, ARF, AP2, WRKY,* and *DOF* families have more members in the test set, with the *E2F* and *MYB* families demonstrating the highest disparity in numbers between control and test-sets. Given the established roles of these families in transcriptional regulation, developmental control, and metabolic coordination, their quantitative enrichment suggests a regulatory reprogramming in the test condition. Conversely, a restricted number of TF families—including GRF and Group G—exhibit a relative enrichment within the Control dataset. When their enrichment was tested using Fisher’s exact test, only the *ERF/DREB* family had a statistically significant enrichment (p-value < 0.05) in the control compared to the Test-set. These (motif-binding) transcription factor families, that are absent or having lower proportion in the control set, are most likely associated with the leaf development regulation, as the control set have genes that do not show any specific expression profile during early leaf development, while the test-set is enriched with transcription factors that are expressed during early leaf development. But we do not know whether these motifs are related to Kranz anatomy development.

### Separating motifs into syntenic and non-syntenic helped to differentiate test and control motifs

Since annotation had a limited role in distinguishing motifs appearing due to ancestry from other motifs, we used a phylogeny-aware synteny framework that considered both lineage/ancestry and relative position. Since multiple studies have hypothesized that rather than acquiring novel motifs, C_4_ plants co-opted motifs from pre existing C_3_ modules independently ((Williams et al., 2016; Reyna-Llorens et al., 2018; Swift et al., 2024), we hypothesized that co-opted C_4_ regulatory motifs would not have a deeply conserved positional architecture across all the plants, especially that these motifs would not be syntenic (a conserved position across orthogroups) across all the C_4_ plants and their C_3_ relatives; as synteny indicates these motifs were acquired through vertical descent. First the motifs were separated into three classes based on shared ancestry (C_4_-only, PACMAD, and PACMAD+BEP), then the relative position within each class was deduced (syntenic, C_4_ shifted, and non-syntenic) (Fig. 5A; see Methods). Fig. 5B shows the results of classification of motifs in test and control sets.

We observed that the proportion of C_4_-only or exclusive motifs was high in the Control, but the lower in the Test set. This might be partly due to imbalance in the group-size of C_3_ species (N=4) or their orthologs compared to C_4_ counterparts; any further drop in sample size of C_3_ sequences can inflate the C_4_ exclusive motifs. Nevertheless, this indicates that C_4_-only motifs are more likely to be observed by chance than others.

There was however an increase in the proportion of other two classes (PACMAD and PACMAD+BEP) in the test set. Within both classes, the proportion of not-syntenic and C_4_-shifted motifs was much higher in the test compared to the control set. Based on this difference, we further focused on these non-syntenic/C_4_-shifted motifs. In total, we identified 336 non-syntenic and C_4_-shifted motifs in the Test coming from 66 genes, and 39 non-syntenic and C_4_-shifted motifs coming from 9 genes in the control set (Supp. Table S9). Out of the genes from test set, 28 genes belonged to study 1 and 38 genes belonged to study 2. Genes with same *cis*-motifs can show differential expression in C_4_ and C_3_ tissues in the same plant, which may be due to the tissue specific *trans* factors present, or other epigenetic changes in the two tissues. When we annotated them, we could only find annotations for 9 non-syntenic and 30 C_4_-shifted motifs in the test set, and 2 non-syntenic and 2 C_4_ shifted motifs in the control set (Fig 6).

**Fig. 6:**
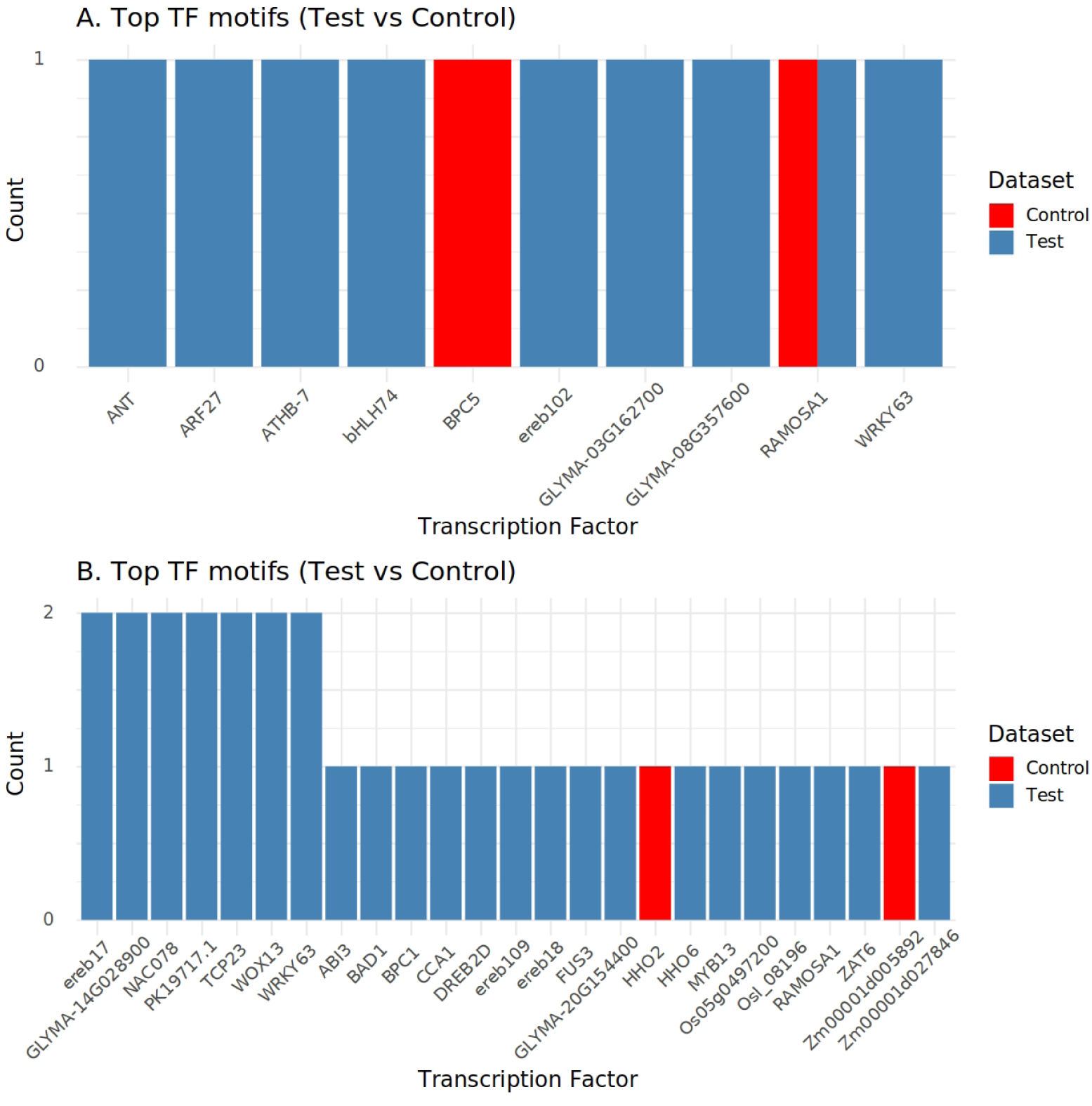
Motif annotation across Test and Control set for the annotated non-syntenic(A) and C_4_-shifted motifs (B)

Since these motifs seemed to have some common binding Transcription factors and the fact that some genes share multiple of these motifs, we created a directed network of Transcription factors binding to these motifs and the genes that have these motifs in their upstream (Fig. 7).

**Fig. 7:**
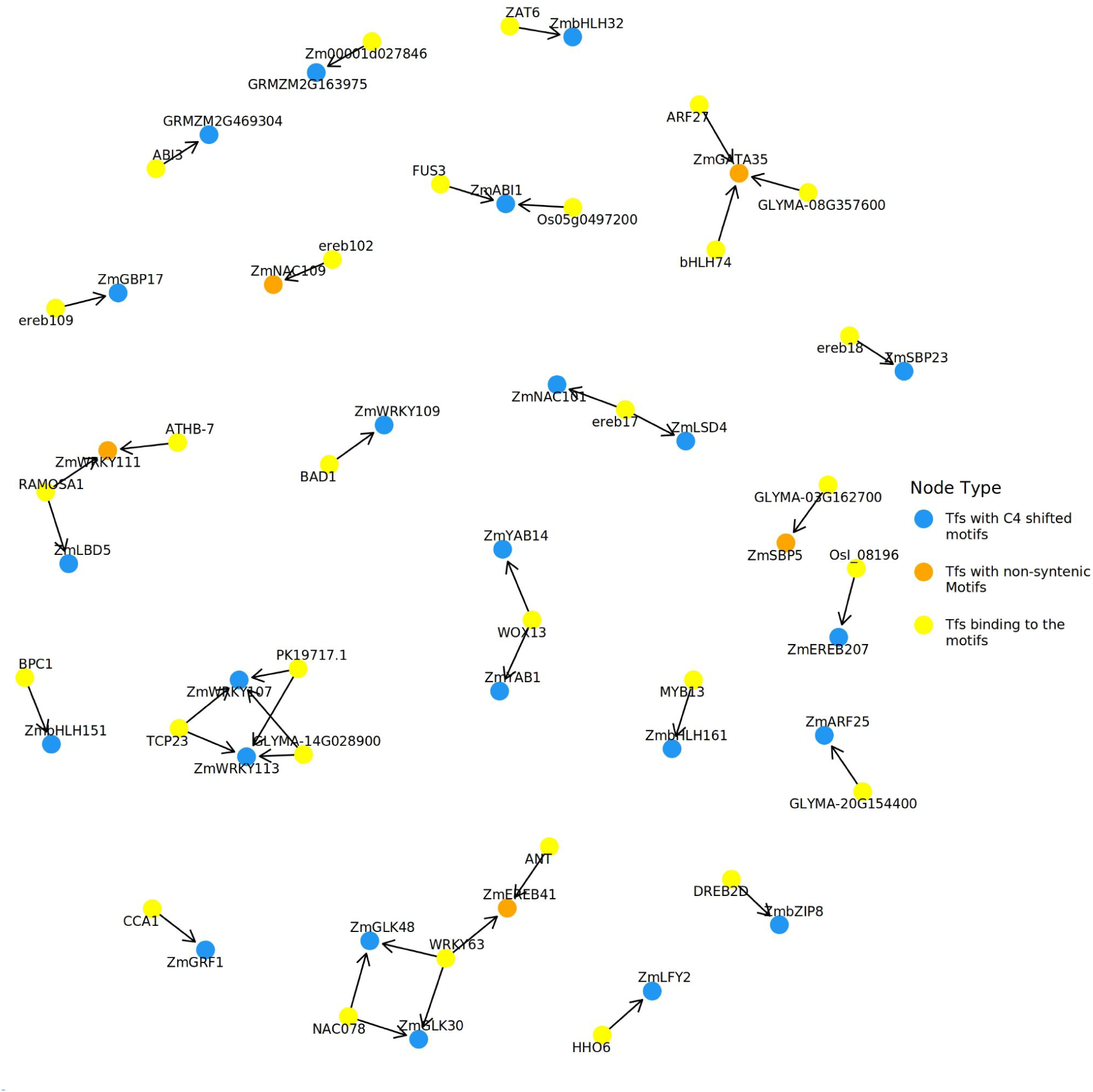
Predicted interaction of genes annotated with C_4_-shifted and Not-syntenic motifs with TFs that bind to them.

We see some expected modules, like how both *GLK* genes are targeted by both *WRKY63* and *NAC078,* or the motifs of *WRKY107* and *WRKY113* have the three motifs that bind to the same 3 transcription factors. There were also interesting pairs like *NAC101* and *LSD4,* which share a motif that can bind to *ABI*, or the *WRKY63* motif shared among 2 *GLK*s and *EREB41*. While only a small portion of the non-syntenic and C_4_-shifted motifs were annotated, the rest of the motifs remain as possible novel or putative regulators for C_4_ regulation.

## Discussion

### Limited Adaptive Coding Evolution observed in the putative Kranz Development Associated Transcription Factors

Coding convergence in core C_4_ metabolic enzymes is well established (Davidson, 2006; Gupta et al., 2020). Enzymes such as *PEPC, PPDK, NADP-ME,* and *CA* originated from gene duplication events, including whole-genome and tandem duplication (Wang et al., 2009), and show elevated dN/dS ratios and accelerated amino acid substitutions along C_4_ branches (Christin et al., 2007, 2009a, 2009b, 2013). Importantly, recruitment of specific isoforms was not random but strongly predicted by high ancestral leaf expression levels in non-C_4_ grasses (Moreno-Villena et al., 2017). Once recruited, these isoforms underwent further regulatory and coding evolution, resulting in increased expression and adaptive amino acid changes.

Similar to C_4_ metabolic enzymes, transcription factor families in C_4_ species show extensive gene duplication relative to their C_3_ counterparts (Tkach et al., 2025), consistent with the broader duplication history described for grass genomes (Wang et al., 2009). This often resulted in uneven ortholog representation, with C_4_ species possessing twice as many copies, or more, than corresponding C_3_ orthologs. Despite this expanded gene space, very few transcription factors showed significant evidence of adaptive coding evolution. From the initial set of 191 candidate transcription factors, only 13 transcription factors showed significantly elevated dN/dS among C_4_ orthologs. This low proportion indicates that C_4_-specific transcription factor reprogramming *infrequently* involved direct changes in the amino acid sequences. The low number can also be due to the constraint posed by search for parallel evolutionary changes, which may at times exclude all instances of gene duplications.

A genome-wide scan across 6,784 orthogroups, involving two independent C_4_ origins, identified only six transcription factors with significant adaptive signatures (Huang et al., 2017), with only one gene common between the two studies (namely, *EREB160*). The five transcription factors, specific to Hunag et al. (2017), were not examined by the present study as their expression patterns showed no or little association with the Kranz development. For the seven transcription factors specific to the present study, they were possibly not examined by Huang et al. (2017) as their study disqualified a lot of orthogroups, something we tried to remedy by providing guide trees for phylogeny construction, helping us preserve a larger set of genes for downstream analysis. Together, these results suggest that transcriptional reprogramming during Kranz evolution likely relied more on regulatory or expression-level modifications rather than widespread protein-coding divergence.

### Amino-acid Substitution Patterns Indicate Convergent Changes in Intrinsically Disordered Regions (IDRs) of candidate Transcription Factors

Investigations into adaptive changes reported a few candidate transcription factors showing distinct substitution clusters, namely *EREB160, CADTFR3, MADS9,* and *C3H28*. Such clusters did not overlap with domains annotated by InterPro. The absence of substitutions within canonical DNA-binding domains suggests that adaptive changes may have occurred in regions affecting protein–protein interactions or regulatory interfaces rather than core DNA-binding activity. This pattern is consistent with fine-tuning of regulatory function (Song and Akey, 2026) rather than complete functional rewiring, and contrasts with the more pronounced coding divergence observed in C_4_ metabolic enzymes (Wang et al., 2009; Huang et al., 2017). Of the two cases showing a strong C_4_-specific substitution pattern, *bHLH116* had it within a conserved domain (helix region). Functionally, the position happens to be the last conserved base of helix-2 of the *bHLH* domain in plants (Thoben and Pucker, 2023; Carretero-Paulet et al., 2010), and is involved in the formation of the hydrophobic core during dimerization (Ferre-D’Amare et al., 1993). Substitution from larger hydrophobic residues to smaller ones can impact the stability of interacting monomers, and it may result in modified regulation of the putative target genes of *bHLH116*.

Collectively, these findings indicate that adaptive coding evolution of candidate transcription factors during C_4_ evolution was limited and restricted to a smaller subset. These genes share common features: membership in expanded transcription factor families, broad expression across tissues, stress-associated functional backgrounds, and activity during early leaf development. The clustering of substitutions outside annotated domains further suggests that evolutionary changes may have refined regulatory interactions rather than redefined core protein functions. Together, this supports a model in which C_4_ anatomical evolution involved selective tuning of stress-integrated regulatory hubs and expression-level reprogramming, rather than widespread innovation in transcription factor coding sequences.

Given the limited evidence for adaptive coding evolution among putative Kranz-associated transcription factors, our results suggest that the emergence of C_4_-specific evolutionary changes was primarily driven by changes at the regulatory level rather than through widespread protein sequence divergence.

### Candidate motifs driving C_4_ anatomy are more likely to be co-opted from pre-existing C_3_ motifs, rather than C_4_-exclusive novel motifs

Our results help address a key gap in understanding the regulatory basis of C_4_ photosynthesis by identifying candidate *cis*-regulatory motifs that may have been recruited during the evolution of Kranz anatomy. While previous studies have pointed to specific regulatory elements in individual genes, a broader, comparative framework across orthologs has been lacking. By systematically identifying motifs enriched in C_4_-associated genes, this study provides a set of candidate regulatory elements that can serve as a foundation for future functional validation. In this way, our work contributes to building a more comprehensive picture of how gene expression patterns underlying C_4_ traits may be encoded at the regulatory level.

Interestingly, the observation that the control dataset contains a higher proportion of C_4_-exclusive motifs challenges the idea that C_4_ evolution is driven by the widespread emergence of novel, lineage-specific regulatory elements. If C_4_-specific expression relied heavily on newly evolved motifs, we would expect these to be enriched in the test dataset rather than the control. Instead, this pattern supports the co-option hypothesis, suggesting that C_4_ evolution primarily involved the reuse and repurposing of pre-existing regulatory elements rather than de novo motif acquisition. This aligns with the broader view that evolutionary innovation often arises through modification of existing regulatory frameworks (Williams et al., 2016; Reyna-Llorens et al., 2018; Swift et al., 2024).

Further support for this model comes from the enrichment of non-syntenic and C_4_-shifted motifs in the test dataset. These motifs, which lack conserved positional architecture across species, point toward a mechanism of regulatory rewiring rather than strict conservation. The absence of strong syntenic constraints suggests that functionally relevant motifs may have been repositioned or independently recruited into new regulatory contexts during C_4_ evolution. Such positional flexibility is consistent with the co-option model, where ancestral regulatory elements are reused in different genomic contexts to generate novel expression patterns.

### Candidate genes with adaptive evolution and those with C_4_ enriched non-syntenic candidate motifs do not intersect

An additional pattern that emerges from our analysis is the apparent separation between genes exhibiting adaptive changes in their protein sequences and those harboring non-syntenic, C_4_-enriched regulatory motifs. Notably, there is minimal overlap between these two groups, suggesting that coding sequence evolution and regulatory rewiring may have largely operated on distinct subsets of genes during C4 evolution. This limited intersection implies that the evolution of C_4_ traits may not have relied on simultaneous modifications at both the protein and regulatory levels within the same genes, but rather on complementary mechanisms acting across different components of the regulatory network. In this context, genes undergoing adaptive coding changes may represent key regulatory nodes where subtle functional tuning was sufficient, while a broader set of genes achieved C_4_-specific expression primarily through the acquisition or repositioning of cis-regulatory elements. Such a division of evolutionary roles is consistent with a modular model of network evolution, where protein-level changes refine specific regulatory interactions, while cis-regulatory changes drive large-scale transcriptional reprogramming (Schmitz et al, 2022; Wagner et al, 2008). However, it is also important to consider that this apparent separation could be influenced by limitations in detection power for both coding selection and motif annotation, and therefore warrants further validation. Nonetheless, this pattern reinforces the idea that regulatory evolution, rather than coordinated coding and regulatory changes within the same loci, was the dominant force shaping the emergence of C_4_-specific gene expression programs.

### Limitations and challenges associated with identifying regulatory components of C_4_ evolution

While trying to identify regulatory genes and *cis*-elements associated with Kranz anatomy we did note a few limitations. One major limiting factor is the availability of a limited number of well annotated C_4_ and C_3_ genomes from PACMAD clade, because of which our sample size for the species for the analysis remained small. A larger sample size with more species with independent origins of C_4_ and more C_3_ from PACMAD clade would result in a much more unbiased analysis and provide more robustness to the results. At the same time, our analysis highlights an inherent challenge in studying changes in regulatory sequences during adaptive evolution through comparative approaches. Motif discovery across multiple species, particularly under a parallel evolutionary framework, is biased toward conserved sequences and shared ancestry, making it difficult to distinguish putative regulatory elements from background signals. This limitation is evident in the large number of enriched motifs that likely reflect ancestral sequence conservation rather than regulatory function. By incorporating a phylogeny-aware synteny framework, we attempted to mitigate this bias and refine candidate selection. Nevertheless, these challenges underscore the need for integrative approaches combining computational predictions with experimental validation to fully resolve the regulatory mechanisms underlying C_4_ evolution. Additionally, our analysis is limited in its ability to detect adaptive changes at the gene family level. The ortholog clusters primarily consist of closely related sequences, limiting the detection of more distantly related homologs that may have independently evolved in different C_4_ lineages. However, including such divergent family members can lead to poorly resolved phylogenetic relationships, indicating that more sophisticated approaches will be required to address this limitation.

## Materials and Methods

The workflow followed for the current study is represented as a flowchart in Figure 1.

### Input geneset of putative regulators of Kranz anatomy

Candidate genes associated with Kranz anatomy were compiled from maize transcriptomic studies of early leaf development and cell-type specification (Wang et al., 2013; Liu et al., 2022), and were compared with a cell-specific C₄ transcriptomic study in switchgrass (Rao et al., 2016). Candidate genes were selected based on their reported differential expression during early leaf development in foliar vs husk primordia, and/or preferential expression in bundle sheath or mesophyll precursor cells. Specifically, Wang et al. (2013) identified developmental regulators based on foliar versus husk leaf expression profiles in maize, while Liu et al. (2022) resolved cell-type–specific regulatory programs using laser capture microdissection of embryonic leaf tissues of maize. Due to substantial overlap with the maize-derived candidates and the maize-focused scope of this study, only genes from Wang et al. (2013) and Liu et al. (2022) were retained for downstream analyses, along with 51 control genes picked from Wang et al. (2013) showing neutral or no differential expression profile during Leaf development. (Supplemental Table S3).

### Selection of species for obtaining orthologs

To find evidence of positive selection and discovery of *cis* elements exclusive to C_4_-specific orthologs, we selected six C_4_ plants having independent origins (*Zea mays, Paspalum vaginatum, Urochloa fusca, Panicum halli, Setaria viridis, Eleusine coracana*) (Christin P.A., et. al., 2009, Grass Phylogeny Working Group II. 2012) and four closely related C_3_ plants (*Dichanthelium oligosanthes, Chasmanthium laxum, Oryza sativa, Brachypodium distachyon*) for the analysis. These plants were highly scrutinised to make sure there are independent origins between all of the C_4_ species, ending with 6 independent origins. Furthermore, we only picked plants with a complete coding sequence (CDS) and and Protein sequences available. These grasses belong to two closely related clades (BEP and PACMAD) within the *Poaceae* family. The PACMAD clade included all C_4_ species, with two C_3_ plants, namely, *Dichanthelium oligosanthes* and *Chasmanthium laxum,* as controls. Such a selection was made to avoid any biases due to the clades the plants belong to. Further C_3_ plants from more distant clades were not selected to avoid stronger clade biases. The independent origins of selected C_4_ species avoid biases due to a common C_4_ ancestor.

### Ortholog cluster (orthogroup) creation

Reference coding sequences (CDS) and protein sequences of selected plants were retrieved from the Phytozome v14 database (https://phytozome-next.jgi.doe.gov) (Goodstein *et al*., 2012). Dichanthelium v.1 genome assembly and gene model annotations were retrieved from NCBI under accession number LWDX00000000(Studer, *et al*, 2016). Ortholog clusters were constructed using the reference CDS sequences using Orthofinder(version 3.0.1b1) (Emms and Kelly, 2019) with-MCL = 1.5,-T-Raxml-NG,-y-split as custom changes. We managed to construct ortholog trees for 182 genes. For the others, the tree was not complete.

### Phylogenetic tree construction and its quality control

An in-house Perl script (GitHub link provided) was used to extract the sequences of orthogroups to which the selected genes belonged. The orthologs’ protein and nucleotide sequences were aligned using the TranslatorX program (Abascal *et al*., 2010) to ensure that the reading frame remained intact. The multiple-aligned protein sequences were used to generate the best tree along with the bootstrap values (100 iterations of bootstrapping) using Randomized Axelerated Maximum Likelihood-Next generation (RAxML-NG) (version 1.2.2) tool (Kozlov et al., 2019). While generating the phylogenetic trees, if the gene-tree differed from species tree, then partially constrained guide trees were provided to ensure the correct species phylogeny was preserved. This resulted in the bootstrap values for all species to be returned as 90-100. So we ignored showing the bootstrap values. After this step, we evaluated the phylogenetic trees for tree complexity and correct species phylogeny. All the alignments used for tree construction, partially constrained guide trees, and the final tree are uploaded in GitHub.

### Examining Positive selection in ortholog clusters

To investigate evidence of adaptive evolution, Phylogenetic Analysis by Maximum Likelihood (PAML, version 4.9) was used which requires a phylogenetic tree representing species phylogeny as closely as possible (Yang, 2007). The best gene tree, along with the aligned nucleotide sequences, was submitted to the *codeml* program from PAML(version 4.9) (Yang, 2007). The free-ratio model was used for preliminary screening, which estimates a branch-specific dN/dS value across the phylogeny.

A higher ratio of dN/dS (or omega) indicates positive selection, and we selected 0.3 as a cutoff for omega. A stringent omega (>1) risk missing positive selection acting on a subset of orthologs/sites, so a threshold was chosen that should be above the values indicating strong purifying selection. Roth and Liberles (2006) reported that an average omega of 0.21 ± 0.04 showed a strong negative selection on most branches in the gene families of higher plants. Phylogenetic trees of the best tree and omega values for every branch were visualized using Figtree (version 1.4.4) (http://tree.bio.ed.ac.uk/software/figtree/). Once both phylogenetic tree and omega values were available, orthogroups with omega values >0.3 for at least 2 C_4_ orthologs belonging to different species were identified and screened out.

To formally evaluate differences in selective pressure between C_3_ and C_4_ lineages for these genes, likelihood ratio tests (LRTs) were performed on the orthologs which based the preliminary screening step, by comparing the nested codon substitution models implemented in codeml following the protocol provided in Álvarez-Carretero et al. (2023 for the branch model test. Specifically, a null model assuming uniform selective pressure across the branches was compared against an alternative model allowing lineage-specific variation in dN/dS. The test statistic (2ΔlnL) was calculated as twice the difference in log-likelihoods between the two models and assessed for significance using a chi-square distribution with 1 degree of freedom. For significant genes, phylogenetic trees annotated with branch-specific dN/dS values were visualised, and branches with dN/dS > 0.3 were highlighted on the trees using FigTree (version 1.4.4). For control set, none of the genes passed preliminary screening and LRT tested was conducted for all as a control for screening.

### Amino-acid substitution pattern in C_4_ orthologs of selected orthogroups

To determine whether amino-acid substitution in C_4_ orthologs were randomly distributed or domain-specific, we analyzed the substitution frequency and substitution scores in the C_4_ set. We first identified the positions in the multiple sequence alignment where C_4_ orthologues have a substitution, and identified the ancestral and substituted amino acids. We then measured the substitution frequency in C_4_ orthologs and highlighted cases with frequency>0.75. The substitution scores were then obtained by averaging the log-odds score for the corresponding C_4_ substitutions from the BLOSUM62 scoring matrix (Henikoff and Henikoff) using a Python script (see Data availability section). The substitution scores are positive when the amino acid that replaces the original is very similar in chemical properties, and negative when the amino acid is very different. We highlighted the positions where the mean substitution frequency>0.75 or the mean substitution score was <-0.5, indicating a large change in those positions. We finally overlaid these results onto domain information retrieved from InterPro (Blum et al., 2025; accessed in Dec. 2025). The visualization and plotting were done using Rstudio (version 1.2.5033) (R Core Team, 2021). The scripts are provided on GitHub.

Cases with mean substitution frequency >0.75 in C_4_ orthologs, were further verified in an extended orthogroup (at family level, i.e., *Poeceae*) obtained from OrthoDB v12.2 (Tegenfeldt et al., 2024; accessed in March 2026). The orthologs were searched using NCBI gene ID for the gene of interest, and their protein sequences were downloaded and multiple-aligned using MAFFT server (Katoh K., et al, 2019; accessed in March 2026) involving iterative refinement method, E-INS-i, meant for sequences with multiple conserved domains and long gaps. The substitution frequency of the amino acid in question in C4 orthologs was estimated.

### Analysis of functional information and C_4_-specific

Gene ontology and functional details for the genes were retrieved from Monocots Plaza 4.5 (Van Bel *et al*., 2022)(https://bioinformatics.psb.ugent.be/plaza).

### Finding *Cis-*regulatory elements that are enriched in C_4_ species

To identify the cis-regulatory motifs enriched in C_4_ grasses, we considered a 1.5kb upstream sequence from the gene start site, excluding the UTRs, from 191 maize genes (transcription factors) that are documented to play a potential role in Kranz anatomy development (Liu et al., 2022; Wang et al., 2017). For the control set, we also extracted a 1.5kb upstream sequence (excluding UTRs) from 55 orthogroups, which are not documented in the literature to take part in Kranz anatomy development. The 1.5kb upstream sequences were extracted using a Python script from Genome sequences and GFF files of selected plants, which were retrieved from the Phytozome v14 database (https://phytozome-next.jgi.doe.gov). These upstream sequences were divided into C_4_ and C_3_ sets (based on grasses from which they are extracted) for motif discovery. The upstream sequences were combined and run in XSTREME, a tool in the MEME suite (Bailey *et al*., 2015), which looks for enrichedmotifs in a positive set compared to a negative input set using MEME (Bailey and Elkan, 1994) and STREME (Bailey, 2021), which are also part of the MEME suite. The XSTREME output files were inspected, and motifs that are identified by MEME and present across all the 6 C_4_ plants were extracted. Exclusivity of these motifs to C_4_ orthologs were verified by running FIMO (version 5.5.9) (Grant et al, 2011). The distribution plots were made using Rscripts attached to github.

### Annotation and synteny calculation of motifs

The motifs were matched to known transcription factor binding profiles, namely JASPER dataset, using TOMTOM (version 5.5.8) (Gupta et al., 2007) from the MEME suite. The plots were made using ggplot in R.

To assess motif synteny, we implemented a custom R-based pipeline that integrates motif position data derived from MEME (via XSTREME output) and FIMO analyses.

For each orthogroup, motif occurrences were mapped across all orthologous sequences, and their relative positions within each sequence were determined based on their order of appearance. Motifs were first classified into presence-based categories according to their phylogenetic distribution. Motifs present in at least four C_4_ orthologs and absent in all C_3_ orthologs were classified as C_4_-only. Motifs present in both C_4_ and C_3_ species within the PACMAD clade were classified as PACMAD, while those present across C_4_ PACMAD, C_3_ PACMAD, and any of the C_3_ BEP species were assigned to the PACMAD+BEP category. Synteny was evaluated separately within each presence class. For each motif, its relative position within a sequence was defined with respect to the ordered arrangement of all motifs in that sequence. A consensus position was then determined across orthologs within each group (C_4_ or C_3_) based on the most frequently observed position, along with a support metric reflecting positional consistency. Motifs were classified as syntenic if the consensus positions between groups were conserved within a tolerance of ±1 position. Motifs were classified as C_4_-shifted if they exhibited distinct consensus positions between C_4_ and C_3_ groups despite strong positional support within each group. Motifs that did not meet these criteria, but had sufficient representation (more than two motifs within the orthogroup), were classified as *Not-syntenic*. Motifs belonging to orthogroups with insufficient data (≤2 motifs) were categorized as *Not enough motifs* and excluded from synteny interpretation. Motif annotation was done as above.

### Construction of interaction network of Genes and Transcription factors that can bind to them

The interaction network was constructed using a Rscript. The Test genes with annotated C_4_-shifted and Not-syntenic motifs, and the Transcription factors that bind to annotated motifs were used as nodes, while motifs were used for constructing directed edges. Ggraph R package (Pedersen, 2025) was used for graph visualisation in R.

## Scripts, graphics and AI

This flowchart was created using Biorender. We have used AI to generate the framework of scripts for data analysis in Python and R, but the authors have verified and modified these scripts to suit their needs.

## Supplementary Material

**Supplementary Figure S1:** Species tree showing the phylogenetic relationship between the species selected for analysis

**Supplementary Figure S2:** Example of a Test Gene with a complex Tree (GRMZM2G344521)

**Supplementary Figure S3:** Example of a Control Gene with a complex Tree (GRMZM2G066142)

**Supplementary Figure S4:** Example of a (A) Gene-tree with erroneous relationship (indicated in Red) and (B) fixed gene-tree made with a partially constrained tree (GRMZM2G028980).

**Supplementary Figure S5:** Publicly available gene expression atlas data of Bhlh33 (GRMZM2G015666 / Zm00001d005939). Maize embryonic leaf development (Liu et 2022)-(A) absolute and (B) relative; Maize gene atlas (Sekhon et al., 2011)-(C) absolute and (D) relative; (E) Maize leaf development gradient (Li et al., 2010)-relative.

**Supplementary Figure S6:** Publicly available gene expression atlas data of Bhlh105 (GRMZM2G082586 / Zm00001d020790) Maize embryonic leaf development (Liu et 2022)-(A) absolute and (B) relative, Maize gene atlas (Sekhon et al., 2011)- (C) absolute and (D) relative, (E) Maize leaf development gradient (Li et al., 2010)- relative.

**Supplementary Figure S7:** Publicly available gene expression atlas data of Ereb160 (GRMZM2G171179 / Zm00001d045044). Maize embryonic leaf development (Liu et 2022)- (A) absolute and (B) relative, Maize gene atlas (Sekhon et al., 2011)- (C) absolute and (D) relative, (E) Maize leaf development gradient (Li et al., 2010)- relative.

**Supplementary Figure S8:** Publicly available gene expression atlas data of Cadtfr3 (GRMZM2G147712 / Zm00001d050242). Maize embryonic leaf development (Liu et 2022)- (A) absolute and (B) relative, Maize gene atlas (Sekhon et al., 2011)- (C) absolute and (D) relative, (E) Maize leaf development gradient (Li et al., 2010)- relative.

**Supplementary Figure S9:** Publicly available gene expression atlas data of Thx8 (GRMZM2G379179 / Zm00001d031266). Maize embryonic leaf development (Liu et 2022)-(A) absolute and (B) relative, Maize gene atlas (Sekhon et al., 2011)- (C) absolute and (D) relative, (E) Maize leaf development gradient (Li et al., 2010)- relative.

**Supplementary Figure S10:** Publicly available gene expression atlas data of Mads9 (GRMZM2G005155 / Zm00001d002332). Maize embryonic leaf development (Liu et 2022)- (A) absolute and (B) relative, Maize gene atlas (Hoopes et 2018)- (C) relative and (D) Maize leaf development gradient (Li et al., 2010)- relative.

**Supplementary Figure S11:** Publicly available gene expression atlas data of C3H28 (GRMZM2G036837 / Zm00001d008322). Maize embryonic leaf development (Liu et 2022)- (A) absolute and (B) relative, Maize gene atlas (Sekhon et al., 2011)- (C) absolute and (D) relative, (E) Maize leaf development gradient (Li et al., 2010)- relative.

**Supplementary Figure S12:** Publicly available gene expression atlas data of BHLH116 (GRMZM2G042895 / Zm00001d024522). Maize embryonic leaf development (Liu et 2022)- (A) absolute and (B) relative, Maize gene atlas (Sekhon et al., 2011)- (C) absolute and (D) relative, (E) Maize leaf development gradient (Li et al., 2010)- relative.

**Supplementary Figure S13:** Extended gene family tree for BHLH116, showing changes in the position with C4-specific changes across related C3 and C4 species. The outer circle at the end leaves indicates the photosynthetic type of the species (C3, C4 or CAM), and the inner circle indicates the residue present, whether Methionine (M), Valine(V), Alanine (A), Isoleucine (I), or Leucine(L).

**Supplementary Figure S14:** Comprehensive expression profiles in multiple RNAseq studies. The left panel shows the expression in rice leaf primordia (van-Campen et al. 2016), the middle panel shows expression of the gene in maize foliar and husk leaves (Wang et al., 2013), and the right panel shows the expression of gene in early leaf primordia (Knauer S, et al. 2019). A to H are the genes in the order of bHLH33, bHLH105, EREB160, CADTFR3, THX8, MADS9, C3H28, bHLH116.

**Supplementary Figure S15:** Mean frequency of motif distribution per orthogroup across 10 replicates of 55 randomly sampled test orthogroups, compared with the control dataset.

**Supplementary Figure S16:** Total vs annotated motifs per orthogroups for test(Analysis) and control dataset.

**Supplementary Table S1:** Genes selected from study 1 (Wang et al., 2013).

**Supplementary Table S2:** Genes selected from study 2 (Liu et al., 2022).

**Supplementary Table S3:** Genes selected for control set.

**Supplementary Table S4:** Maize gene ID of the orthogroups that formed unresolvable complex gene-trees.

**Supplementary Table S5:** Likelihood Ratio Test (LRT) results of the orthogroups from test and control.

**Supplementary Table S6:** Functional annotation of Genes that passed the LRT test.

**Supplementary Table S7:** Enriched motifs matching the previously identified motifs driving C4-specific expression.

**Supplementary Table S9:** Families enriched in control and test sets.

**Supplementary Table S10:** Motifs that are Non-syntenic or C4 shifted in test and control.

## Acknowledgments

This work was funded by DBT-ramalingaswami grant for “Discovery of gene missing components of regulatory network underlying C_4_ pathway/anatomy for translational research” and author Vivek Thakur is grateful for this. Author Angeo Saji was also supported by CSIR-JRF, and is deeply grateful for their support. Author Angeo Saji thanked Nikhitha R and Rhydima Pal (Department of Systems and Computational Biology, School of Life Sciences, University of Hyderabad) for their technical support during data analysis.

## Data availability

All the scripts used, tree files, alignment, and partially constrained species tree are available in a public GitHub repository (Saji 2026) at: https://github.com/angeosaji/Adaptive-evolution-Trees-and-Alignments-and-species-tree.git https://github.com/angeosaji/postitve_selection_scripts.git https://github.com/angeosaji/Cis-motif-analysis.git

## Author contributions

V.T. conceived, designed, and supervised the study. A.S. generated and analyzed the data of transcription factors for evidence of positive selection. G.B. generated data for cis-regulatory elements. A.S and G. B. analyzed the data for cis-regulatory elements. A.S. wrote the first draft. A.S. and V.T. reviewed and finalized it. All authors read and approved the manuscript.

## Abbreviations

Ka: Nonsynonymous substitution rate
Ks: Synonymous substitution rate
SAM: Shoot Apical Meristem
CRE: *Cis-*regulatory elements

