## Supplementary data for "Using adaptive evolutionary signatures to screen candidate regulatory genes and cis-elements associated with Kranz development in monocots"

### Index

### Supplementary Figures

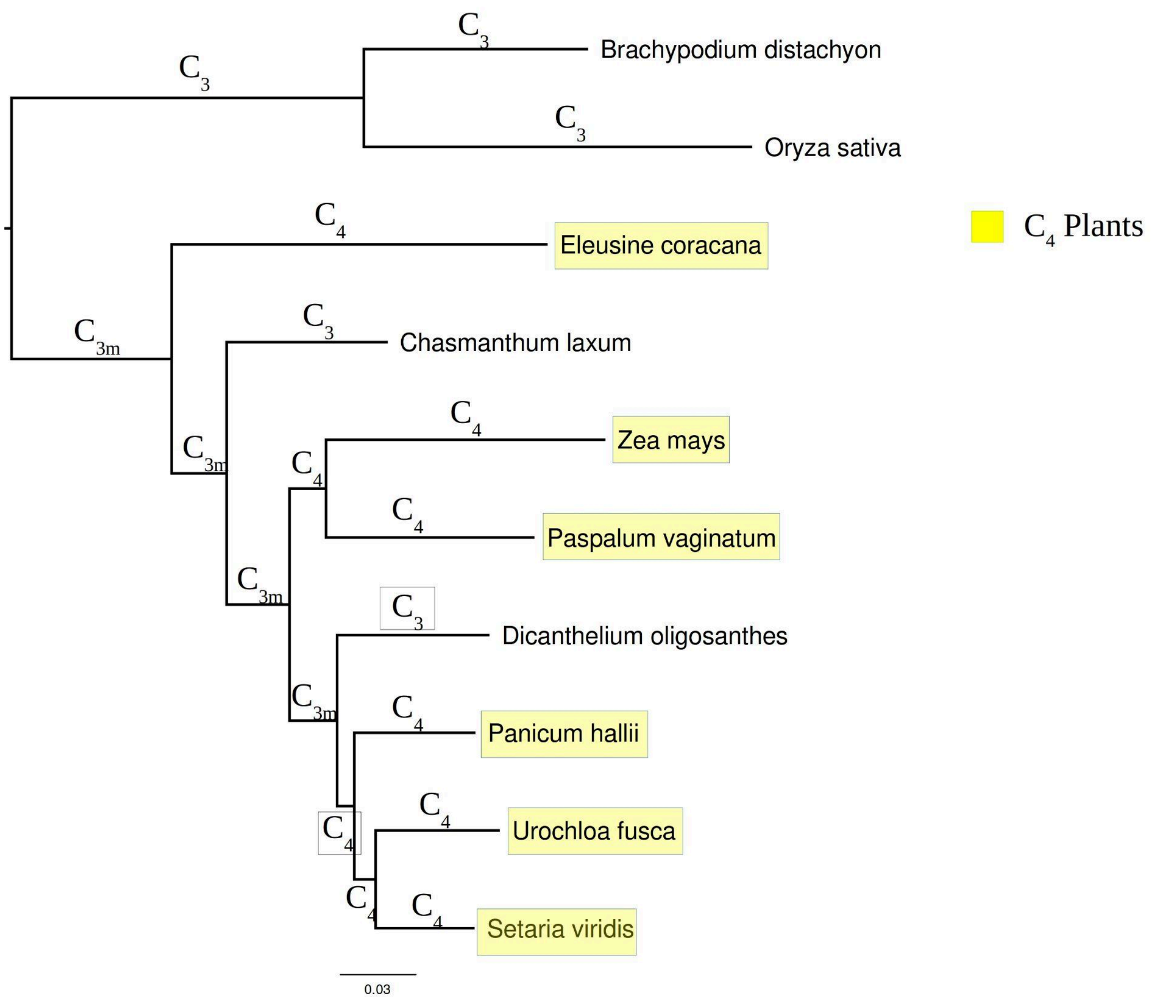

**Supplementary Figure S1:**

**Species tree showing the phylogenetic relationship between the species selected for analysis.** C3 nodes are nodes with purely C3 ancestors/plant, C3m nodes are nodes with mixed ancestry, and C4 nodes are nodes with purely C4 ancestors/plant

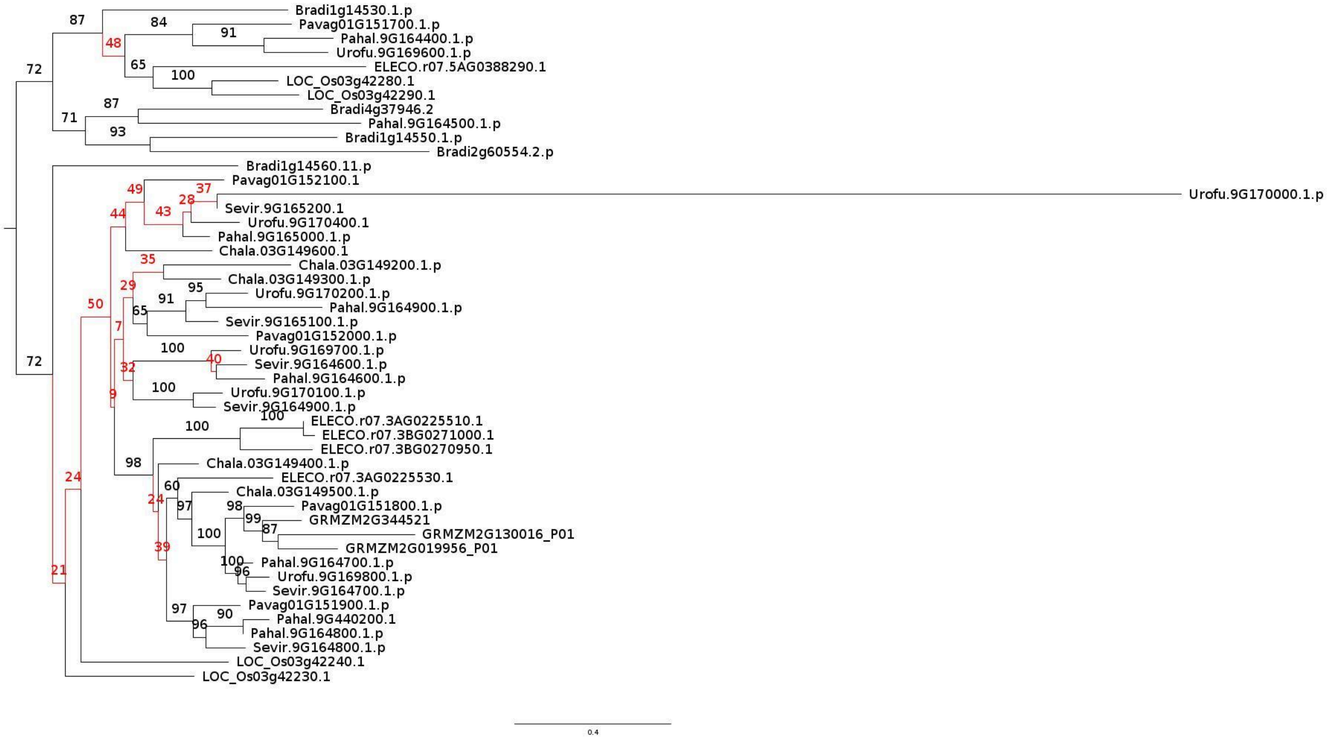

Supplementary Figure S2:

Example of a Test Gene with a complex Tree (GRMZM2G344521)

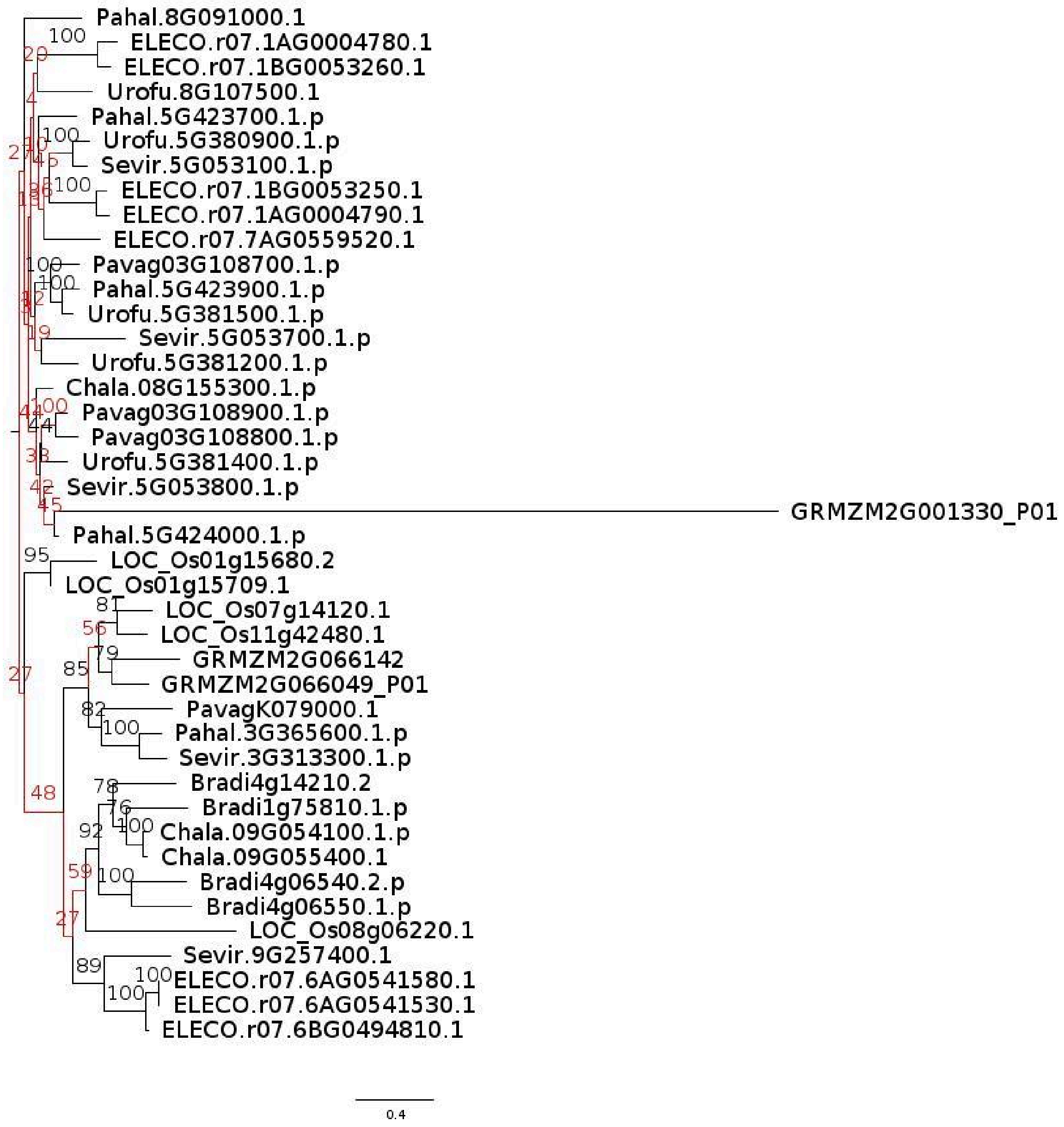

Supplementary Figure S3:

Example of a Control Gene with a complex Tree (GRMZM2G066142)

A.

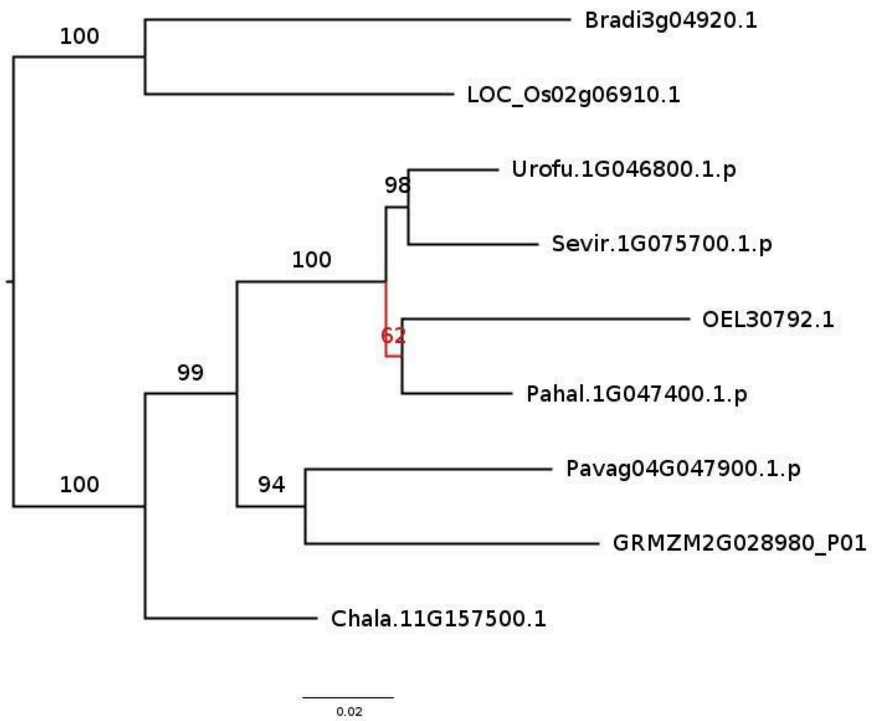

B.

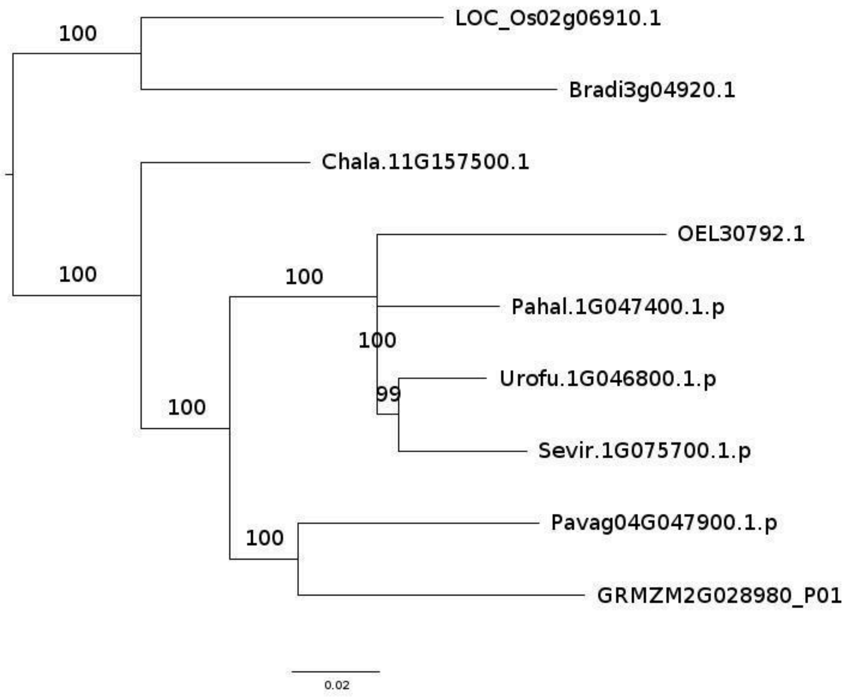

Supplementary Figure S4:

Example of a (A) Gene-tree with erroneous relationship (indicated in Red) and (B) fixed gene-tree made with a partially constrained tree (GRMZM2G028980).

A.

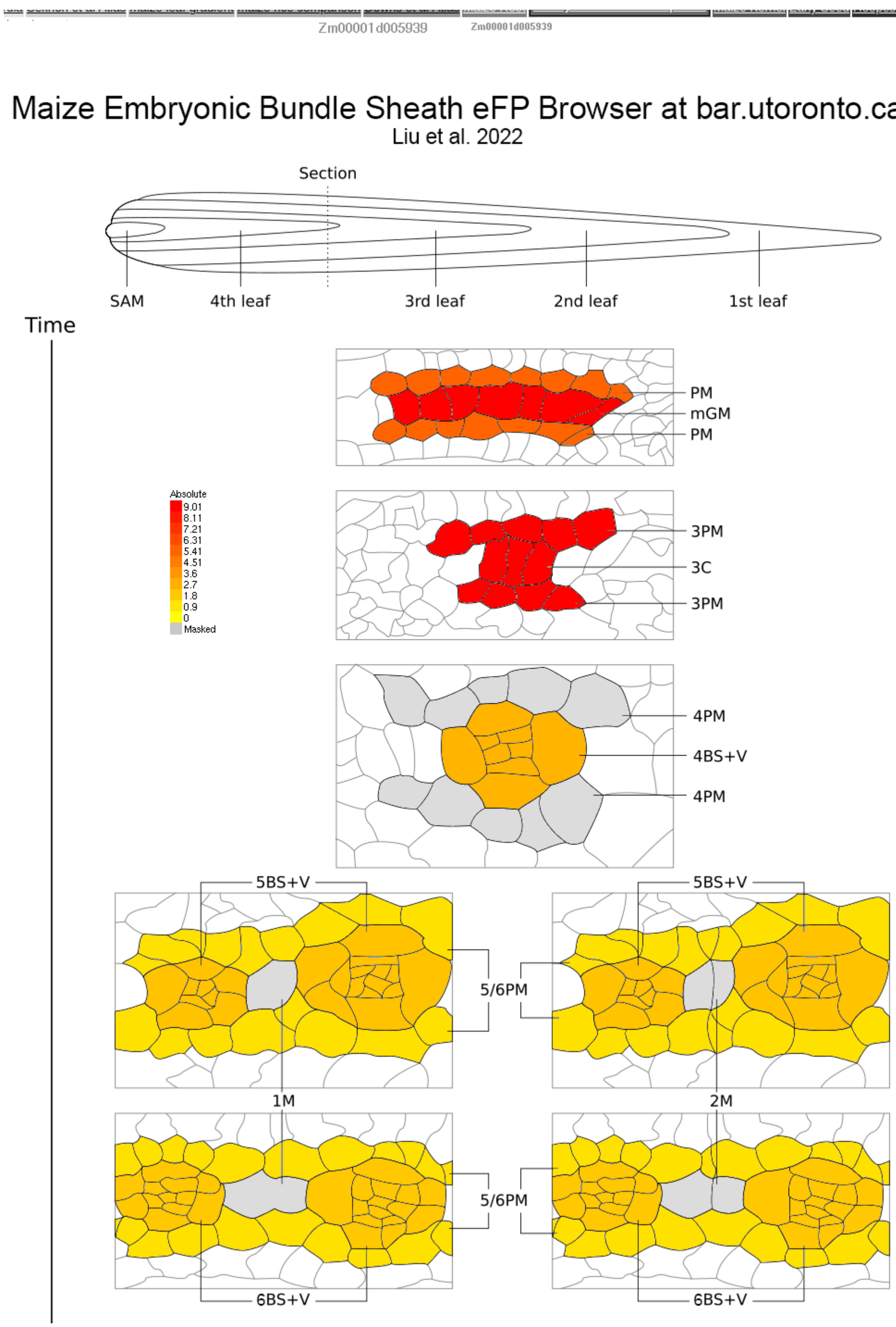
Zm00001d005939
Zm00001d005939
Maize Embryonic Bundle Sheath eFP Browser at bar.utoronto.ca
Liu et al. 2022
Section
SAM
4th leaf
3rd leaf
2nd leaf
1st leaf
Time
PM
mGM
PM
Absolute
9.01
8.11
7.21
6.31
5.41
4.51
3.6
2.7
1.8
0.9
0
Masked
3PM
3C
3PM
4PM
4BS+V
4PM
5BS+V
5BS+V
5/6PM
1M
2M
5/6PM
6BS+V
6BS+V

B.

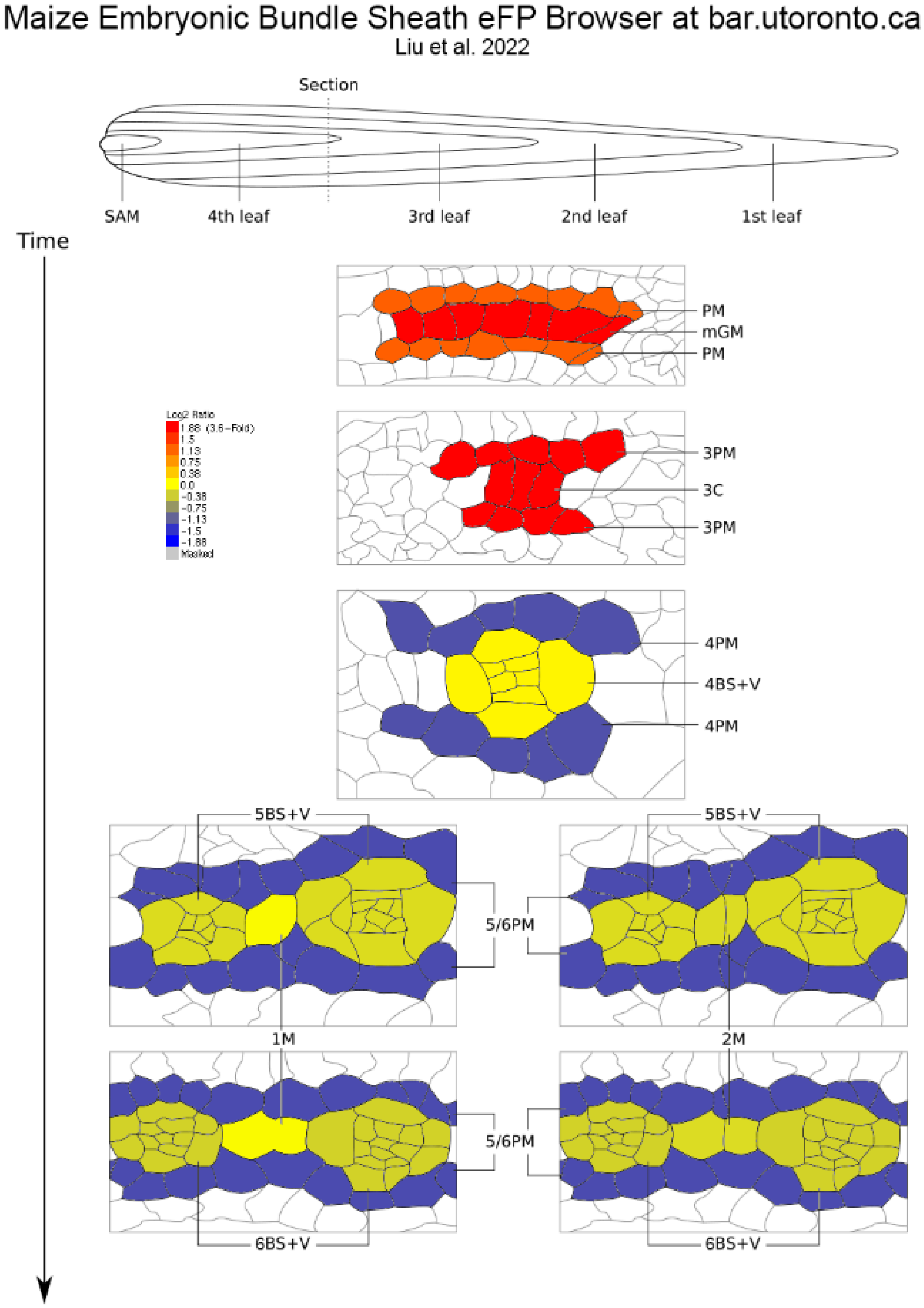
Maize Embryonic Bundle Sheath eFP Browser at bar.utoronto.ca
Liu et al. 2022
Section
SAM
4th leaf
3rd leaf
2nd leaf
1st leaf
Time
PM
mGM
PM
Log2 Ratio
1.88 (3.6-Fold)
1.5
1.13
0.75
0.38
0.0
-0.38
-0.75
-1.13
-1.5
-1.88
Masked
3PM
3C
3PM
4PM
4BS+V
4PM
5BS+V
5BS+V
5/6PM
1M
2M
5/6PM
6BS+V
6BS+V

C.

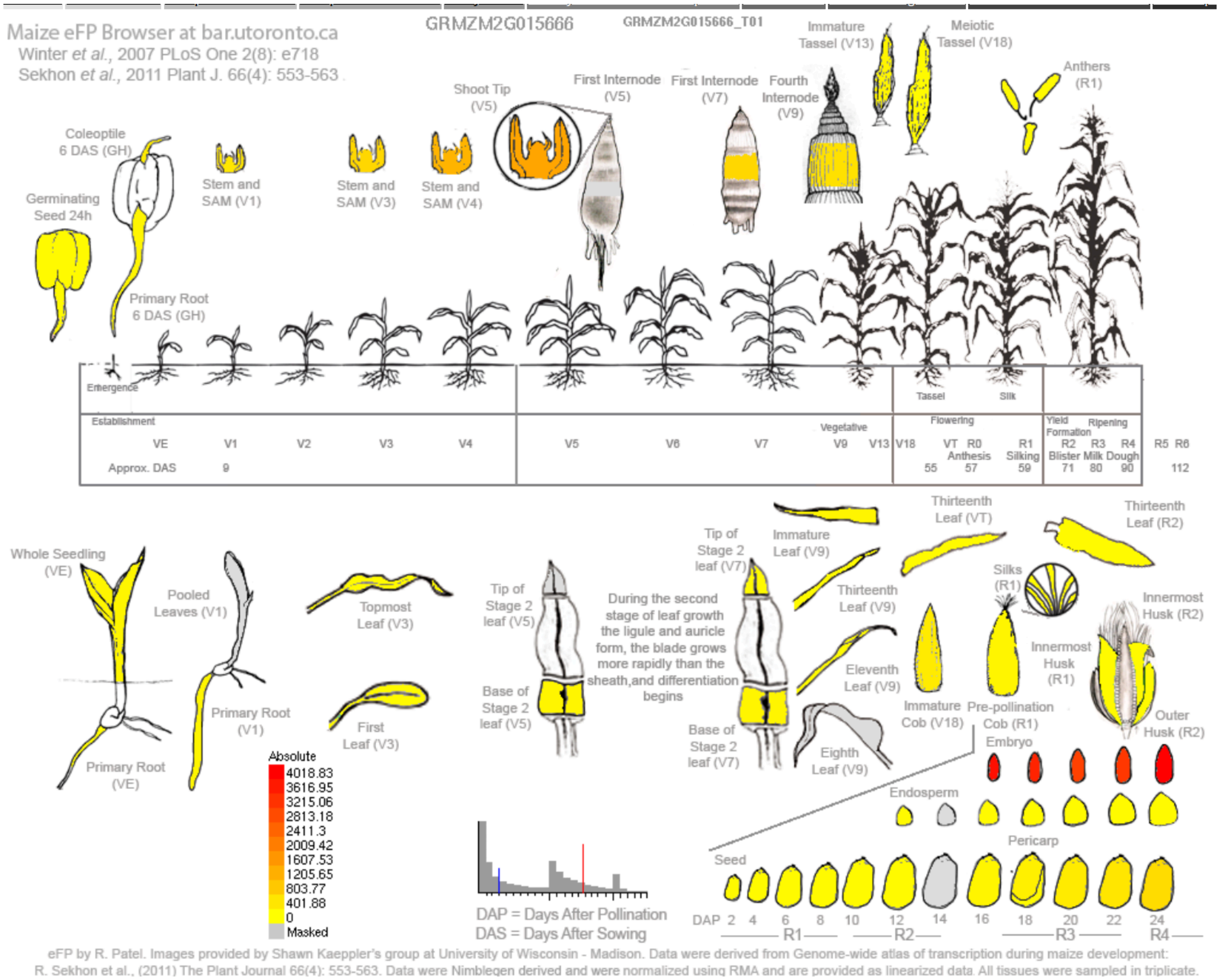

Maize eFP Browser at bar.utoronto.ca
Winter et al., 2007 PLoS One 2(8): e718
Sekhon et al., 2011 Plant J. 66(4): 553-563
GRMZM2G015666
GRMZM2G015666_T01
Immature Tassel (V13)
Meiotic Tassel (V18)
Anthers (R1)
Shoot Tip (V5)
First Internode (V5)
First Internode (V7)
Fourth Internode (V9)
Coleoptile 6 DAS (GH)
Stem and SAM (V1)
Stem and SAM (V3)
Stem and SAM (V4)
Germinating Seed 24h
Primary Root 6 DAS (GH)
Emergence
Tassel
Silk
Establishment
Vegetative
Flowering
Yield Formation
Ripening
VE V1 V2 V3 V4 V5 V6 V7 V9 V13 V18 VT R0 R1 R2 R3 R4 R5 R6
Anthesis
Silking
Blister Milk Dough
Approx. DAS 9 55 57 59 71 80 90 112
Whole Seedling (VE)
Pooled Leaves (V1)
Topmost Leaf (V3)
First Leaf (V3)
Primary Root (V1)
Primary Root (VE)
Tip of Stage 2 leaf (V5)
Base of Stage 2 leaf (V5)
During the second stage of leaf growth the ligule and auricle form, the blade grows more rapidly than the sheath,and differentiation begins
Tip of Stage 2 leaf (V7)
Base of Stage 2 leaf (V7)
Immature Leaf (V9)
Thirteenth Leaf (V9)
Eleventh Leaf (V9)
Eighth Leaf (V9)
Thirteenth Leaf (VT)
Thirteenth Leaf (R2)
Silks (R1)
Innermost Husk (R2)
Innermost Husk (R1)
Outer Husk (R2)
Immature Cob (V18)
Pre-pollination Cob (R1)
Embryo
Endosperm
Pericarp
Seed
Absolute
4018.83
3616.95
3215.06
2813.18
2411.3
2009.42
1607.53
1205.65
803.77
401.88
0
Masked
DAP = Days After Pollination
DAS = Days After Sowing
DAP 2 4 6 8 10 12 14 16 18 20 22 24
R1 R2 R3 R4
eFP by R. Patel. Images provided by Shawn Kaeppler's group at University of Wisconsin - Madison. Data were derived from Genome-wide atlas of transcription during maize development: R. Sekhon et al., (2011) The Plant Journal 66(4): 553-563. Data were Nimblegen derived and were normalized using RMA and are provided as linearized data. All tissues were sampled in triplicate.

D.

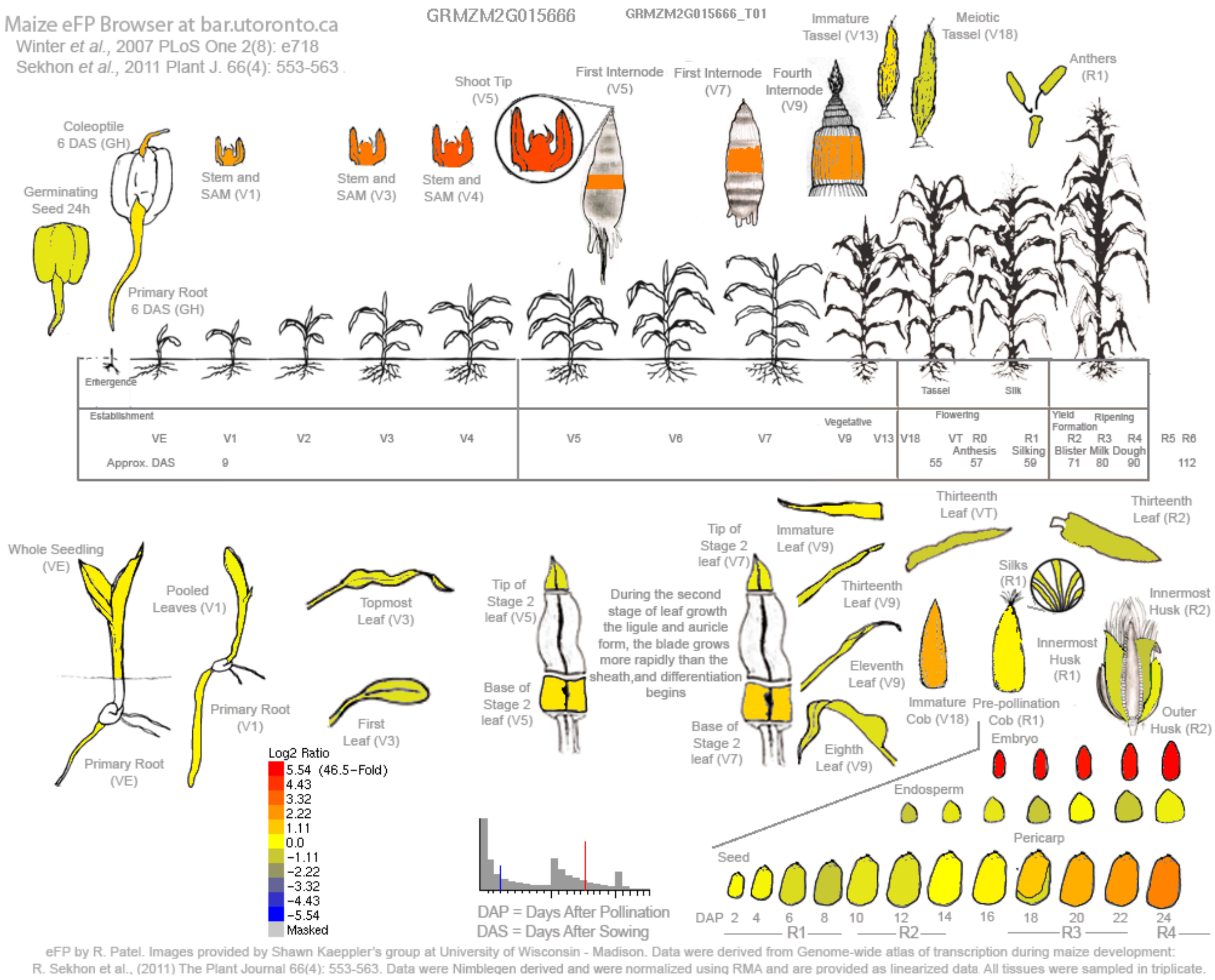
Maize eFP Browser at bar.utoronto.ca
Winter et al., 2007 PLoS One 2(8): e718
Sekhon et al., 2011 Plant J. 66(4): 553-563
GRMZM2G015666
GRMZM2G015666_T01
Log2 Ratio
5.54 (46.5-Fold)
4.43
3.32
2.22
1.11
0.0
-1.11
-2.22
-3.32
-4.43
-5.54
Masked
DAP = Days After Pollination
DAS = Days After Sowing
eFP by R. Patel. Images provided by Shawn Kaeppler's group at University of Wisconsin - Madison. Data were derived from Genome-wide atlas of transcription during maize development:
R. Sekhon et al., (2011) The Plant Journal 66(4): 553-563. Data were Nimblegen derived and were normalized using RMA and are provided as linearized data. All tissues were sampled in triplicate.

E.

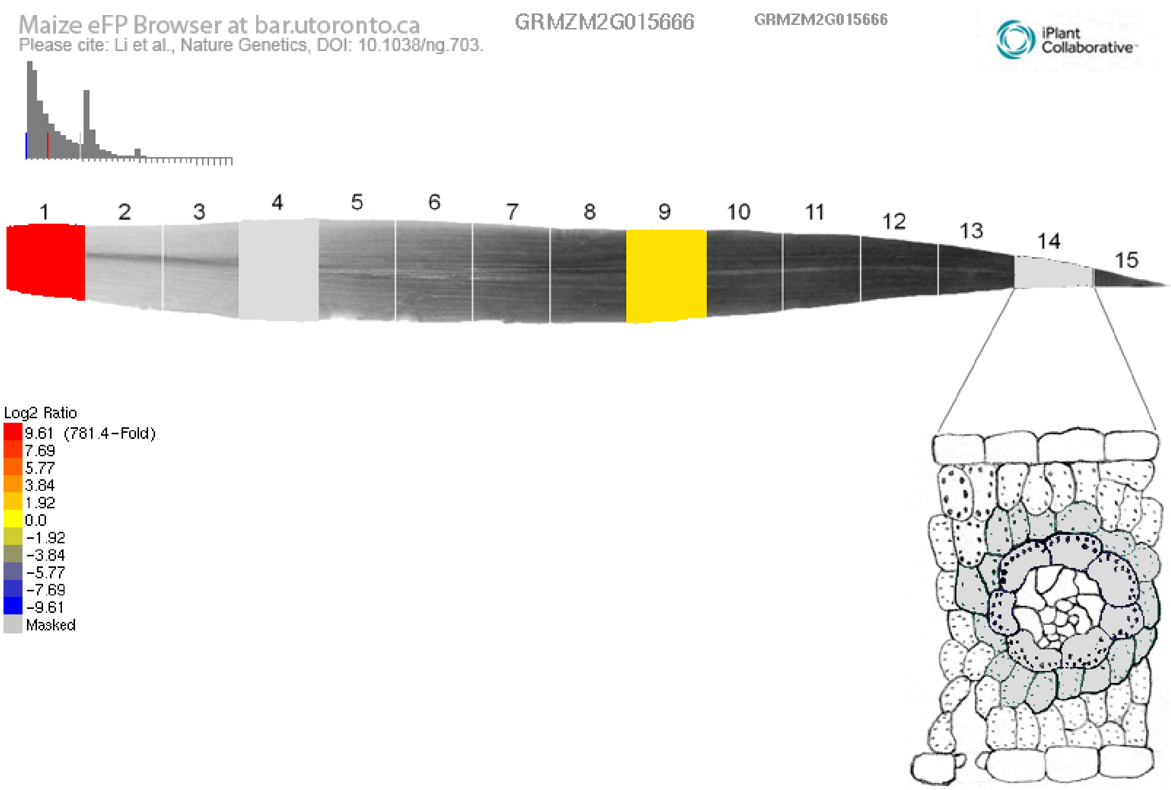

Supplementary Figure S5:

Publicly available gene expression atlas data of **Bhlh33** (GRMZM2G015666 / Zm00001d005939). Maize embryonic leaf development (Liu et 2022)- (A) absolute and (B) relative, Maize gene atlas (Sekhon et al., 2011)- (C) absolute and (D) relative, (E) Maize leaf development gradient (Li et al., 2010)- relative.

A.

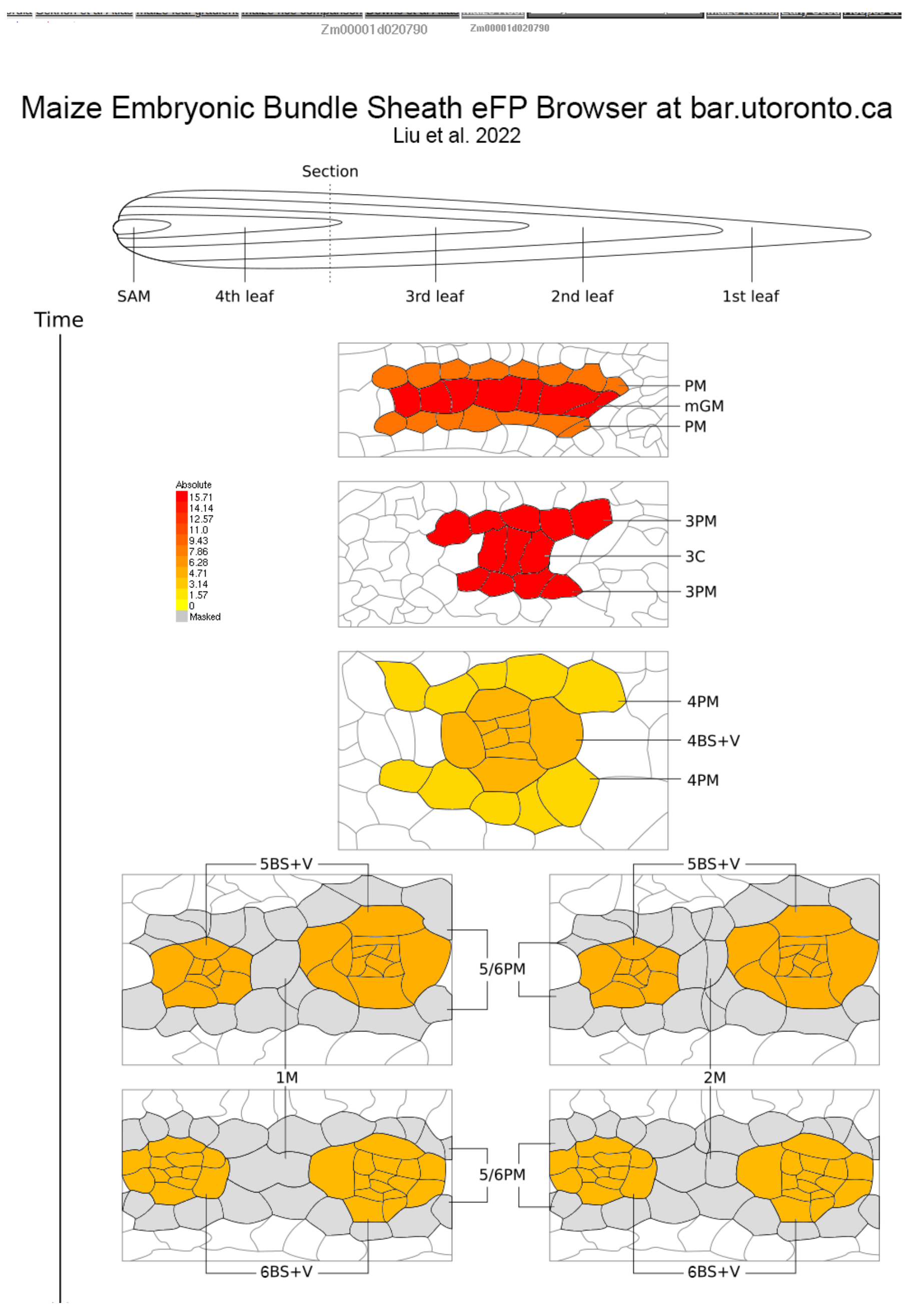
Zm00001d020790
Maize Embryonic Bundle Sheath eFP Browser at bar.utoronto.ca
Liu et al. 2022
Section
SAM
4th leaf
3rd leaf
2nd leaf
1st leaf
Time
PM
mGM
PM
Absolute
15.71
14.14
12.57
11.0
9.43
7.86
6.28
4.71
3.14
1.57
0
Masked
3PM
3C
3PM
4PM
4BS+V
4PM
5BS+V
5BS+V
5/6PM
1M
2M
5/6PM
6BS+V
6BS+V

B.

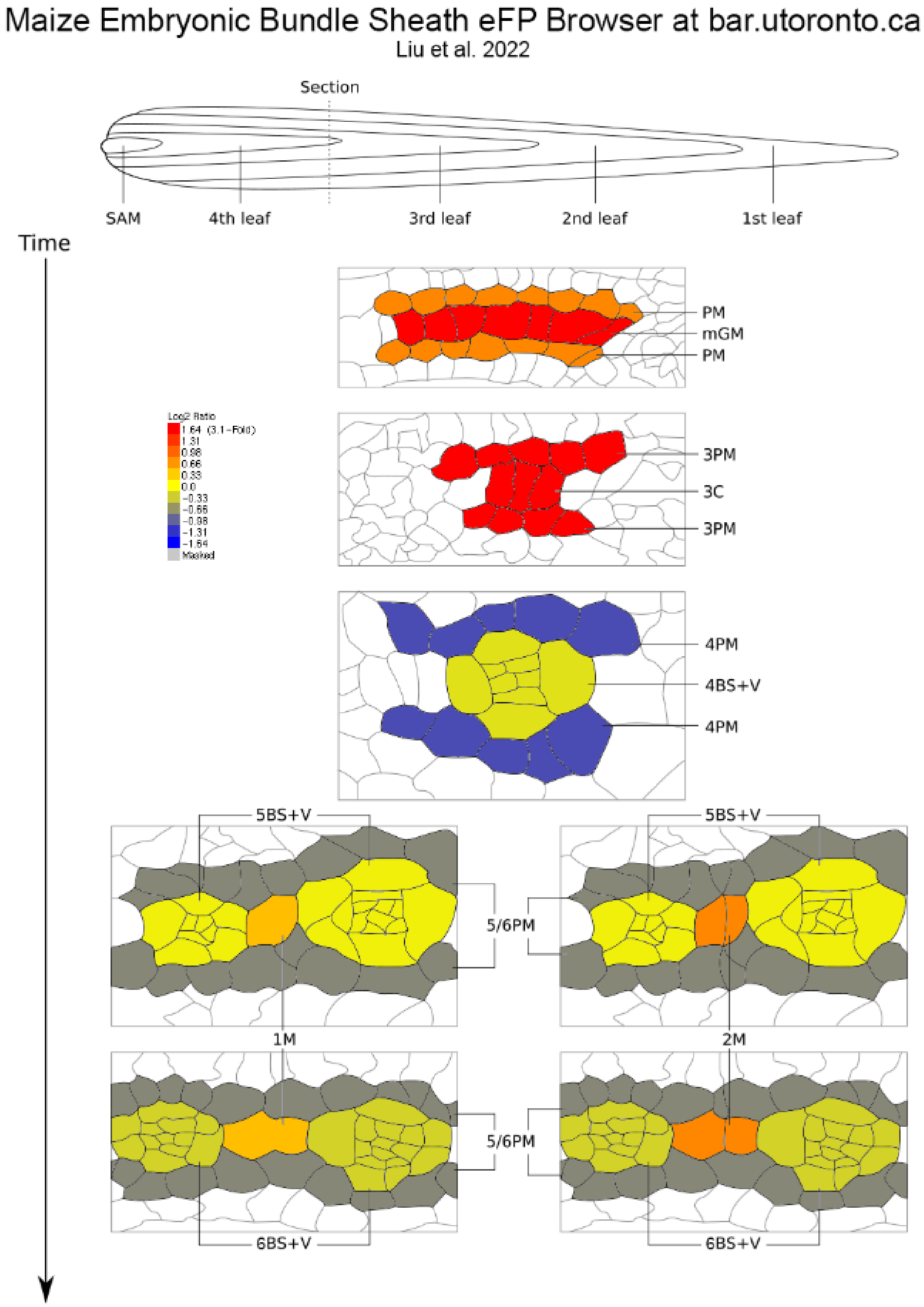
Maize Embryonic Bundle Sheath eFP Browser at bar.utoronto.ca
Liu et al. 2022
Section
SAM
4th leaf
3rd leaf
2nd leaf
1st leaf
Time
PM
mGM
PM
Log2 Ratio
1.64 (3.1-Fold)
1.31
0.98
0.66
0.33
0.0
-0.33
-0.66
-0.98
-1.31
-1.64
Masked
3PM
3C
3PM
4PM
4BS+V
4PM
5BS+V
5BS+V
5/6PM
1M
2M
5/6PM
6BS+V
6BS+V

C.

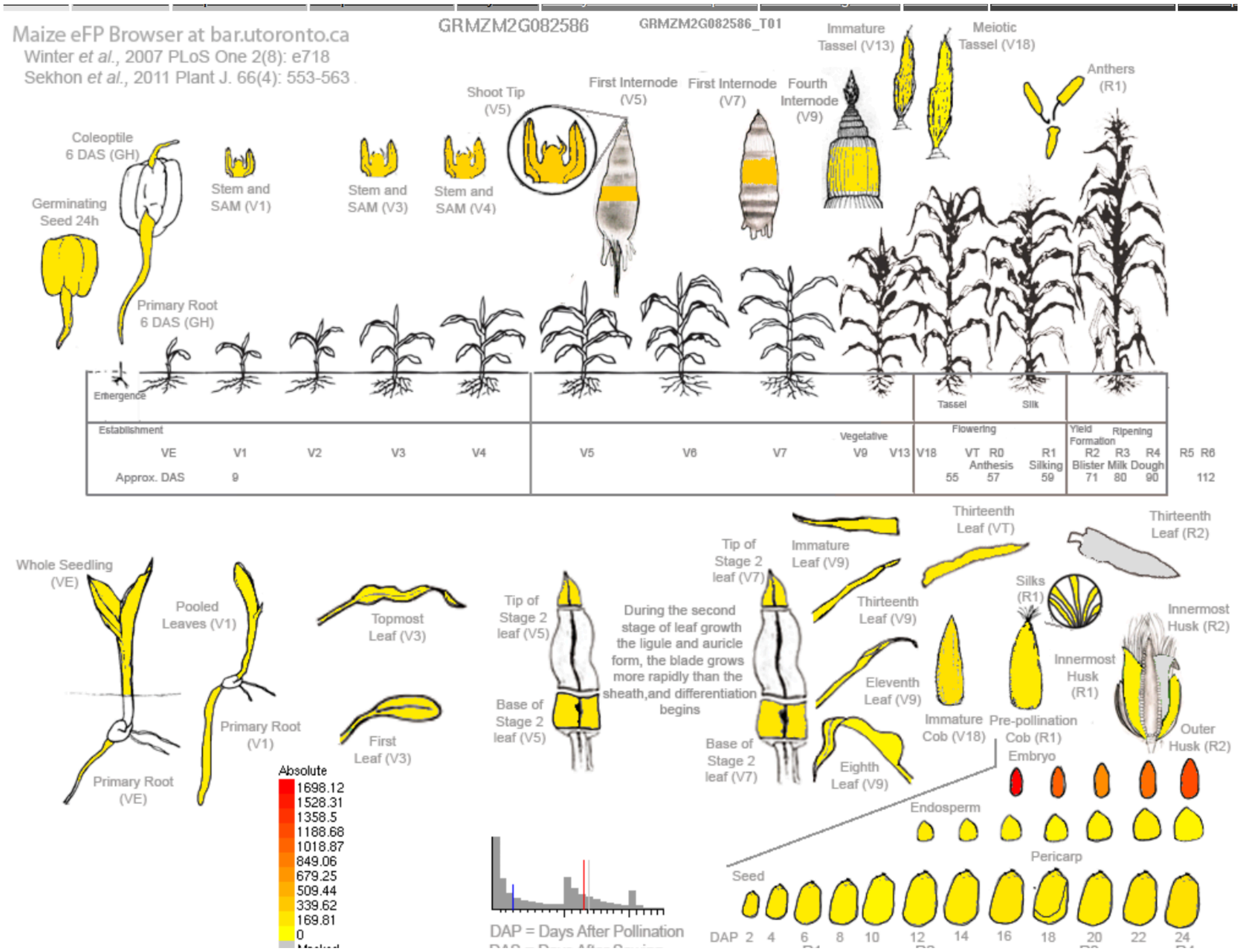
Maize eFP Browser at bar.utoronto.ca
Winter et al., 2007 PLoS One 2(8): e718
Sekhon et al., 2011 Plant J. 66(4): 553-563
GRMZM2G082586
GRMZM2G082586_T01
Immature Tassel (V13)
Meiotic Tassel (V18)
Anthers (R1)
Shoot Tip (V5)
First Internode (V5)
First Internode (V7)
Fourth Internode (V9)
Coleoptile 6 DAS (GH)
Germinating Seed 24h
Stem and SAM (V1)
Stem and SAM (V3)
Stem and SAM (V4)
Primary Root 6 DAS (GH)
Emergence
Tassel
Silk
Establishment
Vegetative
Flowering
Yield Formation
Ripening
VE V1 V2 V3 V4 V5 V6 V7 V9 V13 V18 VT R0 Anthesis R1 Silking R2 Blister R3 Milk R4 Dough R5 R6
Approx. DAS 9 55 57 59 71 80 90 112
Whole Seedling (VE)
Pooled Leaves (V1)
Topmost Leaf (V3)
First Leaf (V3)
Primary Root (V1)
Primary Root (VE)
Tip of Stage 2 leaf (V5)
Base of Stage 2 leaf (V5)
Tip of Stage 2 leaf (V7)
Base of Stage 2 leaf (V7)
During the second stage of leaf growth the ligule and auricle form, the blade grows more rapidly than the sheath,and differentiation begins
Immature Leaf (V9)
Thirteenth Leaf (V9)
Eleventh Leaf (V9)
Eighth Leaf (V9)
Thirteenth Leaf (VT)
Thirteenth Leaf (R2)
Silks (R1)
Innermost Husk (R2)
Innermost Husk (R1)
Outer Husk (R2)
Immature Cob (V18)
Pre-pollination Cob (R1)
Embryo
Endosperm
Pericarp
Seed
DAP 2 4 6 8 10 12 14 16 18 20 22 24
Absolute
1698.12
1528.31
1358.5
1188.68
1018.87
849.06
679.25
509.44
339.62
169.81
0
DAP = Days After Pollination

D.

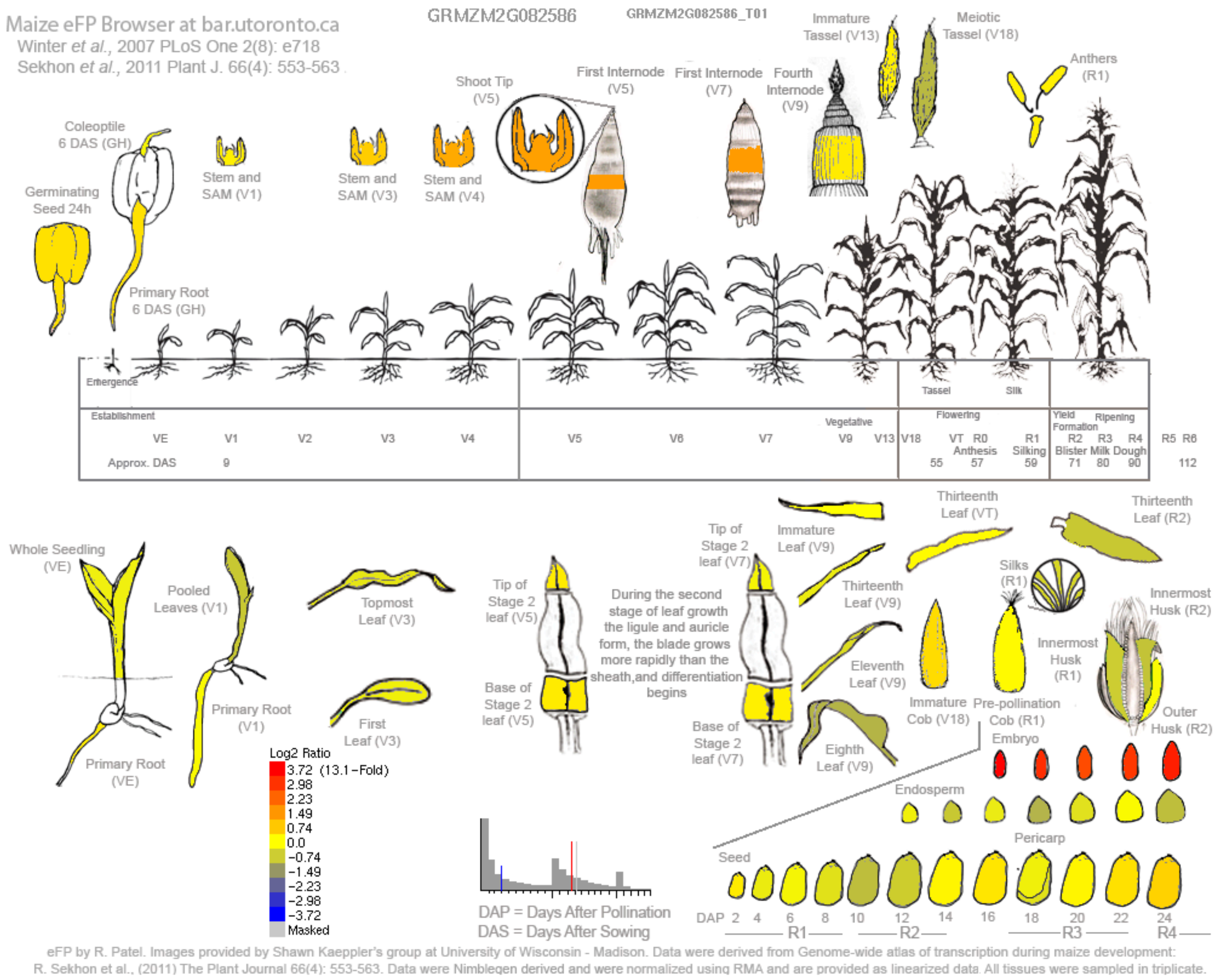

Maize eFP Browser at bar.utoronto.ca
Winter et al., 2007 PLoS One 2(8): e718
Sekhon et al., 2011 Plant J. 66(4): 553-563
GRMZM2G082586
GRMZM2G082586_T01
Immature Tassel (V13)
Meiotic Tassel (V18)
Anthers (R1)
Shoot Tip (V5)
First Internode (V5)
First Internode (V7)
Fourth Internode (V9)
Coleoptile 6 DAS (GH)
Germinating Seed 24h
Stem and SAM (V1)
Stem and SAM (V3)
Stem and SAM (V4)
Primary Root 6 DAS (GH)
Emergence
Establishment
Vegetative
Flowering
Yield Formation
Ripening
Tassel
Silk
VE V1 V2 V3 V4 V5 V6 V7 V9 V13 V18 VT R0 R1 R2 R3 R4 R5 R6
Anthesis Silking Blister Milk Dough
Approx. DAS 9 55 57 59 71 80 90 112
Whole Seedling (VE)
Pooled Leaves (V1)
Topmost Leaf (V3)
First Leaf (V3)
Primary Root (V1)
Primary Root (VE)
Tip of Stage 2 leaf (V5)
Base of Stage 2 leaf (V5)
During the second stage of leaf growth the ligule and auricle form, the blade grows more rapidly than the sheath,and differentiation begins
Tip of Stage 2 leaf (V7)
Base of Stage 2 leaf (V7)
Immature Leaf (V9)
Thirteenth Leaf (VT)
Thirteenth Leaf (R2)
Thirteenth Leaf (V9)
Eleventh Leaf (V9)
Eighth Leaf (V9)
Silks (R1)
Innermost Husk (R2)
Innermost Husk (R1)
Immature Cob (V18)
Pre-pollination Cob (R1)
Outer Husk (R2)
Embryo
Endosperm
Pericarp
Seed
Log2 Ratio
3.72 (13.1-Fold)
2.98
2.23
1.49
0.74
0.0
-0.74
-1.49
-2.23
-2.98
-3.72
Masked
DAP = Days After Pollination
DAS = Days After Sowing
DAP 2 4 6 8 10 12 14 16 18 20 22 24
R1 R2 R3 R4
eFP by R. Patel. Images provided by Shawn Kaeppler's group at University of Wisconsin - Madison. Data were derived from Genome-wide atlas of transcription during maize development: R. Sekhon et al., (2011) The Plant Journal 66(4): 553-563. Data were Nimblegen derived and were normalized using RMA and are provided as linearized data All tissues were sampled in triplicate.

E.

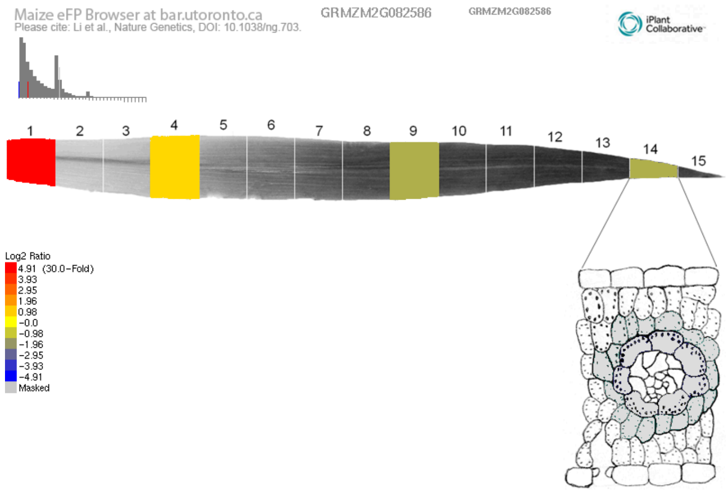

Supplementary Figure S6:

Publicly available gene expression atlas data of **Bhlh105** (GRMZM2G082586 / Zm00001d020790), Maize embryonic leaf development (Liu et 2022)- (A) absolute and (B) relative, Maize gene atlas (Sekhon et al., 2011)- (C) absolute and (D) relative, (E) Maize leaf development gradient (Li et al., 2010)- relative.

A. Maize embryonic leaf development (Liu et 2022)- Absolute

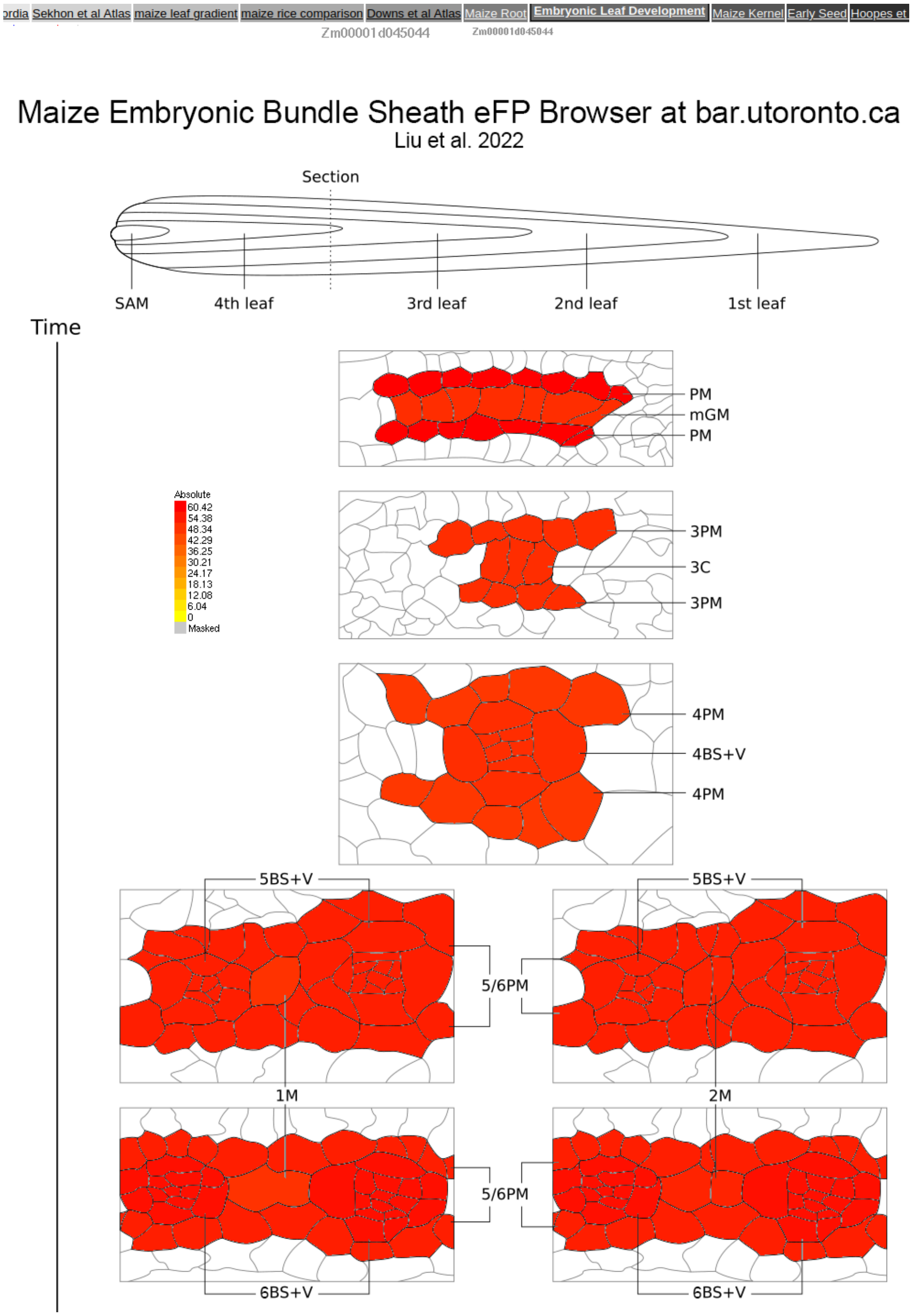

B. Maize embryonic leaf development (Liu et 2022)- Relative

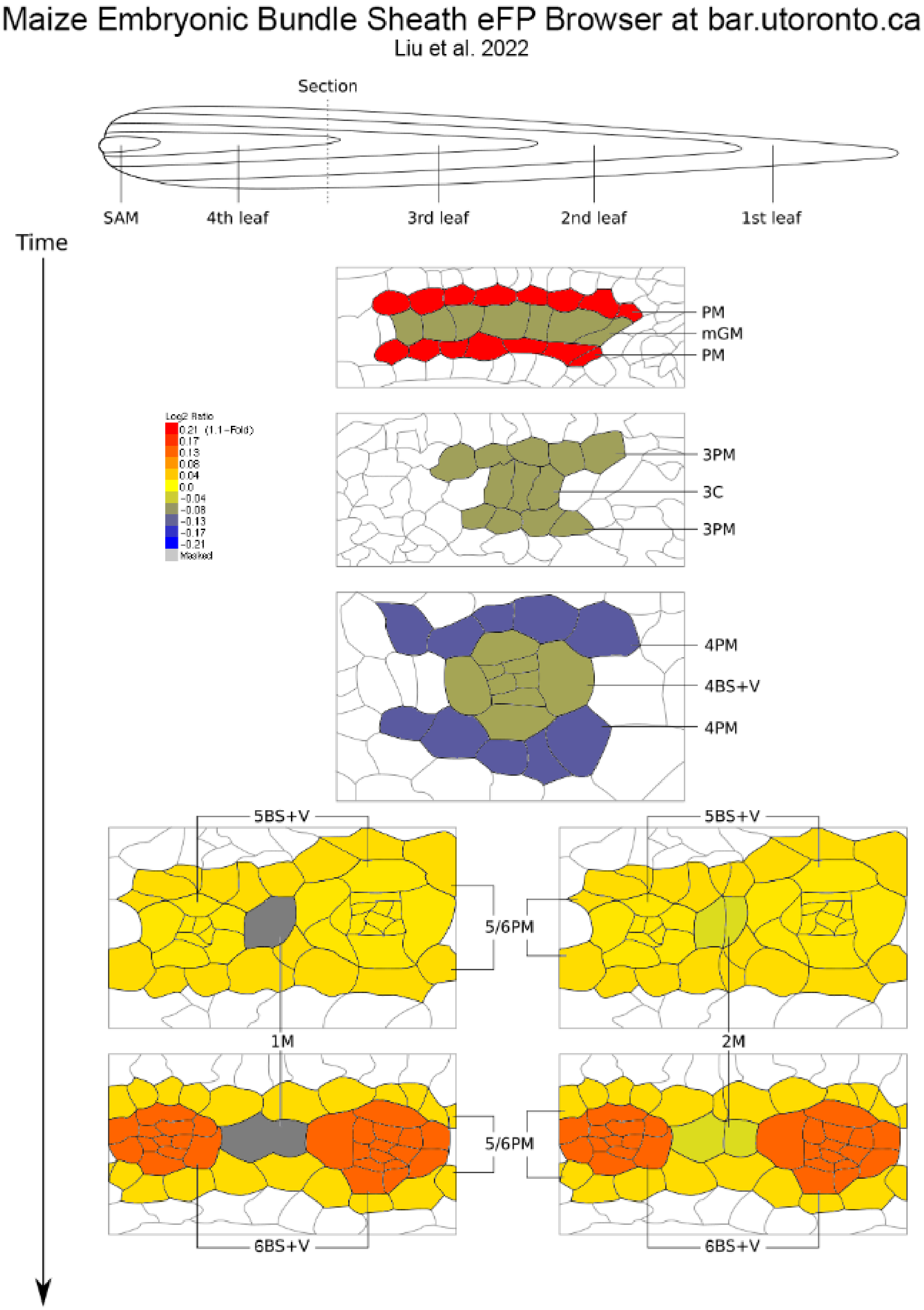

C. Maize gene atlas (Sekhon et al., 2011)- Absolute

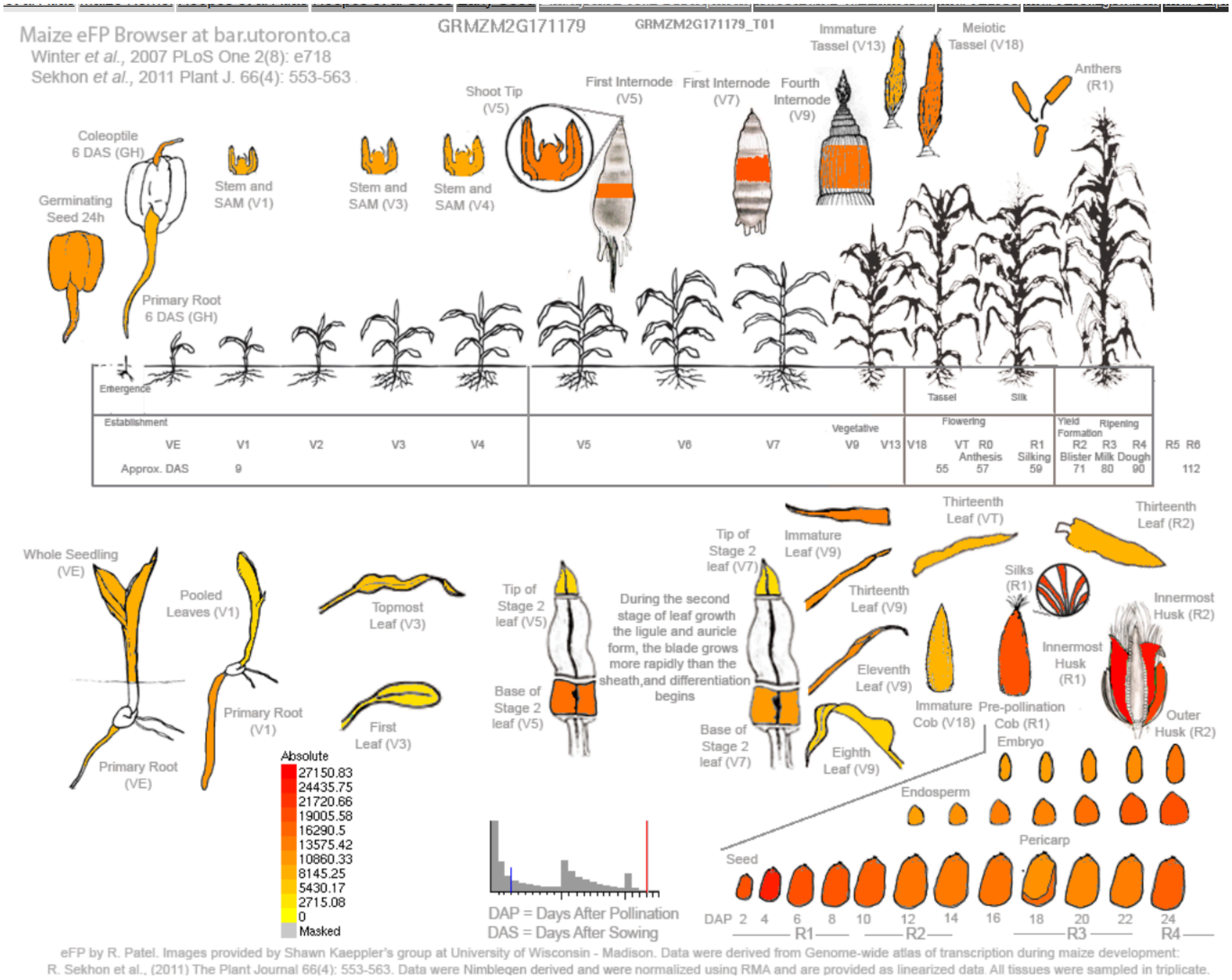

D.  Maize gene atlas (Sekhon et al., 2011)- Relative

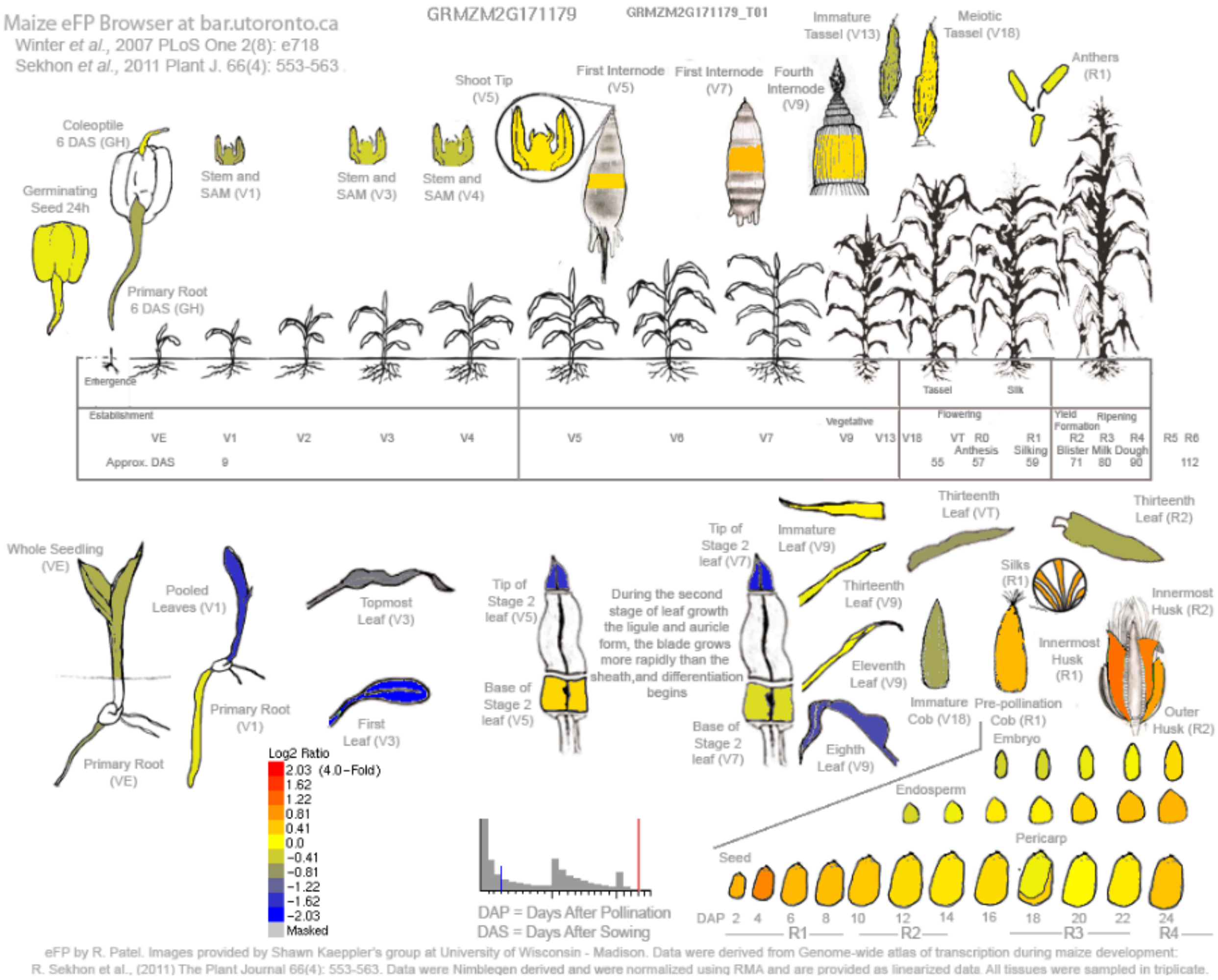

E. Maize leaf development gradient (Li et al., 2010)- Relative

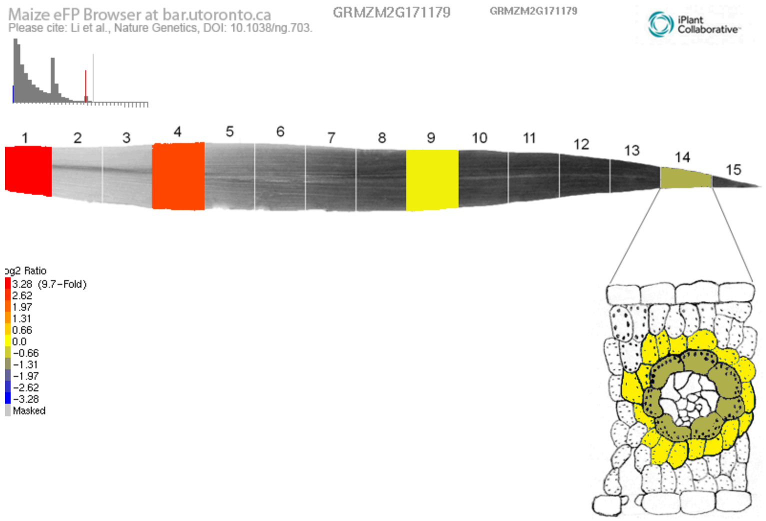

Supplementary Figure S7:

Publicly available gene expression atlas data of **Ereb160** (GRMZM2G171179 / Zm00001d045044). Maize embryonic leaf development (Liu et 2022)- (A) absolute and (B) relative, Maize gene atlas (Sekhon et al., 2011)- (C) absolute and (D) relative, (E) Maize leaf development gradient (Li et al., 2010)- relative.

A. Maize embryonic leaf development (Liu et 2022)- Absolute

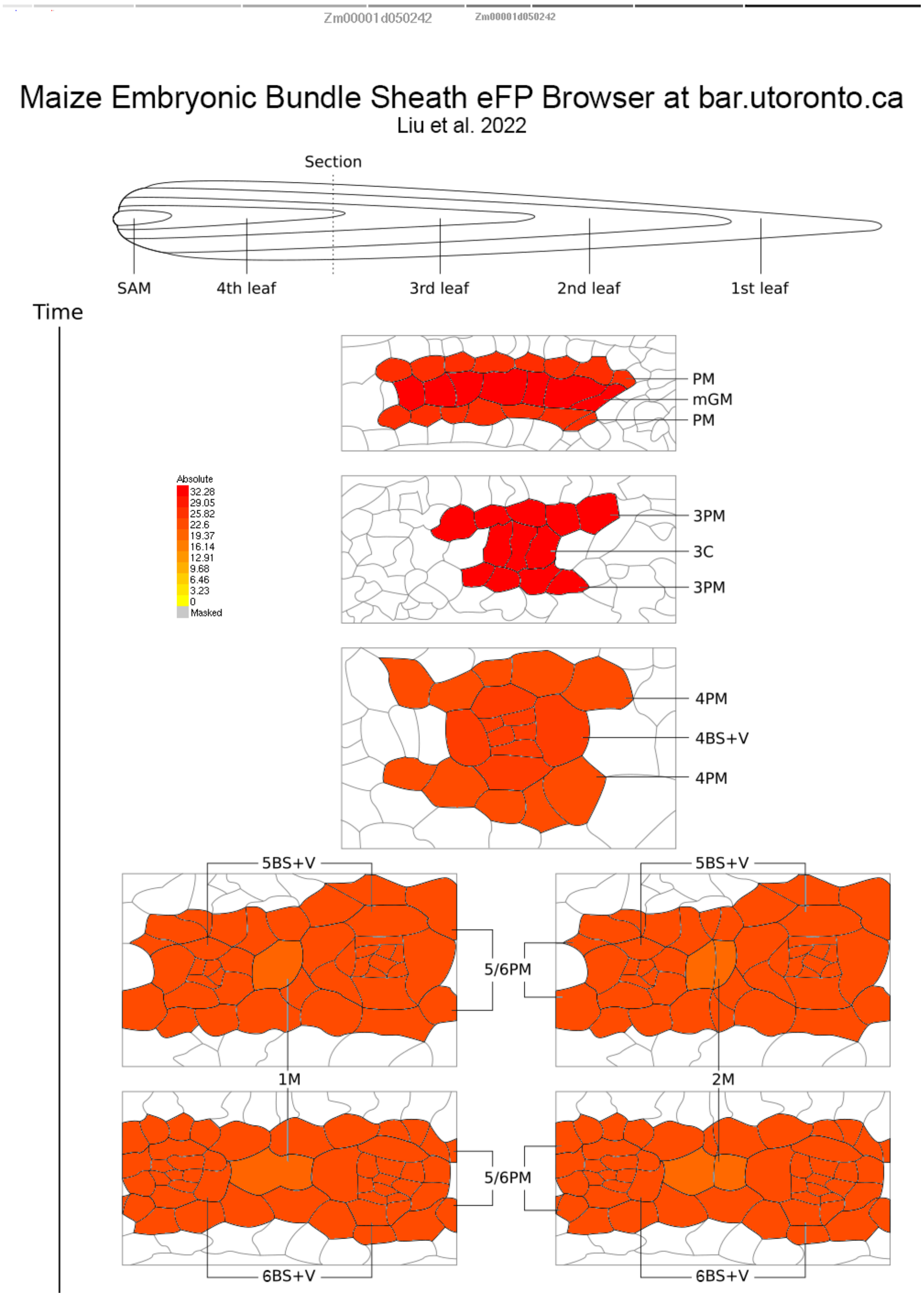

B. Maize embryonic leaf development (Liu et 2022)- Relative

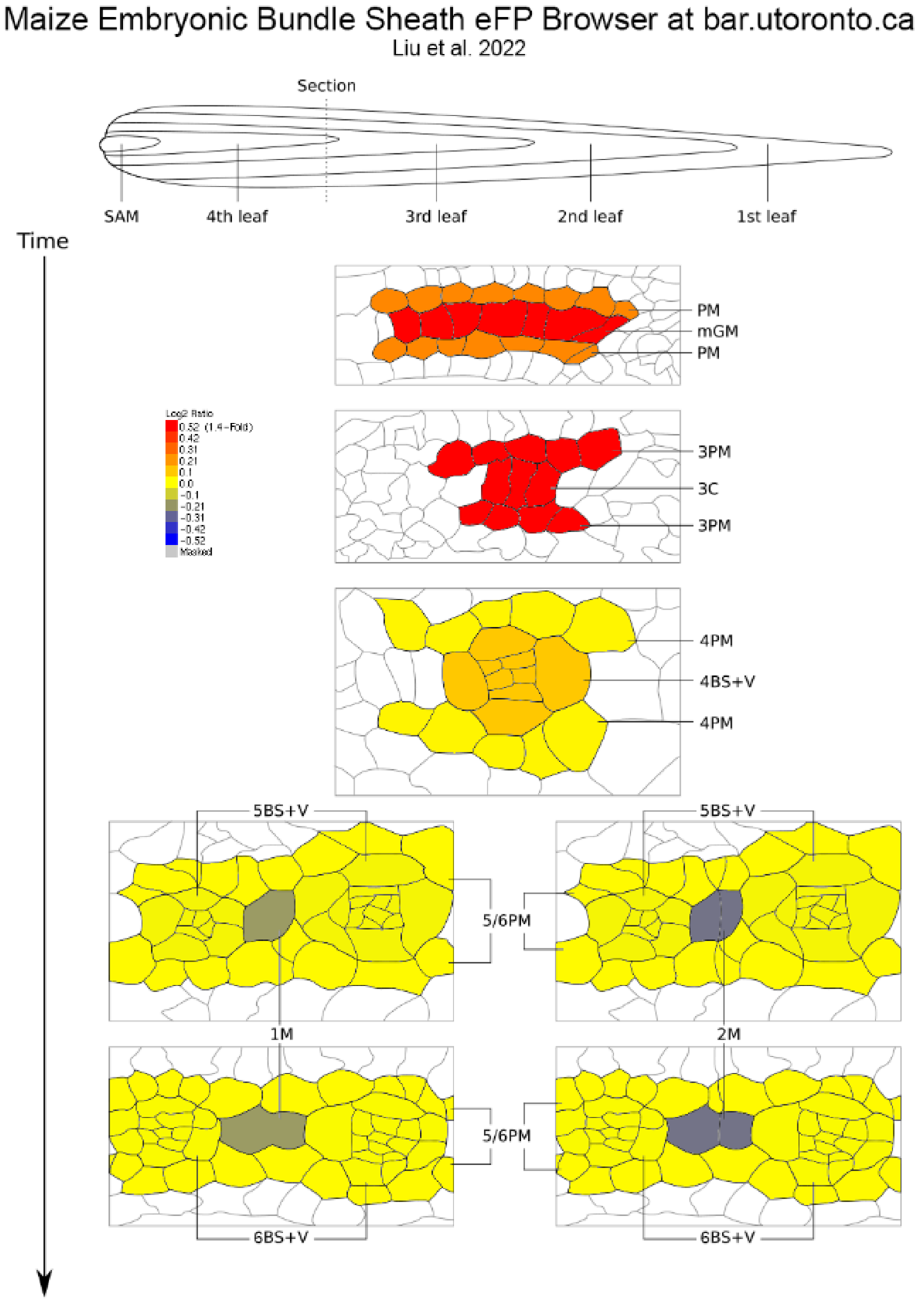

C. Maize gene atlas (Sekhon et al., 2011)- Absolute

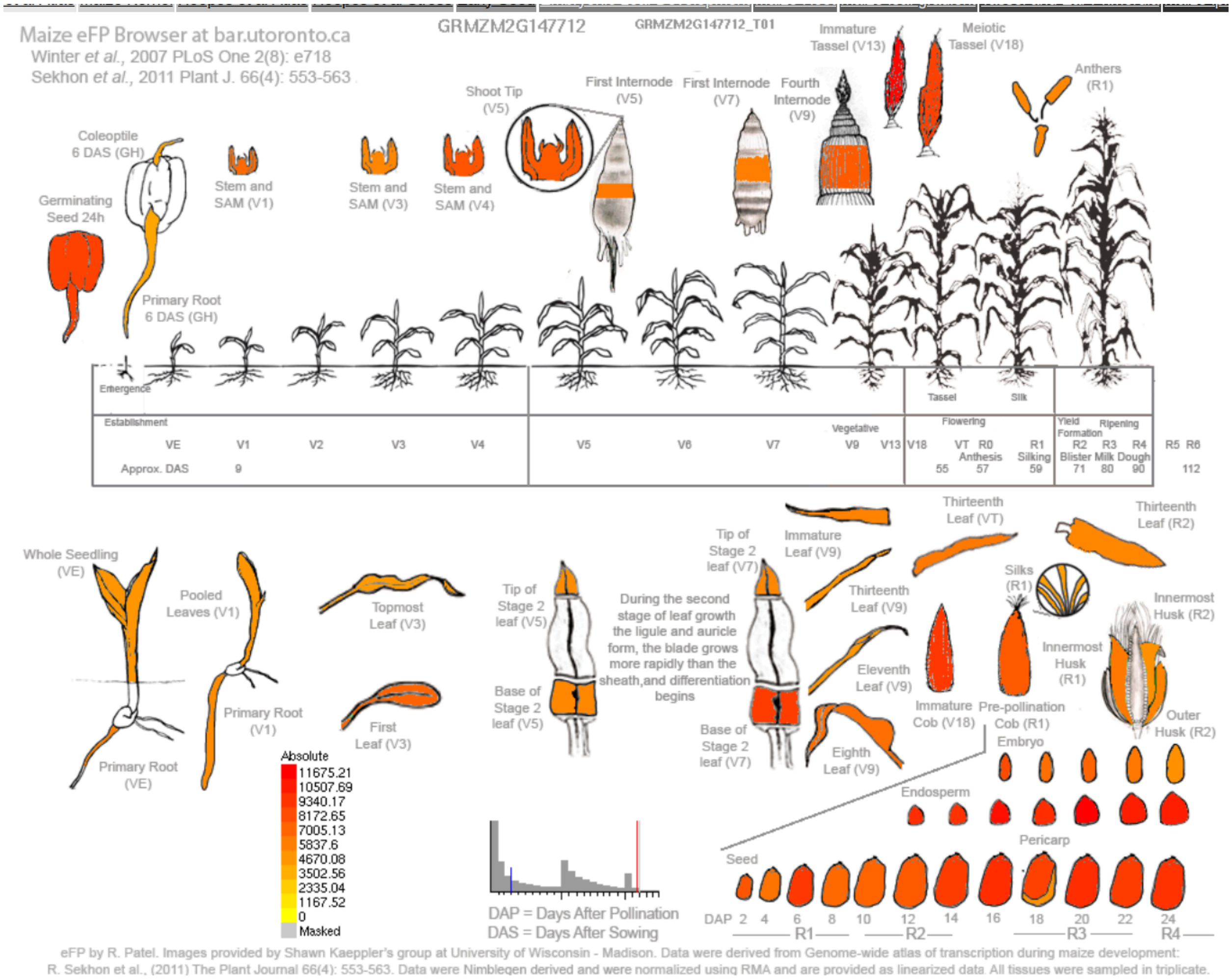

D. Maize gene atlas (Sekhon et al., 2011)- Relative

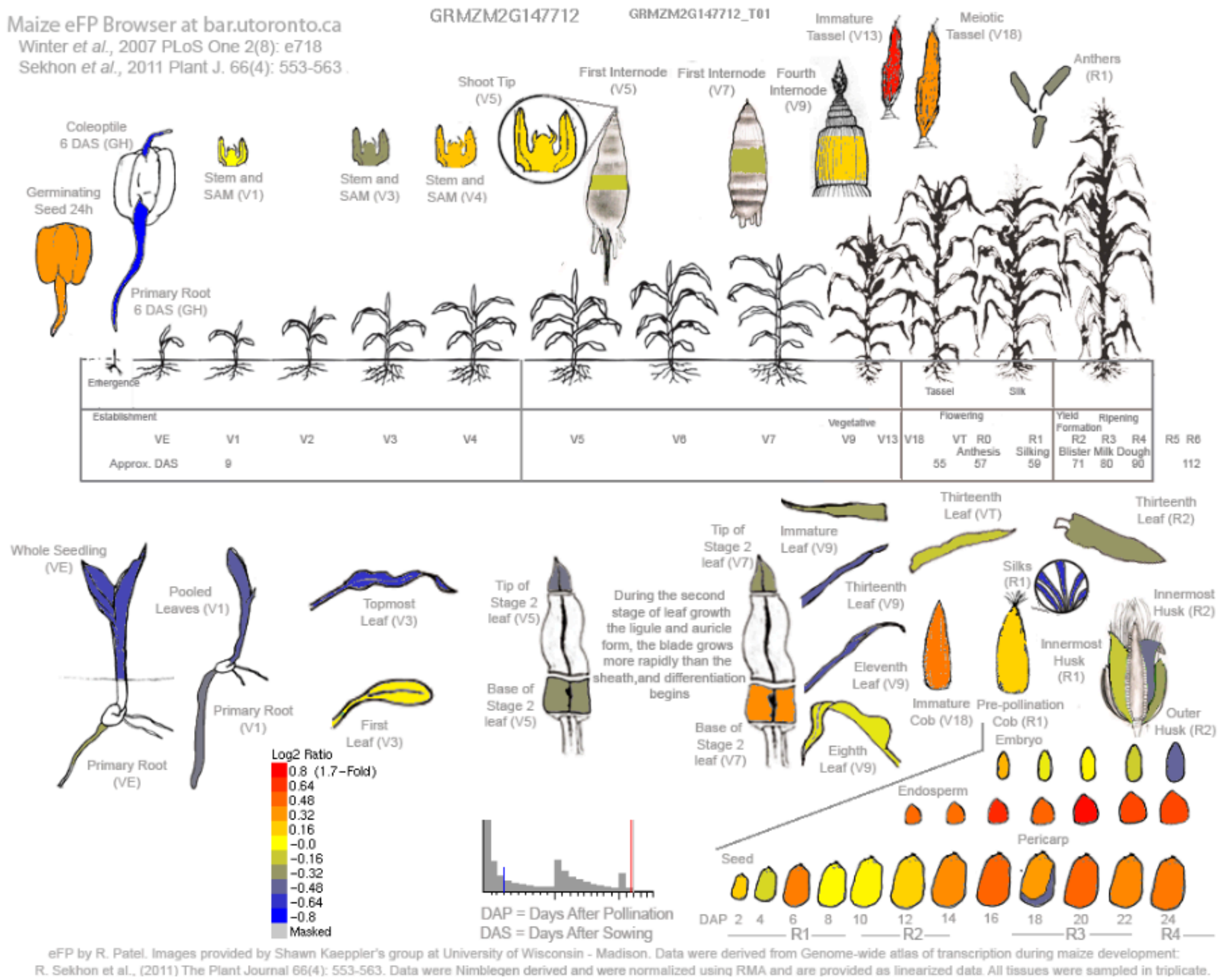

E. Maize leaf development gradient (Li et al., 2010)- Relative

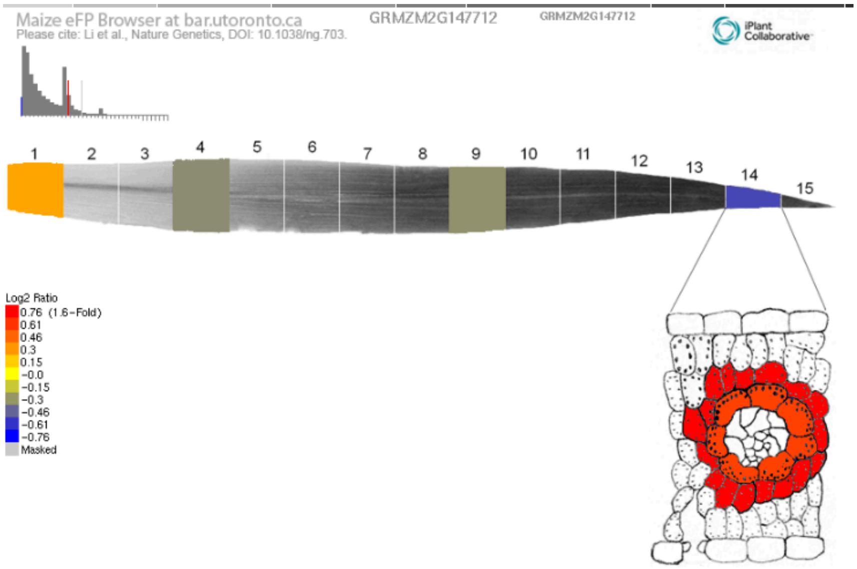

Supplementary Figure S8:

Publicly available gene expression atlas data of **Cadtfr3** (GRMZM2G147712 / Zm00001d050242). Maize embryonic leaf development (Liu et 2022)- (A) absolute and (B) relative, Maize gene atlas (Sekhon et al., 2011)- (C) absolute and (D) relative, (E) Maize leaf development gradient (Li et al., 2010)- relative.

A. Maize embryonic leaf development (Liu et 2022)- Absolute

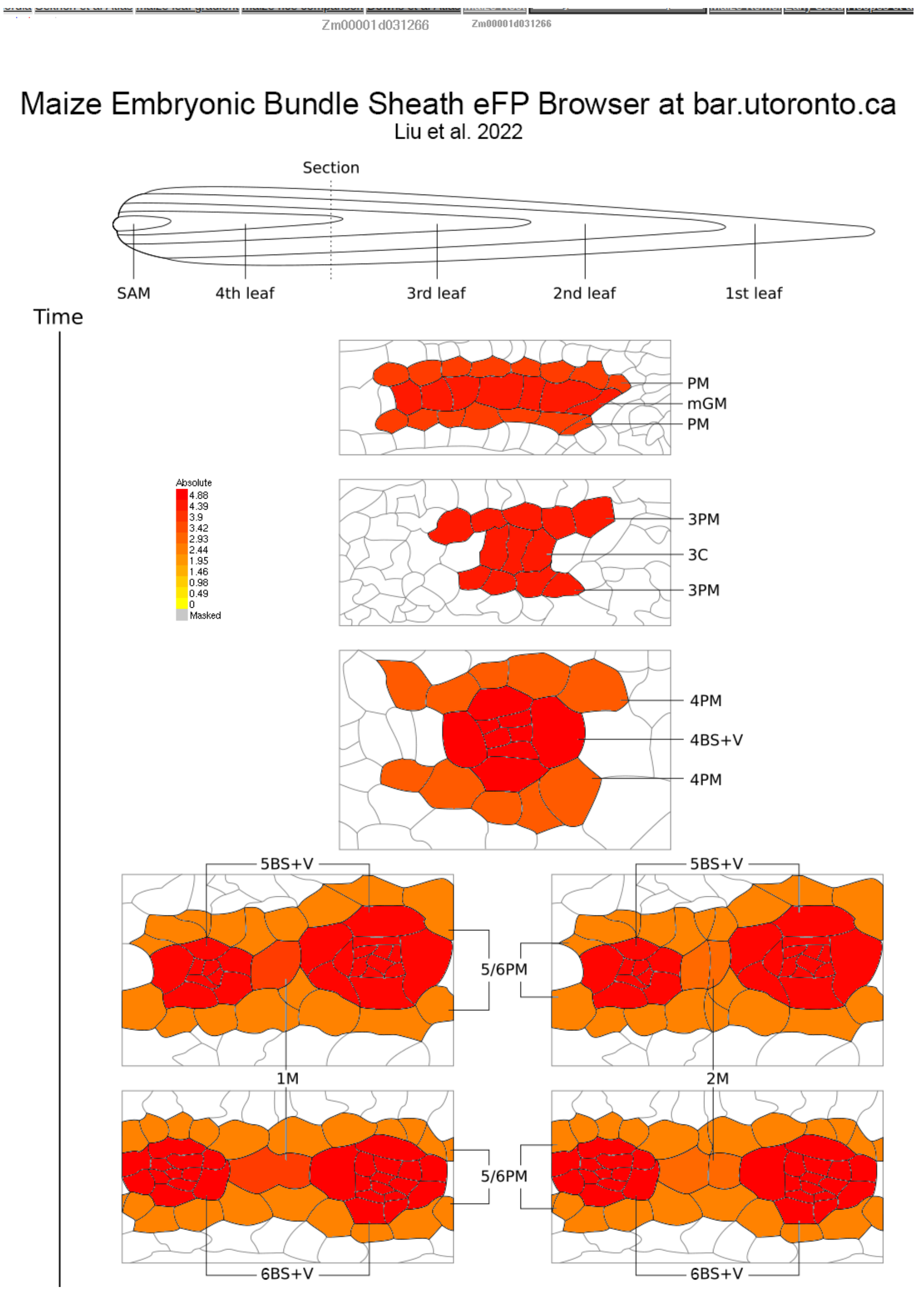

B. Maize embryonic leaf development (Liu et 2022)- Relative

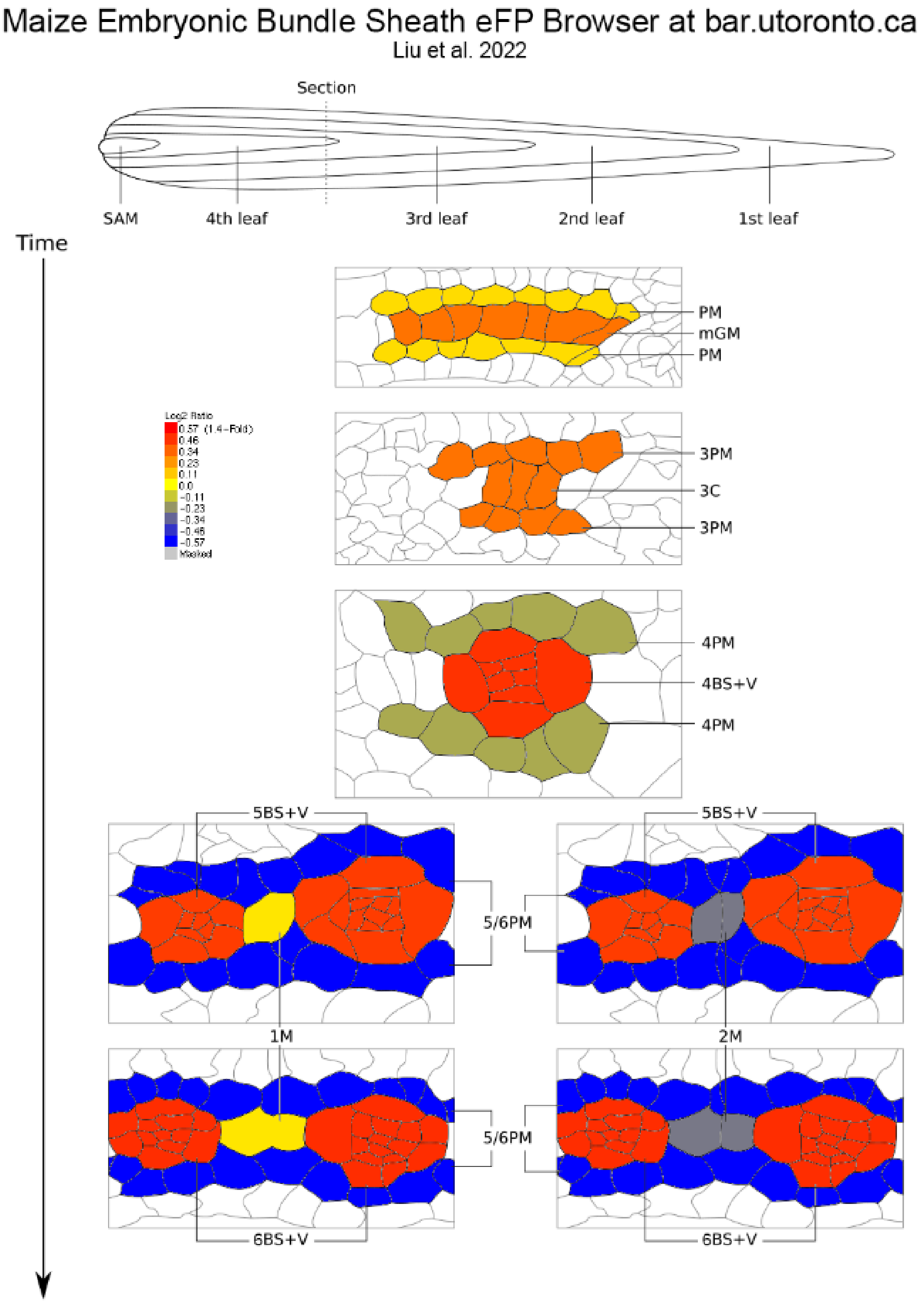

C. Maize gene atlas (Sekhon et al., 2011)- Absolute

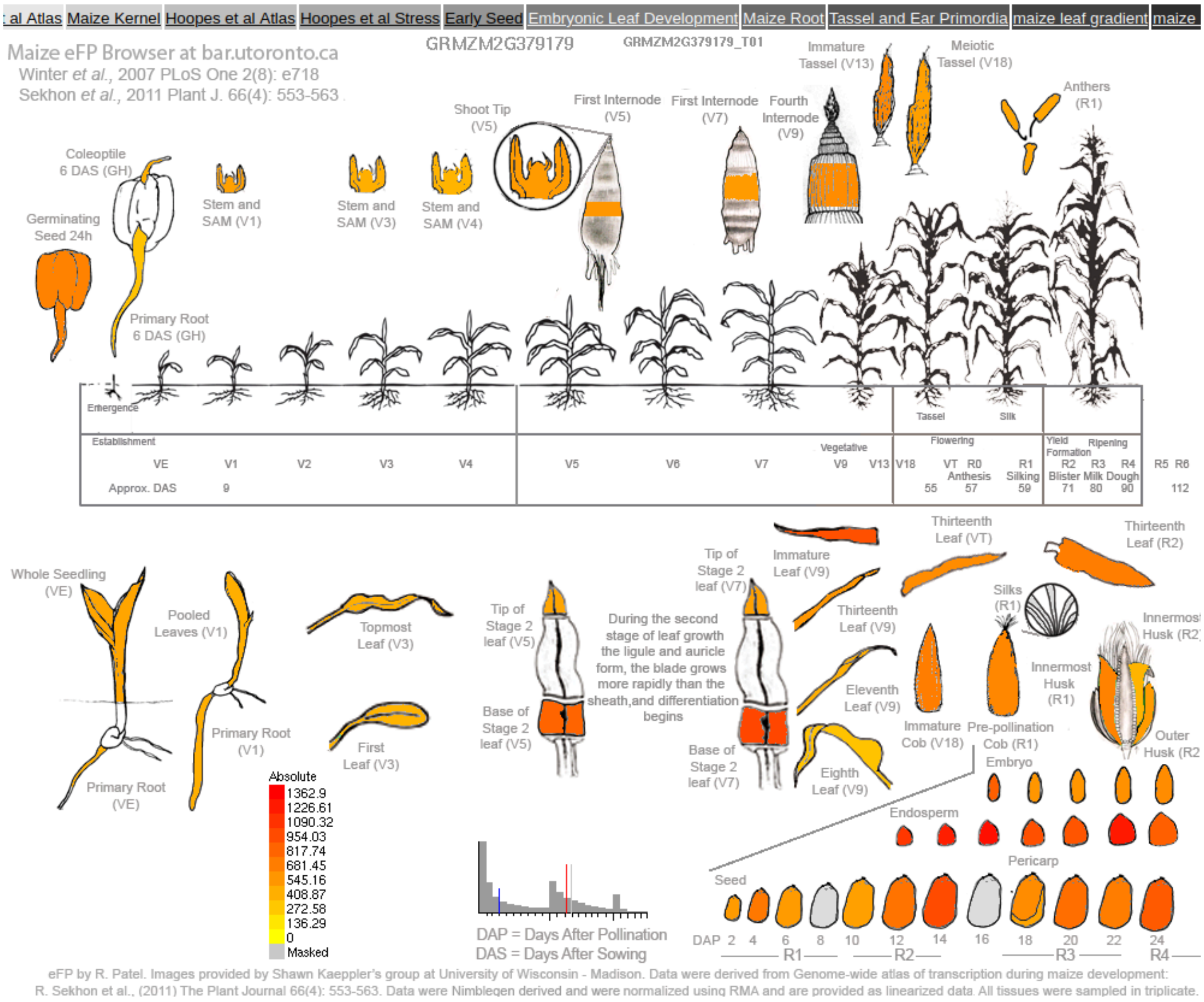

D. Maize gene atlas (Sekhon et al., 2011)- Relative

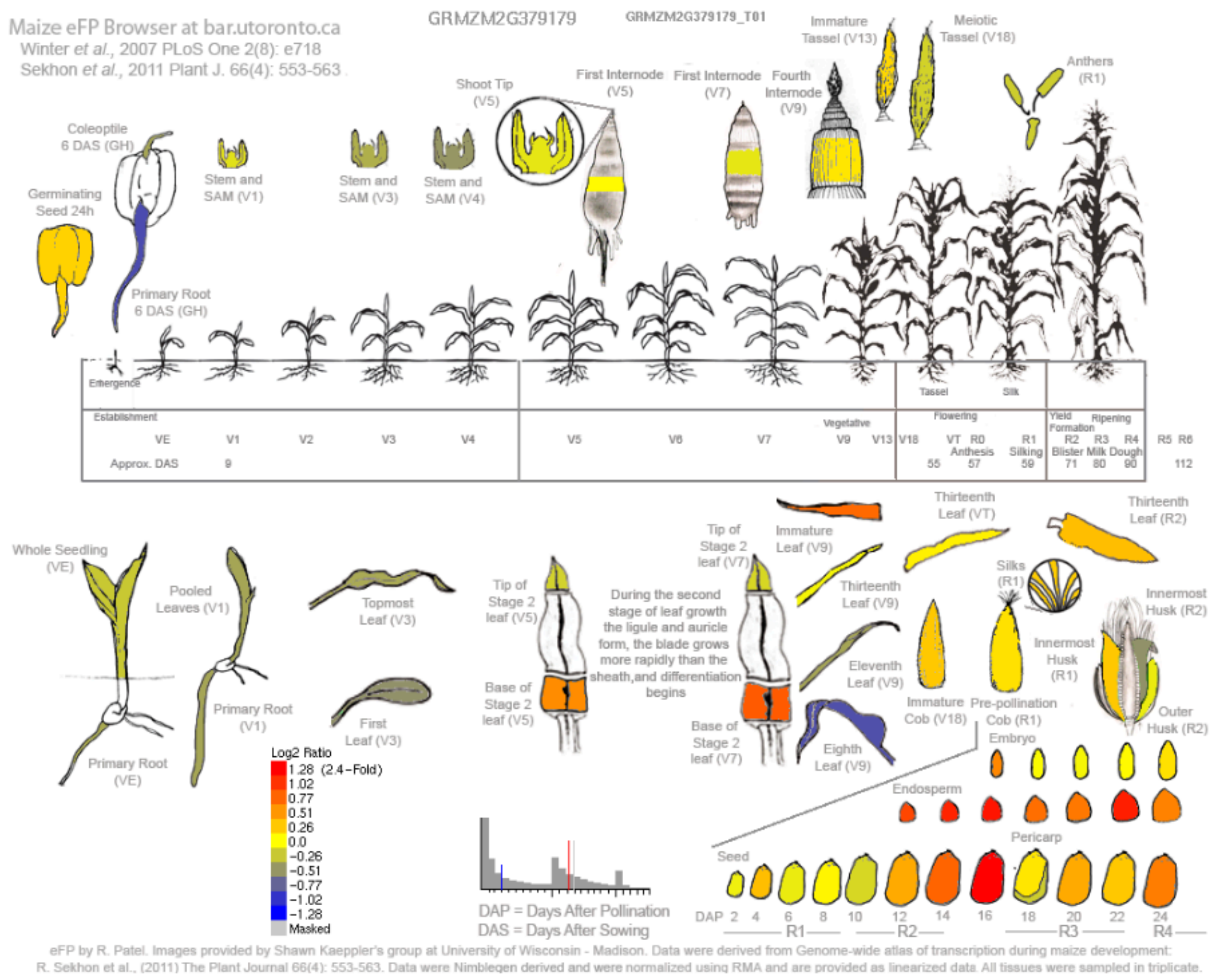

E. Maize leaf development gradient (Li et al., 2010)- Relative

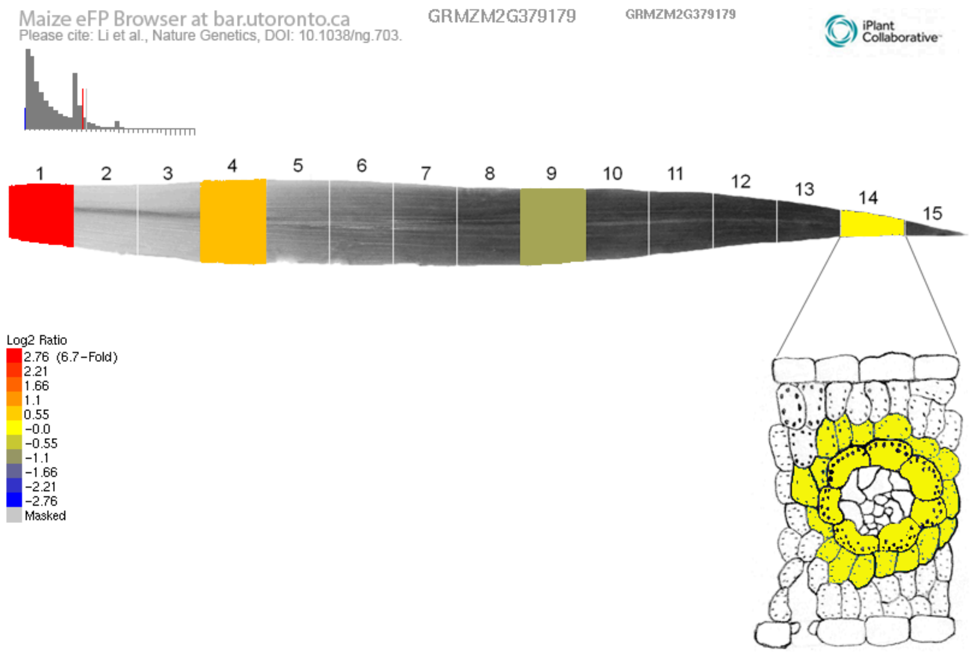

Supplementary Figure S9:

Publicly available gene expression atlas data of **Thx8** (GRMZM2G379179 / Zm00001d031266). Maize embryonic leaf development (Liu et 2022)- (A) absolute and (B) relative, Maize gene atlas (Sekhon et al., 2011)- (C) absolute and (D) relative, (E) Maize leaf development gradient (Li et al., 2010)- relative.

A. Maize embryonic leaf development (Liu et 2022)- Absolute

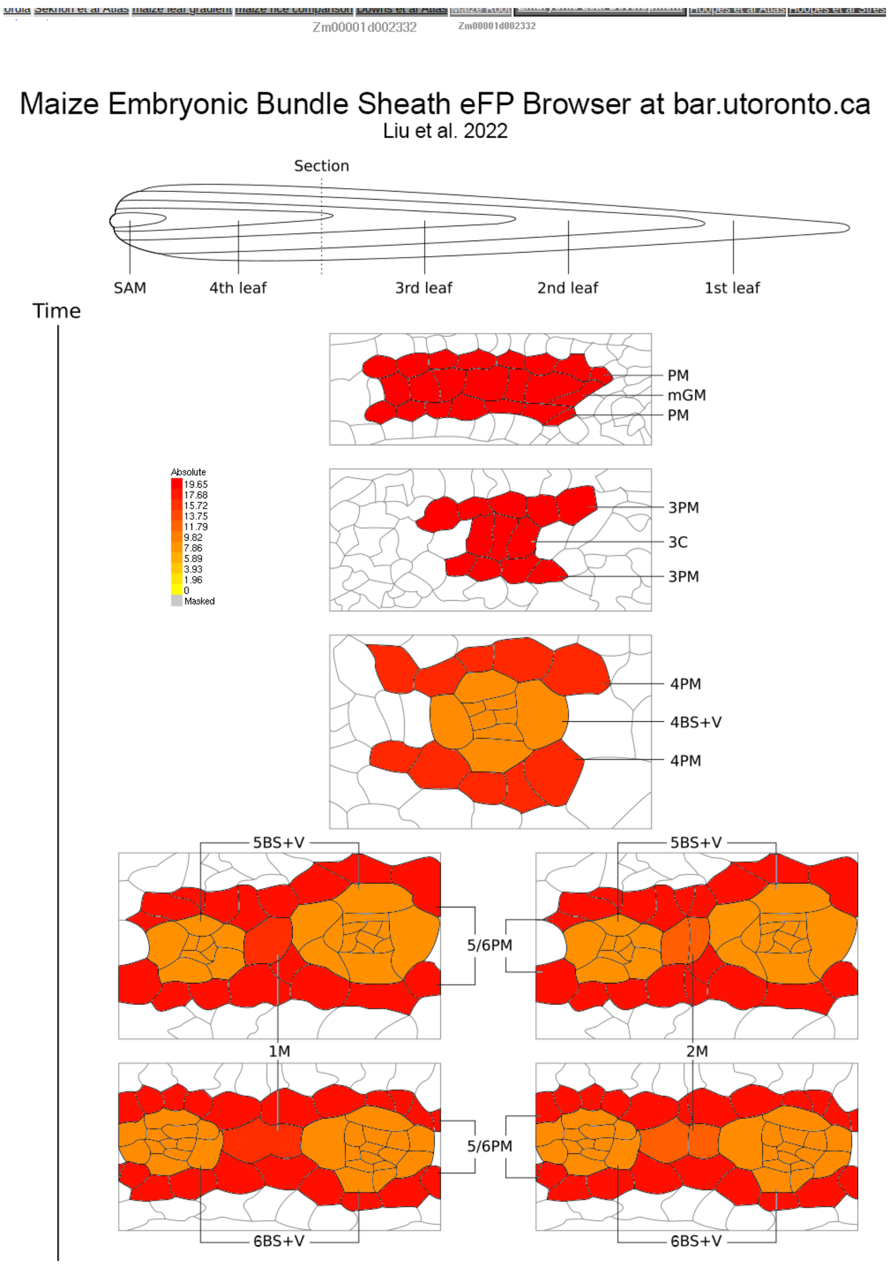

B. Maize embryonic leaf development (Liu et 2022)- Relative

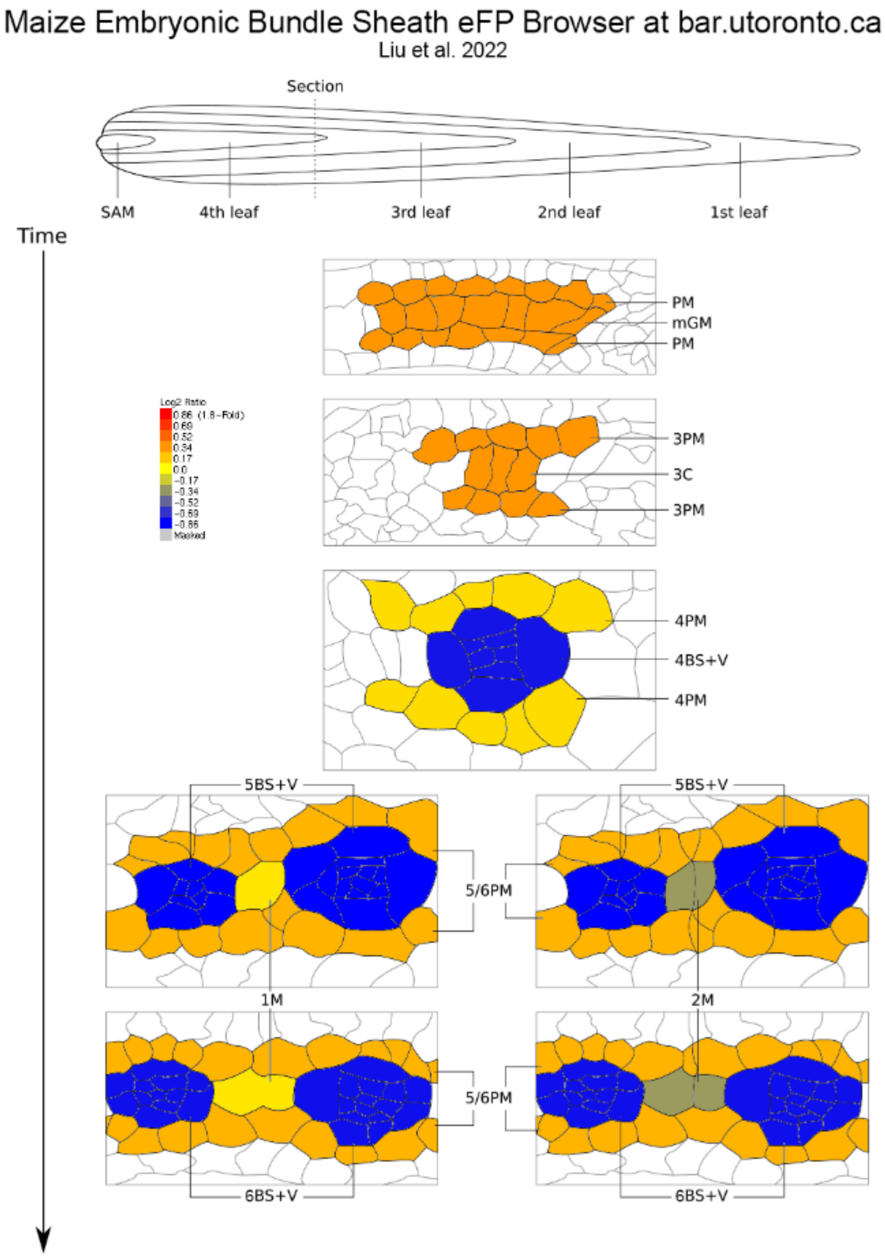

C. Maize gene atlas (Sekhon et al., 2011)- Absolute

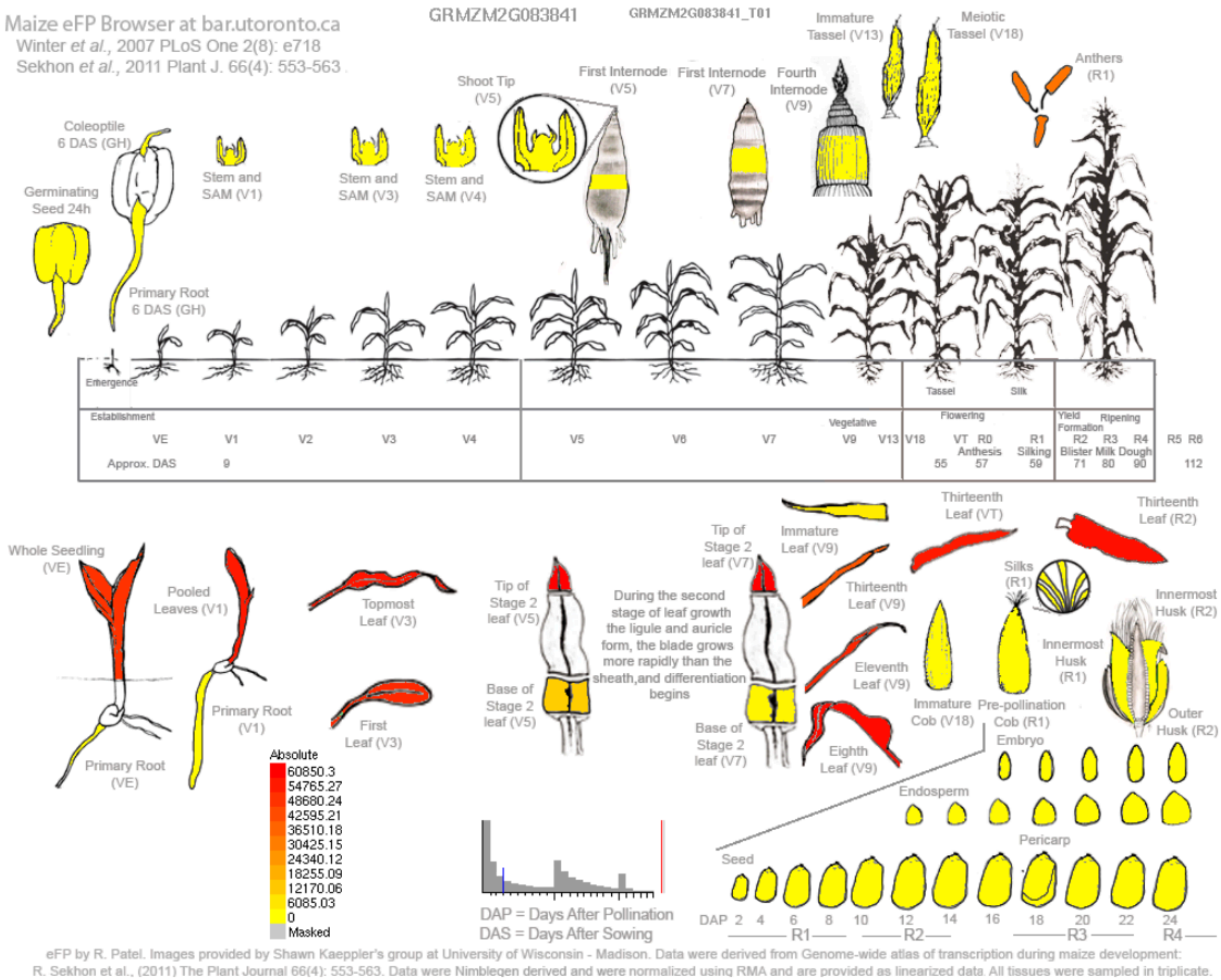

E. Maize gene atlas (Sekhon et al., 2011)- Relative

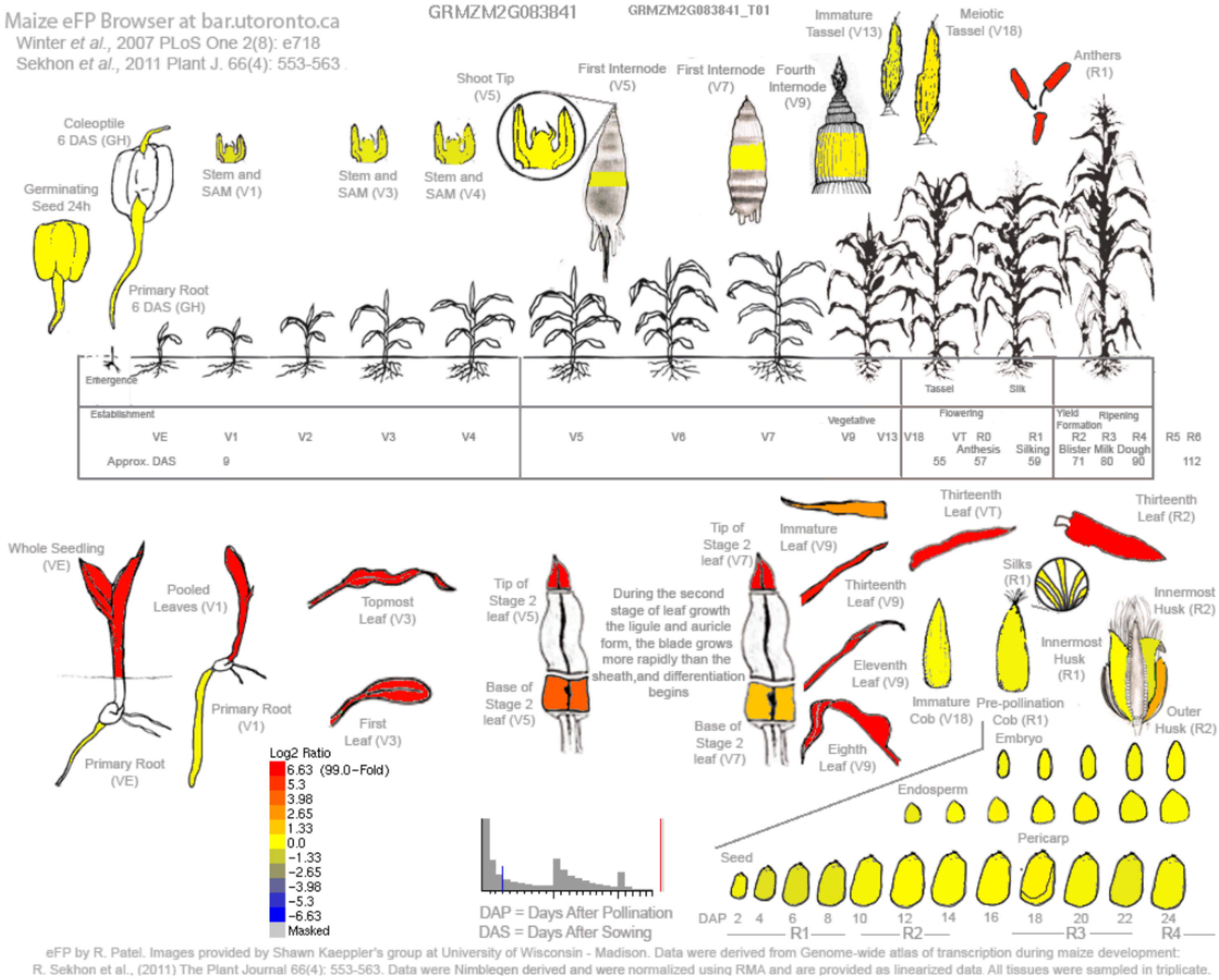

D. Maize leaf development gradient (Li et al., 2010)- Relative

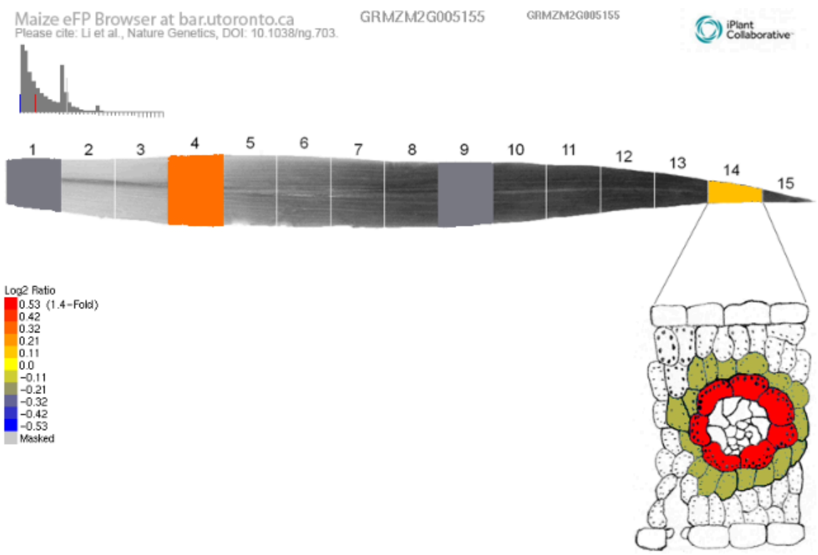

Supplementary Figure S10:

Publicly available gene expression atlas data of **Mads9** (GRMZM2G005155 / Zm00001d002332). Maize embryonic leaf development (Liu et 2022)- (A) absolute and (B) relative, Maize gene atlas (Hoopes et 2018)- (C) relative and (D) Maize leaf development gradient (Li et al., 2010)- relative.

A. Maize embryonic leaf development (Liu et 2022)- Absolute

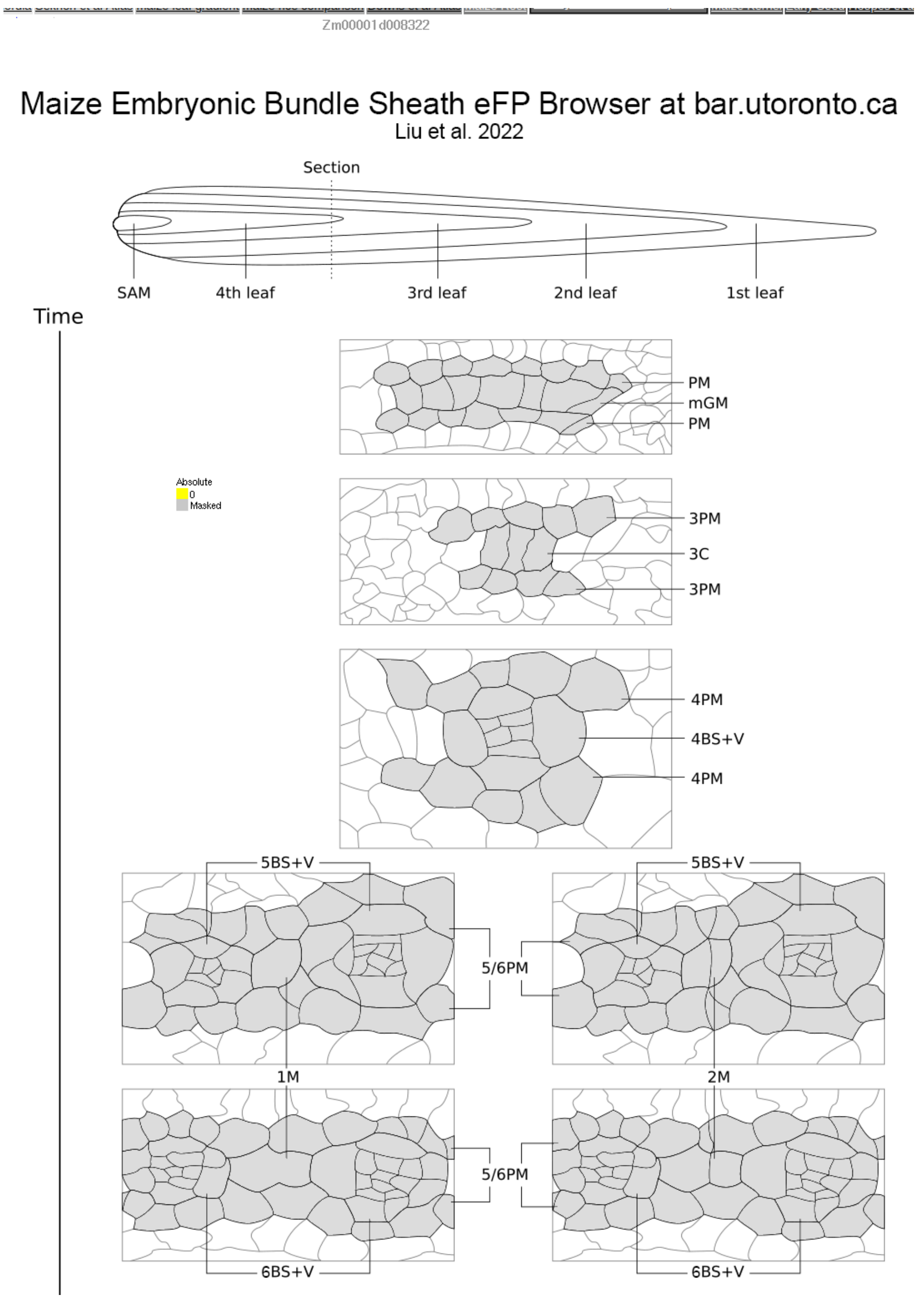

B. Maize embryonic leaf development (Liu et 2022)- Relative

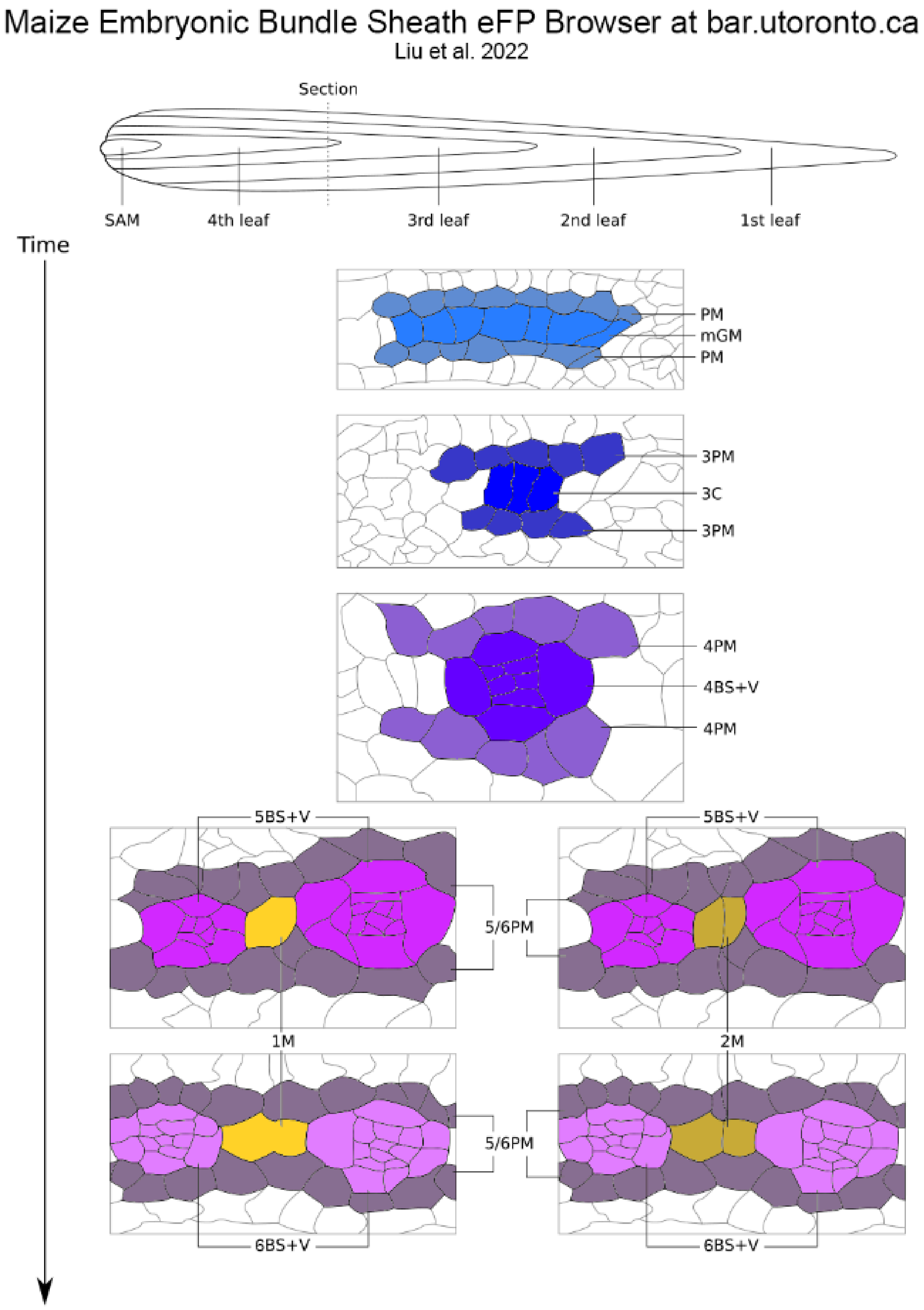

C. Maize gene atlas (Sekhon et 2011)- Absolute

- The requested Primary gene / probeset ID "Zm00001d008322" cannot be found in group1 datasource

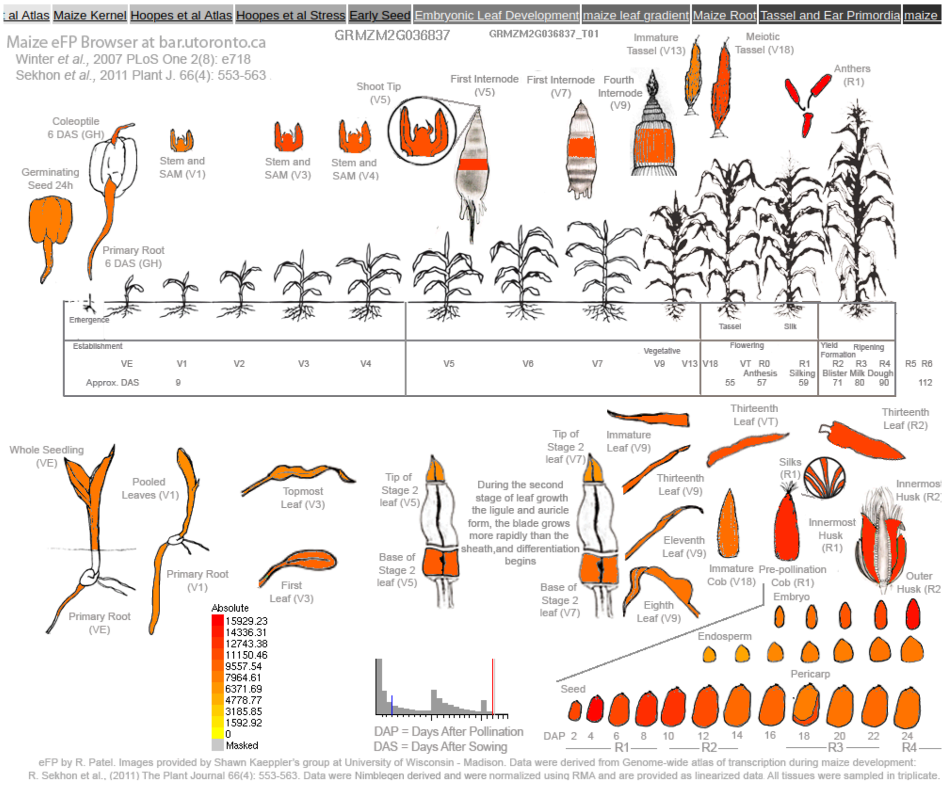

D. Maize gene atlas (Sekhon et 2011)- Relative

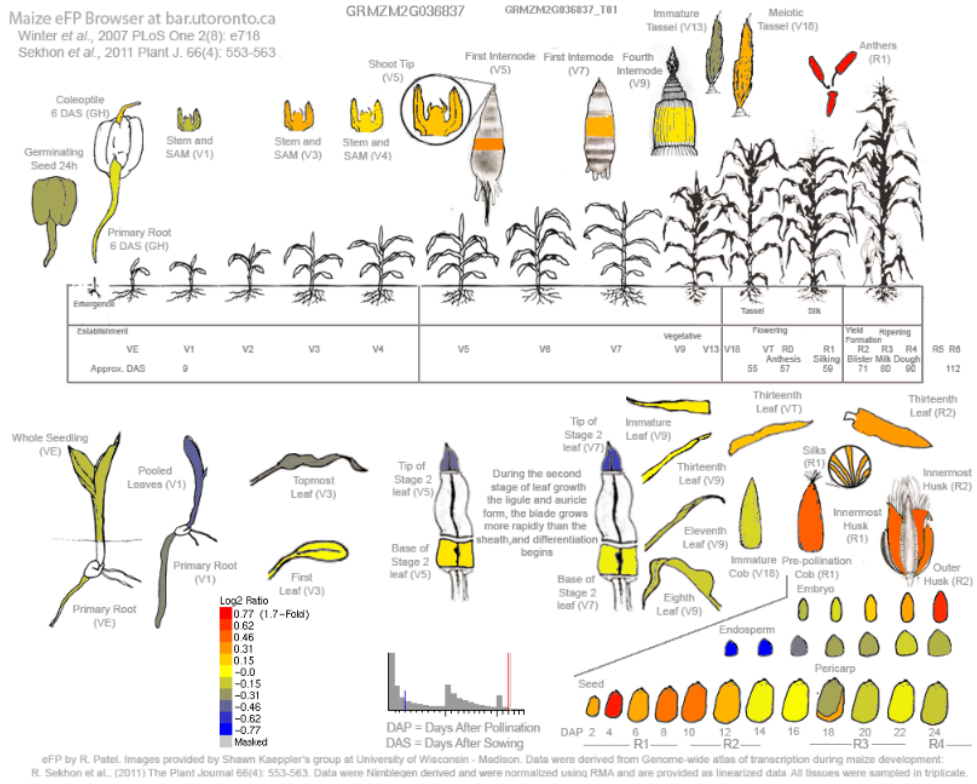

E. Maize leaf development gradient (Li et al., 2010)- Relative

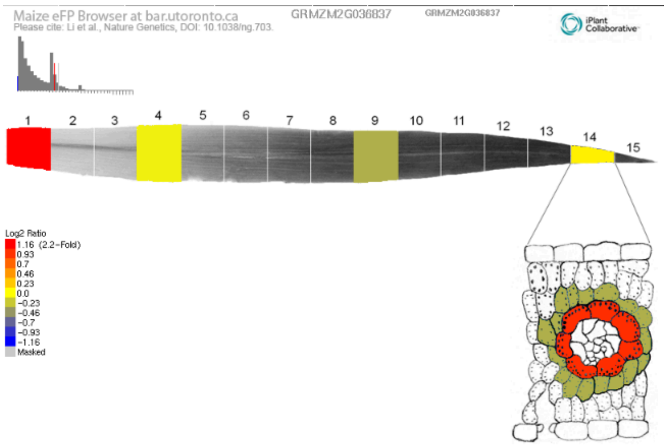

#### Supplementary Figure S11:

Publicly available gene expression atlas data of **C3H28** (GRMZM2G036837 / Zm00001d008322). Maize embryonic leaf development (Liu et 2022)- (A) absolute and (B) relative, Maize gene atlas (Sekhon et al., 2011)- (C) absolute and (D) relative, (E) Maize leaf development gradient (Li et al., 2010)- relative.

A. Maize embryonic leaf development (Liu et 2022)- Absolute

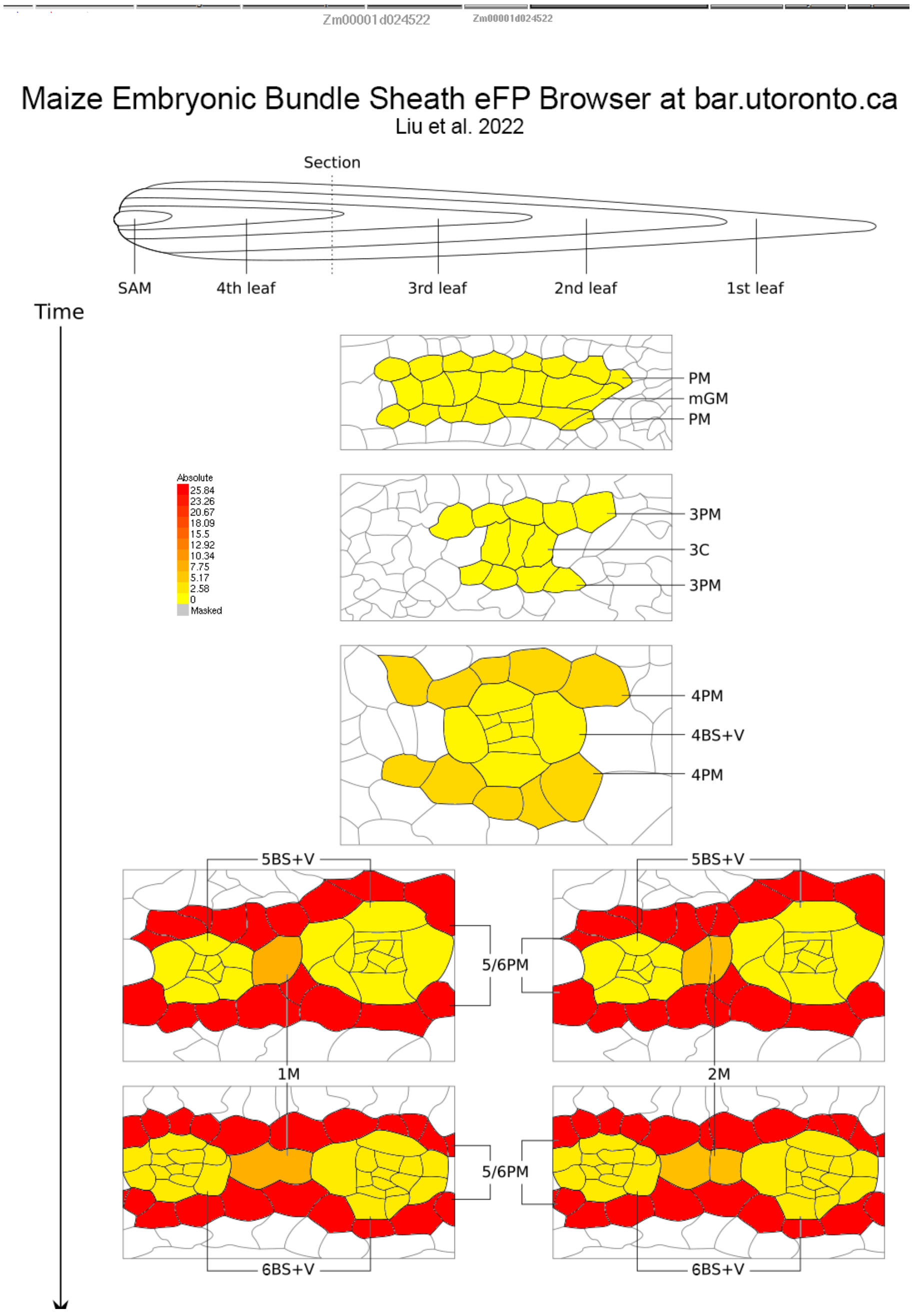

B. Maize embryonic leaf development (Liu et 2022)- Relative

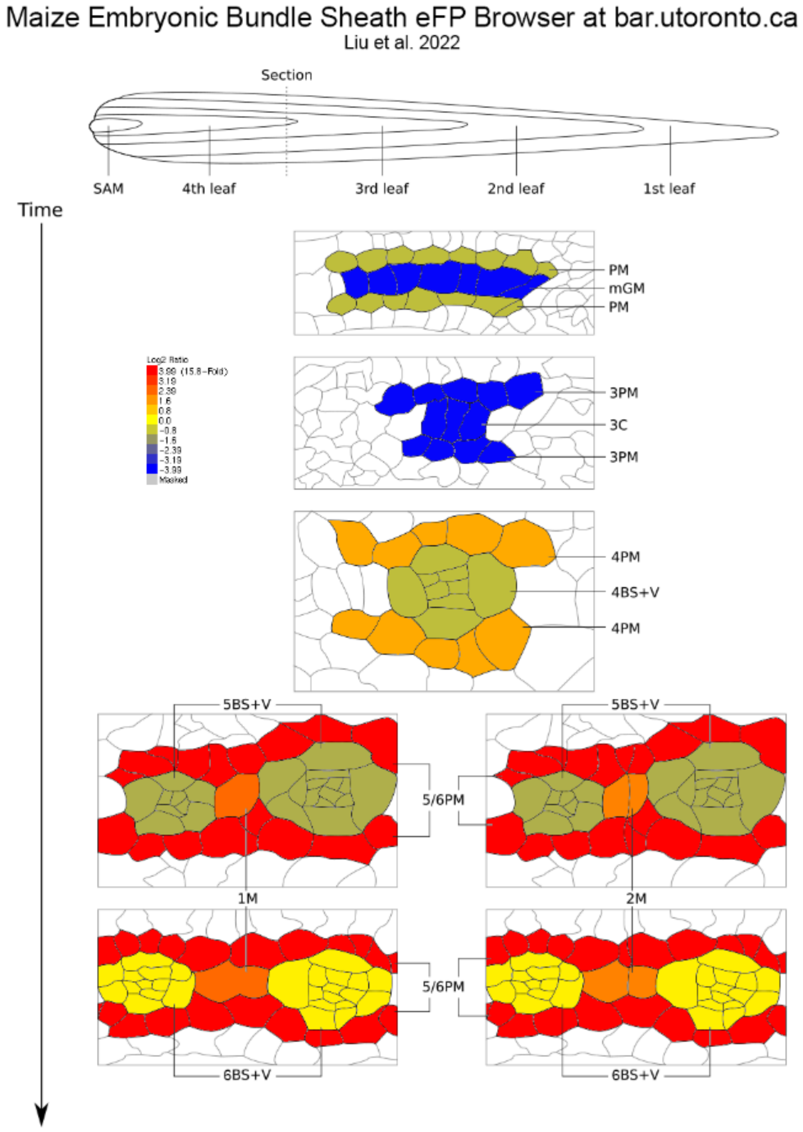

C. Maize gene atlas (Sekhon et al., 2011)- Absolute

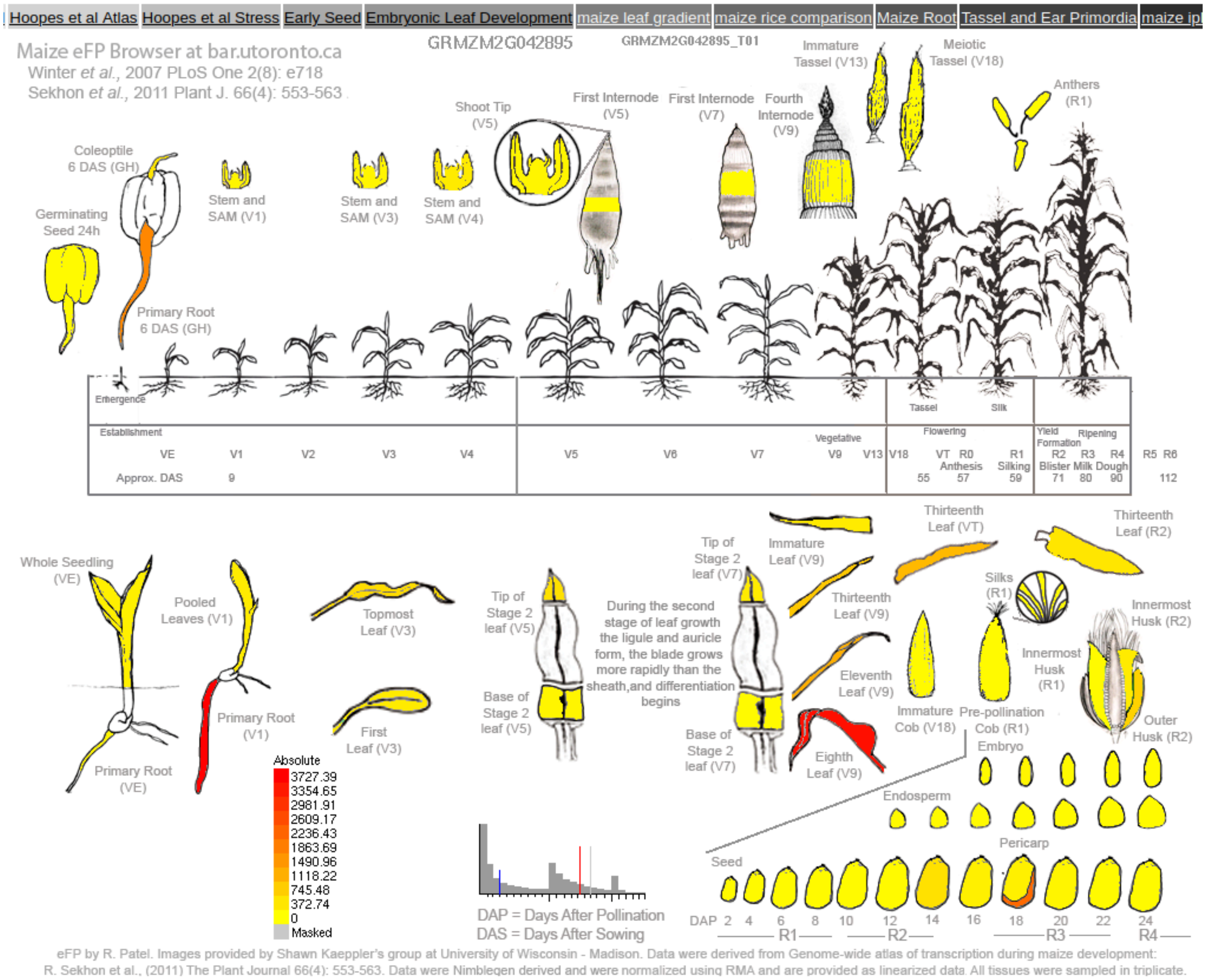

D. Maize gene atlas (Sekhon et al., 2011)- Relative

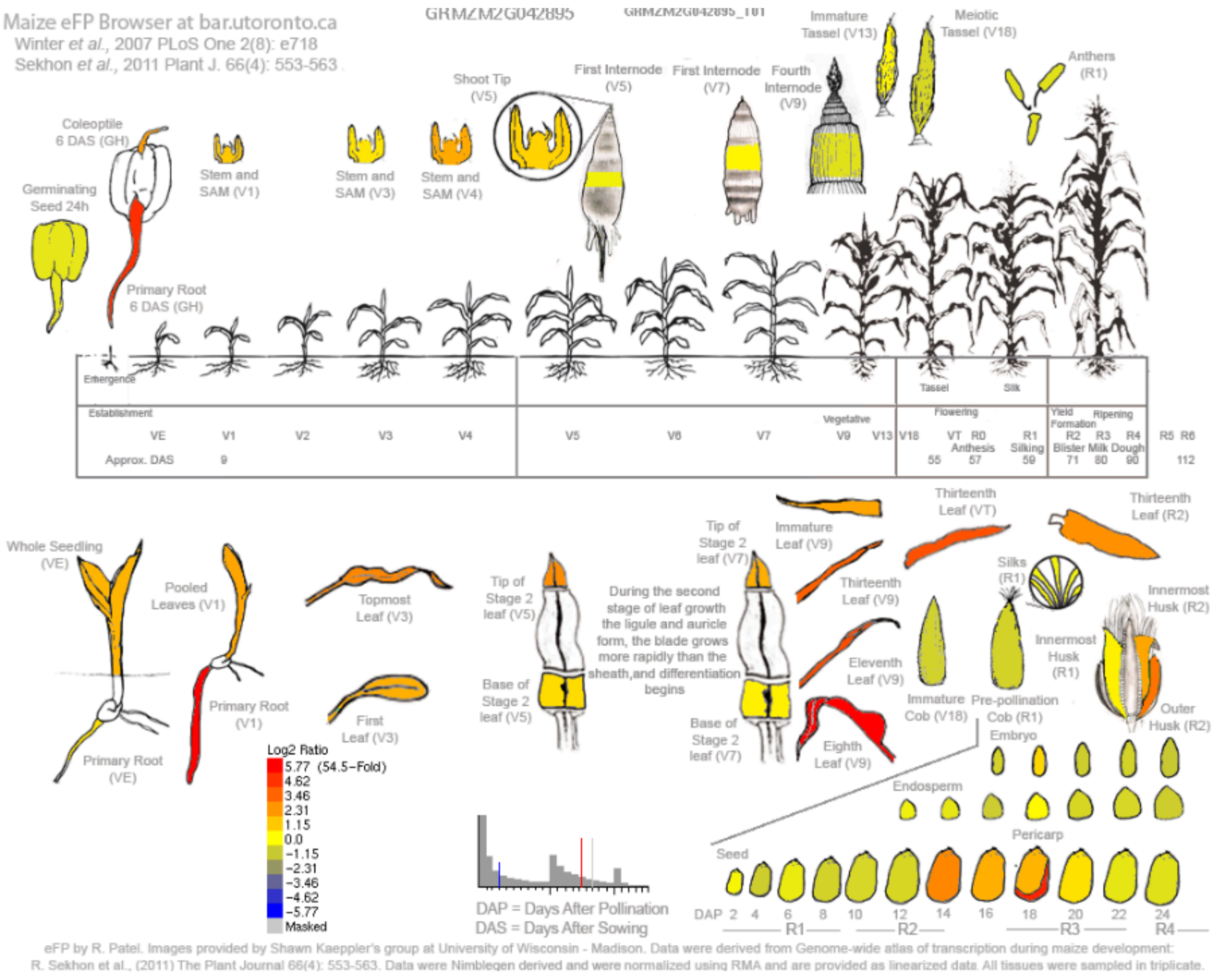

E. Maize leaf development gradient (Li et al., 2010)- Relative

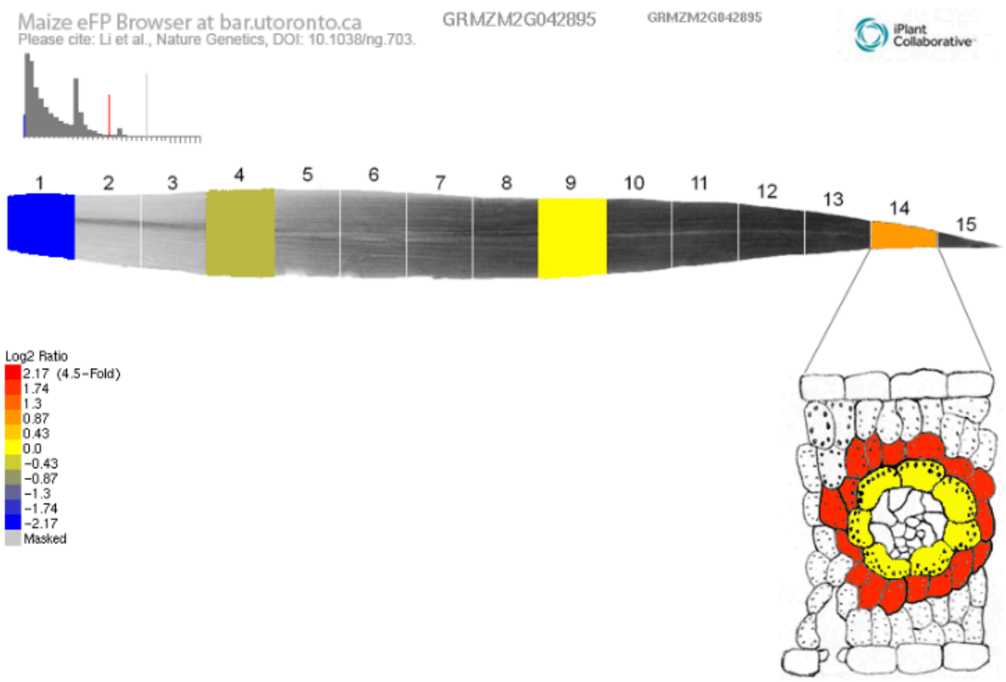

Supplementary Figure S12:

Publicly available gene expression atlas data of **BHLH116** (GRMZM2G042895 / Zm00001d024522). Maize embryonic leaf development (Liu et 2022)- (A) absolute and (B) relative, Maize gene atlas (Sekhon et al., 2011)- (C) absolute and (D) relative, (E) Maize leaf development gradient (Li et al., 2010)- relative.

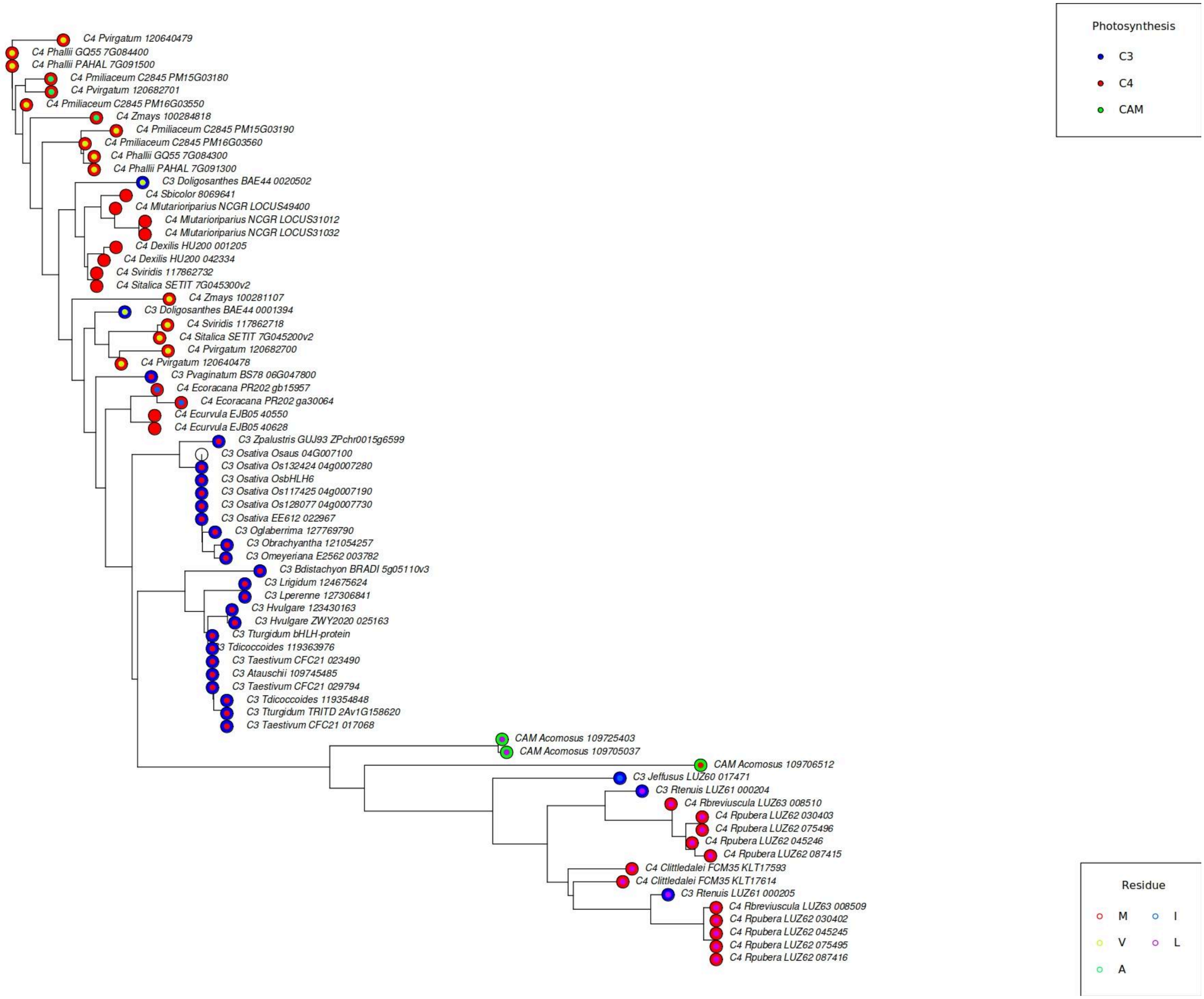

Supplementary Figure S13:

Extended gene family tree for BHLH116, showing changes in the position with $C_4$-specific changes across related $C_3$ and $C_4$ species. The outer circle at the end leaves indicates the photosynthetic type of the species ($C_3$, $C_4$ or CAM), and the inner circle indicates the residue present, whether Methionine (M), Valine(V), Alanine (A), Isoleucine (I), or Leucine(L). This is an unrooted tree made from aligning bHLH116 gene, it does not accurately represent the phylogenetic relations.

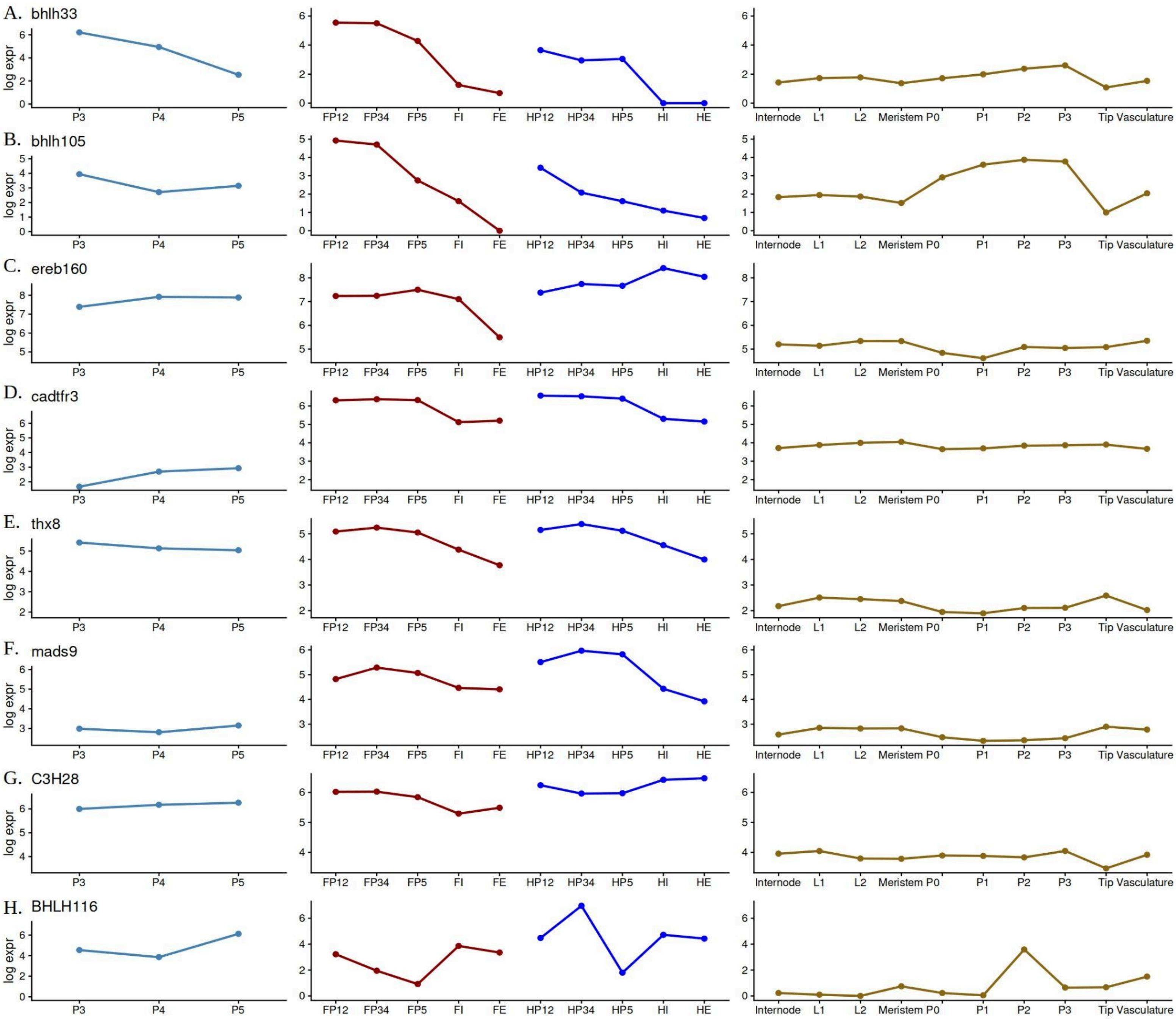

Supplementary Figure S14:

Comprehensive expression profiles in multiple RNAseq studies. The left panel shows the expression in rice leaf primordia (van-Campen et al. 2016), the middle panel shows expression of the gene in maize foliar and husk leaves (Wang et al., 2013), and the right panel shows the expression of gene in early leaf primordia (Knauer S, et al. 2019). A to H are the genes in the order of *bHLH33, bHLH105, EREB160, CADTFR3, THX8, MADS9, C3H28, bHLH116*.

.

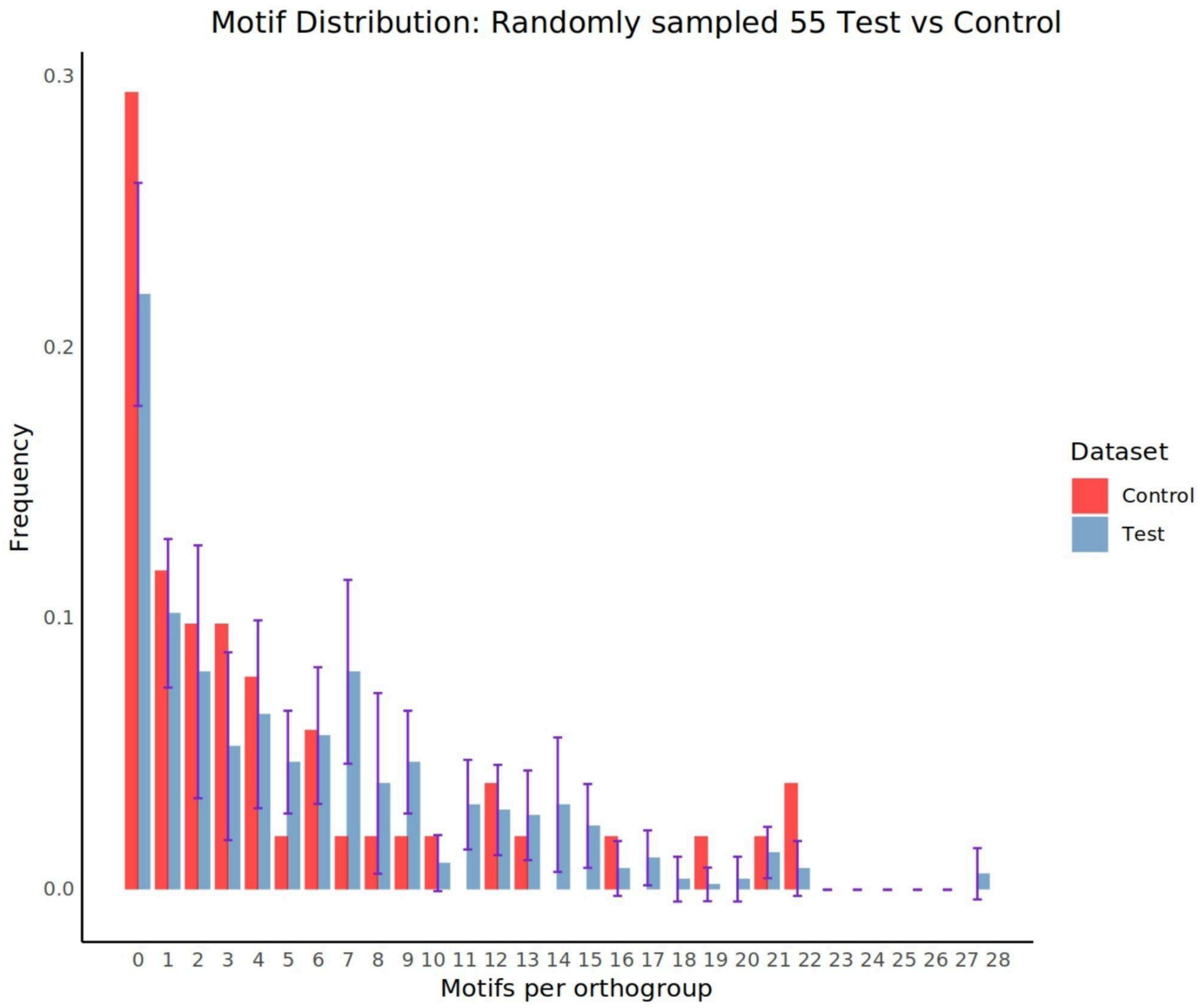

Supplementary Figure S15:

Mean frequency of motif distribution per orthogroup across 10 replicates of 55 randomly sampled test orthogroups, compared with the control dataset.

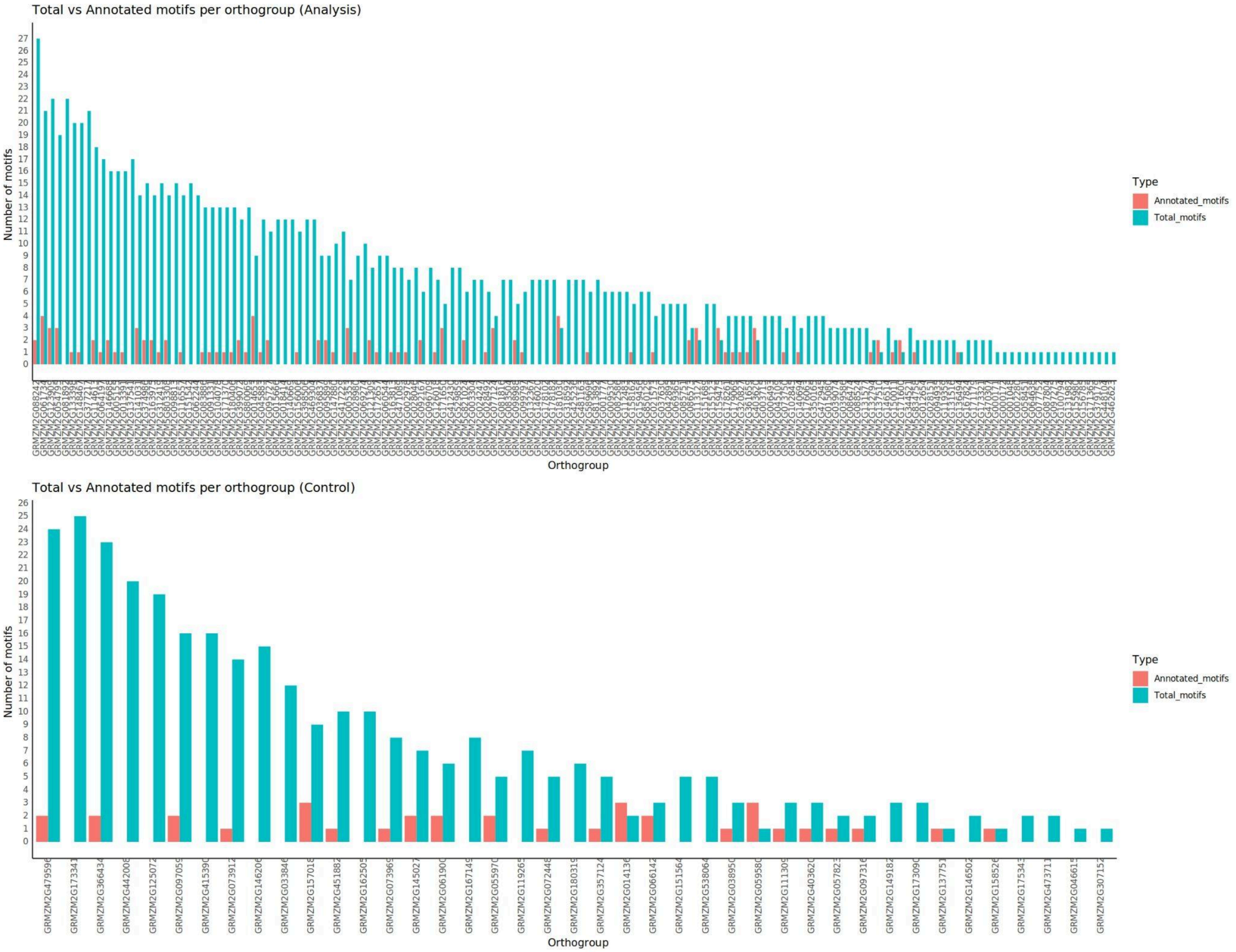

Supplementary Figure S16:

Total vs annotated motifs per orthogroups for test(Analysis) and control dataset.

### Supplementary Tables :

#### Supplementary Table S1:

**Genes selected from study 1 (Wang et al., 2013; Refer to tables S26 and S27 of this reference).** Under the columns labeled “Foliar profile” and “Husk profile”, the prefixes “F” and “H” denote foliar and husk tissues, respectively, across developmental progression from primordium (P1) to P3/4 to P5. In both tissues, A-type profiles (FA, HA) show increasing trends—either continuous (A1), early increase followed by stabilization (A2), or late-stage increase (A3)—while D-type profiles (FD, HD) show corresponding decreasing patterns across these stages (D1–D3). FN and HN represent genes or TFs with no clear developmental trend (neutral) in foliar and husk tissues, respectively. Genes without complete orthogroups are highlighted in red.

| Gene ID | Gene ID (Gramene database) | EntrezGene | Profile Evidence | Present in Study 2? | Foliar profile | Husk profile |
|---|---|---|---|---|---|---|
| GRMZM2G151542 | Zm00001d011499 | hypothetical protein LOC100193610 | Positive Regulator | NO | FA1 | HN |
| GRMZM2G471089 | Zm00001d039434 | NA | Positive Regulator | NO | FA1 | NA |
| GRMZM2G039074 | Zm00001d033310 | hypothetical protein LOC100216818 | Positive Regulator | NO | FA2 | HN |
| GRMZM2G045883 | Zm00001d018119 | hypothetical protein LOC100217195 | Positive Regulator | NO | FA2 | HN |
| GRMZM2G098988 | Zm00001d046332 | hypothetical protein LOC100217078 | Positive Regulator | NO | FA2 | HN |
| GRMZM2G123900 | Zm00001d030727 | hypothetical protein LOC100273972 | Positive Regulator | NO | FA2 | HN |
| GRMZM2G132794 | Zm00001d029607 | NA | Positive Regulator | NO | FA2 | HN |
| GRMZM2G136494 | Zm00001d026684 | NA | Positive Regulator | NO | FA2 | HN |
| GRMZM2G163975 | Zm00001d040186 | NA | Positive Regulator | NO | FA2 | HN |
| GRMZM2G172657 | Zm00001d021973 | hypothetical protein LOC100383480 | Positive Regulator | NO | FA2 | HN |
| GRMZM2G377217 | Zm00001d051328 | NA | Positive Regulator | NO | FA2 | HN |
| GRMZM5G893117 | Zm00001d018260 | NA | Positive Regulator | NO | FA2 | HN |
| GRMZM2G028046 | Zm00001d053006 | hypothetical protein LOC100191534 | Positive Regulator | YES | FA2 | HN |
| GRMZM2G150011 | Zm00001d020037 | zinc finger, C2H2 type family protein | Positive Regulator | YES | FA2 | HN |
| GRMZM2G399072 | Zm00001d027878 | NA | Positive Regulator | YES | FD3 | HD2 |
| GRMZM2G146688 | Zm00001d007840 | NA | Positive Regulator | YES | FD3 | HD3 |
| GRMZM2G040924 | Zm00001d017385 | P-type R2R3 Myb protein | Positive Regulator | NO | FD3 | HN |
| GRMZM2G069365 | Zm00001d051573 | zinc finger | Positive Regulator | NO | FD3 | HN |

| | | | | | | |
|---|---|---|---|---|---|---|
| | | homeodomain protein 1 | | | | |
| GRMZM2G095899 | Zm00001d033407 | hypothetical protein LOC100272590 | Positive Regulator | NO | FD3 | HN |
| GRMZM2G097275 | Zm00001d015233 | hypothetical protein LOC100279234 | Positive Regulator | NO | FD3 | HN |
| GRMZM2G098813 | Zm00001d026231 | zea floricaula/leafy1 | Positive Regulator | NO | FD3 | HN |
| GRMZM2G111045 | Zm00001d051269 | hypothetical protein LOC100272429 | Positive Regulator | NO | FD3 | HN |
| GRMZM2G121309 | Zm00001d016277 | IAA7 - auxin-responsive Aux/IAA family member | Positive Regulator | NO | FD3 | HN |
| GRMZM2G126018 | Zm00001d006028 | hypothetical protein LOC100217104 | Positive Regulator | NO | FD3 | HN |
| GRMZM2G318592 | Zm00001d036711 | hypothetical protein LOC100277679 | Positive Regulator | NO | FD3 | HN |
| GRMZM2G015666 | Zm00001d005939 | DNA-binding protein | Positive Regulator | YES | FD3 | HN |
| GRMZM2G021573 | Zm00001d048004 | hypothetical protein LOC100384380 | Positive Regulator | YES | FD3 | HN |
| GRMZM2G082586 | Zm00001d020790 | hypothetical protein LOC100280340 | Positive Regulator | YES | FD3 | HN |
| GRMZM2G312419 | Zm00001d008251 | NA | Positive Regulator | NO | FD3 | NA |
| GRMZM2G002280 | Zm00001d007204 | LOC100191204 | Positive Regulator | NO | FN | HA3 |
| GRMZM2G171365 | Zm00001d048474 | MADS1, MADS1 | Positive Regulator | NO | FN | HD2 |
| AC215201.3_FG008 | - | NA | Positive Regulator | NO | FN | HN |
| GRMZM2G061734 | Zm00001d006091 | hypothetical protein LOC100279240 | Positive Regulator | NO | FN | HN |
| GRMZM2G131516 | Zm00001d052380 | protein SCARECROW | Positive Regulator | NO | FN | HN |
| GRMZM2G131577 | Zm00001d032668 | LOC100194356 | Positive Regulator | NO | FN | HN |
| GRMZM2G140669 | Zm00001d017409 | GATA zinc finger family protein | Positive Regulator | NO | FN | HN |
| GRMZM2G148467 | Zm00001d026175 | NA | Positive Regulator | NO | FN | HN |
| GRMZM2G178182 | Zm00001d006110 | hypothetical protein LOC100194274, bHLH transcription factor GBOF-1 | Positive Regulator | NO | FN | HN |
| GRMZM2G425236 | Zm00001d013409 | ZF-HD protein dimerisation region containing protein | Positive Regulator | NO | FN | HN |
| GRMZM2G462623 | Zm00001d011597 | NA | Positive Regulator | NO | FN | HN |
| GRMZM2G469304 | Zm00001d028284 | ternary complex factor MIP1 | Positive Regulator | NO | FN | HN |
| GRMZM2G472945 | Zm00001d015485 | NA | Positive Regulator | NO | FN | HN |
| GRMZM5G850129 | Zm00001d037117 | growth-regulating factor 7-like | Positive Regulator | NO | FN | HN |

| | | | | | | |
|---|---|---|---|---|---|---|
| GRMZM2G178102 | Zm00001d041489 | hypothetical protein LOC100274567 | Positive Regulator | YES | FN | HN |
| GRMZM2G119359 | Zm00001d045533 | NA | Positive Regulator | NO | NA | HN |
| GRMZM2G374986 | Zm00001d048448 | NA | Positive Regulator | NO | NA | HN |
| GRMZM2G417229 | Zm00001d017784 | NA | Positive Regulator | NO | NA | HN |
| GRMZM5G887276 | Zm00001d007339 | hypothetical protein LOC100275291 | Positive Regulator | NO | NA | HN |
| GRMZM2G045109 | Zm00001d046096 | NA | Negative Regulator | NO | FA1 | HN |
| GRMZM2G064638 | Zm00001d018119 | hypothetical protein LOC100381548 | Negative Regulator | NO | FA1 | HN |
| GRMZM2G086573 | Zm00001d002025 | hypothetical protein LOC100384122 | Negative Regulator | NO | FA1 | HN |
| GRMZM2G000842 | Zm00001d038296 | NA | Negative Regulator | NO | FA2 | HA2 |
| GRMZM2G137541 | Zm00001d005964 | DNA binding protein, hypothetical protein LOC100383015 | Negative Regulator | NO | FA2 | HA2 |
| GRMZM2G176063 | Zm00001d041549 | NA | Negative Regulator | NO | FA2 | HA2 |
| GRMZM2G140694 | Zm00001d017788 | hypothetical protein LOC100216759 | Negative Regulator | NO | FA2 | HN |
| GRMZM2G085751 | Zm00001d053391 | hypothetical protein LOC100194372, DNA binding protein | Negative Regulator | NO | FA3 | HN |
| GRMZM2G180406 | Zm00001d042373 | LOC100278938, hypothetical protein LOC100382035 | Negative Regulator | NO | FA3 | HN |
| GRMZM2G028980 | Zm00001d053819 | NA | Negative Regulator | YES | FA3 | HN |
| GRMZM2G003715 | Zm00001d024543 | hypothetical protein LOC100383037 | Negative Regulator | NO | FN | HA2 |
| GRMZM2G060544 | Zm00001d029506 | hypothetical protein LOC100275184 | Negative Regulator | NO | FN | HA2 |
| GRMZM2G062244 | Zm00001d006687 | NA | Negative Regulator | NO | FN | HA2 |
| GRMZM2G137510 | Zm00001d023736 | MADS-box transcription factor 14 | Negative Regulator | NO | FN | HA2 |
| GRMZM2G181030 | Zm00001d026017 | hypothetical protein LOC100273483 | Negative Regulator | NO | FN | HD2 |
| GRMZM2G005155 | Zm00001d002332 | hypothetical protein LOC100272346 | Negative Regulator | NO | FN | HN |
| GRMZM2G077124 | Zm00001d022550 | hypothetical protein LOC100191211 | Negative Regulator | NO | FN | HN |
| GRMZM2G132367 | Zm00001d027991 | homeodomain-leucine zipper transcription factor TaHDZipI-1 | Negative Regulator | NO | FN | HN |
| GRMZM2G156006 | Zm00001d045120 | NA | Negative Regulator | NO | FN | HN |
| GRMZM2G171600 | Zm00001d042313 | hypothetical protein | Negative Regulator | NO | FN | HN |

| | | |
|---|---|---|
| | | LOC100384320 |

**Supplementary Table S2:**

**Genes selected from study 2 (Liu et al., 2022).** The module "M2" indicates genes with higher expression in pre-Bundle Sheath cells than in pre-mesophyll cells; M3 and M4 indicate genes expressed in pre-Bundle Sheath cells but decreased in later stages. Genes without complete orthogroups are highlighted in red.

| Gene ID | Gene ID (Gramene database) | TF Family | Module |
|---|---|---|---|
| AC202864.3_FG002 | Zm00001d038801 | GATA | M2 |
| AC203865.3_FG001 | Zm00001d041444 | CPP | M2 |
| AC234164.1_FG004 | Zm00001d045507 | GRAS | M2 |
| GRMZM2G000171 | Zm00001d014795 | bZIP | M2 |
| GRMZM2G001048 | Zm00001d049094 | B3 | M2 |
| GRMZM2G002978 | Zm00001d003836 | Trihelix | M2 |
| GRMZM2G003304 | Zm00001d032316 | HD-ZIP | M2 |
| GRMZM2G005353 | Zm00001d025944 | YABBY | M2 |
| GRMZM2G006493 | Zm00001d023636 | C3H | M2 |
| GRMZM2G008234 | Zm00001d018731 | AP2 | M2 |
| GRMZM2G008898 | Zm00001d007768 | bHLH | M2 |
| GRMZM2G009530 | Zm00001d031135 | GATA | M2 |
| GRMZM2G013391 | Zm00001d012746 | WRKY | M2 |
| GRMZM2G015080 | Zm00001d005029 | GRAS | M2 |
| GRMZM2G015666 | Zm00001d005939 | bHLH | M2 |
| GRMZM2G018414 | Zm00001d033396 | GRF | M2 |
| GRMZM2G018487 | Zm00001d033965 | WRKY | M2 |
| GRMZM2G021573 | Zm00001d048004 | AP2 | M2 |
| GRMZM2G024247 | Zm00001d043946 | HMG | M2 |
| GRMZM2G028046 | Zm00001d053006 | C2H2 | M2 |
| GRMZM2G028151 | Zm00001d034204 | AP2 | M2 |
| GRMZM2G028492 | Zm00001d026536 | C3H | M2 |
| GRMZM2G028980 | Zm00001d053819 | ARF | M2 |
| GRMZM2G030744 | Zm00001d018416 | bHLH | M2 |
| GRMZM2G034840 | Zm00001d001945 | ARF | M2 |
| GRMZM2G036837 | Zm00001d008322 | C3H | M2 |
| GRMZM2G037200 | Zm00001d017595 | C3H | M2 |
| GRMZM2G042895 | Zm00001d024522 | bHLH | M2 |
| GRMZM2G054795 | Zm00001d002829 | YABBY | M2 |

| | | | |
|---|---|---|---|
| GRMZM2G060114 | Zm00001d006629 | mTERF | M2 |
| GRMZM2G064197 | Zm00001d020019 | G2-like | M2 |
| GRMZM2G065538 | Zm00001d011639 | RAV | M2 |
| GRMZM2G069274 | Zm00001d043076 | WOX | M2 |
| GRMZM2G073750 | Zm00001d041497 | ARF | M2 |
| GRMZM2G073826 | Zm00001d016000 | MYB | M2 |
| GRMZM2G077752 | Zm00001d011843 | SRS | M2 |
| GRMZM2G081816 | Zm00001d044242 | bHLH | M2 |
| GRMZM2G081892 | Zm00001d008546 | ERF | M2 |
| GRMZM2G082586 | Zm00001d020790 | bHLH | M2 |
| GRMZM2G083504 | Zm00001d006065 | bHLH | M2 |
| GRMZM2G083886 | Zm00001d018208 | GeBP | M2 |
| GRMZM2G085427 | Zm00001d028380 | MYB_related | M2 |
| GRMZM2G086072 | Zm00001d027709 | E2F/DP | M2 |
| GRMZM2G086474 | Zm00001d026046 | bHLH | M2 |
| GRMZM2G087804 | Zm00001d039260 | G2-like | M2 |
| GRMZM2G088242 | Zm00001d005843 | HSF | M2 |
| GRMZM2G088524 | Zm00001d029963 | MYB_related | M2 |
| GRMZM2G091331 | Zm00001d026252 | WRKY | M2 |
| GRMZM2G092167 | Zm00001d052317 | LBD | M2 |
| GRMZM2G094935 | Zm00001d038812 | HB-other | M2 |
| GRMZM2G095727 | Zm00001d047761 | Pseudo ARR-B | M2 |
| GRMZM2G096709 | Zm00001d021362 | GRF | M2 |
| GRMZM2G099797 | Zm00001d028265 | ARR-B | M2 |
| GRMZM2G099862 | Zm00001d007465 | GRF | M2 |
| GRMZM2G100794 | Zm00001d004651 | C3H | M2 |
| GRMZM2G102845 | Zm00001d015243 | ARF | M2 |
| GRMZM2G111123 | Zm00001d044355 | B3 | M2 |
| GRMZM2G111204 | Zm00001d005584 | HB-other | M2 |
| GRMZM2G111906 | Zm00001d036356 | MYB_related | M2 |
| GRMZM2G113127 | Zm00001d031310 | NF-YC | M2 |
| GRMZM2G114613 | Zm00001d041474 | LSD | M2 |
| GRMZM2G116557 | Zm00001d011953 | ARF | M2 |
| GRMZM2G123909 | Zm00001d043969 | GATA | M2 |
| GRMZM2G131982 | Zm00001d002806 | DBB | M2 |
| GRMZM2G133398 | Zm00001d042396 | B3 | M2 |

| | | | |
|---|---|---|---|
| GRMZM2G146020 | Zm00001d041920 | bZIP | M2 |
| GRMZM2G146514 | Zm00001d049223 | C3H | M2 |
| GRMZM2G146688 | Zm00001d007840 | AP2 | M2 |
| GRMZM2G147712 | Zm00001d050242 | NF-YB | M2 |
| GRMZM2G147880 | Zm00001d013307 | WRKY | M2 |
| GRMZM2G148723 | Zm00001d039271 | bHLH | M2 |
| GRMZM2G150011 | Zm00001d020037 | C2H2 | M2 |
| GRMZM2G151689 | Zm00001d046740 | C3H | M2 |
| GRMZM2G155123 | Zm00001d010974 | MIZ | M2 |
| GRMZM2G155980 | Zm00001d034813 | FAR1 | M2 |
| GRMZM2G156785 | Zm00001d051427 | HMG | M2 |
| GRMZM2G158162 | Zm00001d022440 | B3 | M2 |
| GRMZM2G159456 | Zm00001d027859 | bHLH | M2 |
| GRMZM2G159475 | Zm00001d041536 | GRAS | M2 |
| GRMZM2G163813 | Zm00001d053775 | SBP | M2 |
| GRMZM2G167824 | Zm00001d032502 | YABBY | M2 |
| GRMZM2G171179 | Zm00001d045044 | ERF | M2 |
| GRMZM2G171370 | Zm00001d005244 | bZIP | M2 |
| GRMZM2G171650 | Zm00001d042315 | M-type_MADS | M2 |
| GRMZM2G178102 | Zm00001d041489 | HD-ZIP | M2 |
| GRMZM2G178261 | Zm00001d033876 | GRF | M2 |
| GRMZM2G179069 | Zm00001d035651 | Dof | M2 |
| GRMZM2G314546 | Zm00001d043452 | HB-other | M2 |
| GRMZM2G320827 | Zm00001d002782 | Trihelix | M2 |
| GRMZM2G336533 | Zm00001d013003 | NAC | M2 |
| GRMZM2G350165 | Zm00001d000291 | bHLH | M2 |
| GRMZM2G379179 | Zm00001d031266 | Trihelix | M2 |
| GRMZM2G385543 | Zm00001d013357 | bHLH | M2 |
| GRMZM2G398506 | Zm00001d021947 | WRKY | M2 |
| GRMZM2G399072 | Zm00001d027878 | AP2 | M2 |
| GRMZM2G403620 | Zm00001d030737 | MYB | M2 |
| GRMZM2G412430 | Zm00001d029402 | bHLH | M2 |
| GRMZM2G427087 | Zm00001d025982 | Trihelix | M2 |
| GRMZM2G448104 | Zm00001d026362 | MYB_related | M2 |
| GRMZM2G481163 | Zm00001d029974 | Trihelix | M2 |
| GRMZM2G488465 | Zm00001d052781 | Pseudo ARR-B | M2 |

| | | | |
|---|---|---|---|
| GRMZM2G529859 | Zm00001d017391 | YABBY | M2 |
| GRMZM2G531738 | Zm00001d037836 | MYB | M2 |
| GRMZM2G589696 | Zm00001d051577 | Dof | M2 |
| GRMZM5G803308 | Zm00001d046632 | MYB | M2 |
| GRMZM5G813892 | Zm00001d029875 | MYB_related | M2 |
| GRMZM5G821024 | Zm00001d004857 | bZIP | M2 |
| GRMZM5G834758 | Zm00001d014858 | HMG | M2 |
| GRMZM5G880069 | Zm00001d013709 | WRKY | M2 |
| GRMZM5G887286 | Zm00001d007339 | C2H2 | M2 |
| GRMZM5G887975 | Zm00001d009604 | GATA | M2 |
| GRMZM2G014653 | Zm00001d042609 | NAC | M3 |
| GRMZM2G141031 | Zm00001d020683 | C2H2 | M3 |
| GRMZM2G166946 | Zm00001d043368 | TCP | M3 |
| GRMZM2G344521 | Zm00001d033324 | B3 | M3 |
| GRMZM2G361659 | Zm00001d052288 | E2F/DP | M3 |
| GRMZM2G001777 | Zm00001d007675 | HB-other | M4 |
| GRMZM2G005126 | Zm00001d024703 | C3H | M4 |
| GRMZM2G012654 | Zm00001d019865 | NF-YB | M4 |
| GRMZM2G035405 | Zm00001d014377 | ARF | M4 |
| GRMZM2G037630 | Zm00001d029489 | NF-YA | M4 |
| GRMZM2G039586 | Zm00001d046354 | GATA | M4 |
| GRMZM2G058451 | Zm00001d021701 | bHLH | M4 |
| GRMZM2G086949 | Zm00001d026540 | ARF | M4 |
| GRMZM2G092609 | Zm00001d016732 | bZIP | M4 |
| GRMZM2G104078 | Zm00001d050960 | NAC | M4 |
| GRMZM2G112483 | Zm00001d003180 | bZIP | M4 |
| GRMZM2G168269 | Zm00001d020975 | MBD | M4 |
| GRMZM2G173321 | Zm00001d024321 | B3 | M4 |
| GRMZM2G470307 | Zm00001d039306 | MYB | M4 |
| GRMZM5G865212 | Zm00001d041549 | Dof | M4 |

**Supplementary Table S3:**

**Genes selected for the control set (source: Wang et al., 2013).** Under the columns labeled "Foliar profile" and "Husk profile", the prefixes "F" and "H" denote foliar and husk tissues, respectively, across developmental progression from primordium (P1) to P3/4 to P5. In both tissues, A-type profiles (FA, HA) show increasing trends—either continuous (A1), early increase followed by stabilization (A2), or late-stage increase (A3)—while D-type profiles (FD, HD) show corresponding decreasing patterns across these stages (D1–D3). FN and HN represent genes or TFs with no clear developmental trend (neutral) in foliar and husk tissues, respectively. Genes without complete orthogroups are highlighted in red.

| Genes IDs | Annotation | Expression profile in Study 1 | |
|---|---|---|---|
| | | Foliar profile | Husk profile |
| GRMZM2G014136 | acyl-coenzyme A oxidase 2 | FN | HN |
| GRMZM2G033846 | caltractin | FN | HN |
| GRMZM2G038950 | mitochondrial import inner membrane translocase subunit TIM14 | FN | HN |
| GRMZM2G046615 | terpene synthase 2 | NA | HN |
| GRMZM2G055970 | nucleic acid binding protein | FN | HN |
| GRMZM2G057823 | aldolase1 | FN | HN |
| GRMZM2G059580 | elongation factor 1-gamma 3 | FN | HN |
| GRMZM2G061900 | ras-related protein ARA-3 | FN | HN |
| GRMZM2G062481 | acyl carrier protein | FN | HN |
| GRMZM2G066142 | agmatine coumaroyltransferase | FN | HN |
| GRMZM2G072448 | cysteine protease | NA | NA |
| GRMZM2G073912 | 3,4-dihydroxy-2-butanone kinase | FN | HN |
| GRMZM2G073969 | | NA | NA |
| GRMZM2G097059 | MADS-domain transcription factor 4 | NA | NA |
| GRMZM2G097316 | root cap protein 1 | NA | NA |
| GRMZM2G104613 | 3-isopropylmalate dehydrogenase 2 | FN | HN |
| GRMZM2G111309 | amidase | FN | HN |
| GRMZM2G117146 | sugar transport protein 8 | NA | NA |
| GRMZM2G119265 | beta-1,3-galactosyltransferase sqv-2 | NA | NA |
| GRMZM2G125072 | 3Fe-4S ferredoxin | FN | HN |
| GRMZM2G137751 | calcineurin B-like protein 4 | NA | NA |
| GRMZM2G145027 | microtubule motor | FN | HN |
| GRMZM2G146206 | triosephosphate isomerase, cytosolic | FN | HN |

| | | | |
|---|---|---|---|
| GRMZM2G146502 | root cap periphery gene2 | NA | NA |
| GRMZM2G149182 | glutathione S-transferase GST 34 | NA | HN |
| GRMZM2G151564 | calcium ion binding protein | FN | HN |
| GRMZM2G157018 | ATP synthase D chain, mitochondrial | FN | HN |
| GRMZM2G157629 | seed maturation protein | NA | NA |
| GRMZM2G158526 | centromeric histone H3 | FN | HN |
| GRMZM2G160887 | GTP binding protein | FN | HN |
| GRMZM2G162505 | chitinase 2 | NA | NA |
| GRMZM2G167149 | pectinesterase inhibitor domain containing protein | NA | NA |
| GRMZM2G168143 | ferredoxin | FN | HN |
| GRMZM2G173090 | heat shock factor protein HSF30 | NA | NA |
| GRMZM2G173341 | alpha-1,4-glucan-protein synthase | FN | HN |
| GRMZM2G175343 | flavonol synthase/flavanone 3-hydroxylase | NA | HN |
| GRMZM2G307152 | transcriptional factor TINY | NA | HN |
| GRMZM2G328780 | disulfide oxidoreductase/ monooxygenase/ oxidoreductase | NA | NA |
| GRMZM2G347047 | CDPK protein | NA | NA |
| GRMZM2G357124 | LOC100284520, RALF | NA | NA |
| GRMZM2G366434 | protein BABY BOOM 1 | NA | NA |
| GRMZM2G375517 | 22.0 kDa class IV heat shock protein | NA | NA |
| GRMZM2G403620 | rough sheath2 | FN | HN |
| GRMZM2G415390 | | FN | NA |
| GRMZM2G417945 | limonoid UDP-glucosyltransferase | FN | HN |
| GRMZM2G442008 | peroxidase 65 | NA | NA |
| GRMZM2G451882 | calmodulin binding protein | FN | HN |
| GRMZM2G473711 | beta-glucanase | FN | NA |
| GRMZM2G479596 | calmodulin binding protein | NA | NA |
| GRMZM2G538064 | endoglucanase | NA | HN |

Supplementary Table S4:
Maize gene ID of the orthogroups that formed unresolvable complex gene-trees.

| Genes | Dataset |
|---|---|
| GRMZM2G060544 | test |
| GRMZM2G095899 | test |
| GRMZM2G098988 | test |
| GRMZM2G131577 | test |
| GRMZM2G132794 | test |
| GRMZM2G136494 | test |
| GRMZM2G146688 | test |
| GRMZM2G156006 | test |
| GRMZM2G172657 | test |
| GRMZM2G462623 | test |
| GRMZM5G887276 | test |
| GRMZM2G008234 | test |
| GRMZM2G008898 | test |
| GRMZM2G060114 | test |
| GRMZM2G086072 | test |
| GRMZM2G091331 | test |
| GRMZM2G111204 | test |
| GRMZM2G113127 | test |
| GRMZM2G123909 | test |
| GRMZM2G151689 | test |
| GRMZM2G156785 | test |
| GRMZM2G314546 | test |
| GRMZM2G336533 | test |
| GRMZM2G344521 | test |
| GRMZM5G834758 | test |
| GRMZM5G887975 | test |
| GRMZM2G059580 | control |
| GRMZM2G061900 | control |
| GRMZM2G066142 | control |
| GRMZM2G137751 | control |
| GRMZM2G145027 | control |

| | |
|---|---|
| GRMZM2G347047 | control |
| GRMZM2G357124 | control |
| GRMZM2G366434 | control |
| GRMZM2G167149 | control |

Supplementary Table S5:

Likelihood Ratio Test (LRT) results of the orthogroups of the test-set, where the $C_4$ branches showed relatively higher dN/dS (>0.3), and the entire control-set. The test statistic (2ΔlnL) denotes twice the difference in log-likelihoods between alternative and null codon models and is used to evaluate model fit in likelihood ratio tests.

| Set | Gene | 2ΔlnL | P value | Passed/Failed |
|---|---|---|---|---|
| Test | GRMZM2G171179 | 30.73 | 2.92E-08 | Passed |
| | GRMZM2G123900 | 23.9 | 1.01E-06 | Passed |
| | GRMZM2G042895 | 20.4 | 6.29E-06 | Passed |
| | GRMZM2G005155 | 12.62 | 3.82E-04 | Passed |
| | GRMZM2G015666 | 7.59 | 5.85E-03 | Passed |
| | GRMZM2G082586 | 7.59 | 5.85E-03 | Passed |
| | GRMZM2G131516 | 7.05 | 7.90E-03 | Passed |
| | GRMZM2G015080 | 7.05 | 7.90E-03 | Passed |
| | GRMZM2G147712 | 6.12 | 1.33E-02 | Passed |
| | GRMZM2G379179 | 5.6 | 1.79E-02 | Passed |
| | GRMZM2G036837 | 4.39 | 3.62E-02 | Passed |
| | GRMZM2G064638 | 4.26 | 3.90E-02 | Passed |
| | GRMZM2G085751 | 4.26 | 3.90E-02 | Passed |
| | GRMZM2G140694 | 3.582496 | 5.84E-02 | Failed |
| | GRMZM2G021573 | 2.196138 | 1.38E-01 | Failed |
| | GRMZM2G045883 | 2.140888 | 1.43E-01 | Failed |
| | GRMZM2G179069 | 1.927878 | 1.65E-01 | Failed |
| | GRMZM2G087804 | 1.777066 | 1.83E-01 | Failed |
| | GRMZM2G040924 | 1.415976 | 2.34E-01 | Failed |
| | GRMZM2G081816 | 1.37165 | 2.42E-01 | Failed |
| | GRMZM2G062244 | 1.267228 | 2.60E-01 | Failed |
| | GRMZM2G150011 | 1.238698 | 2.66E-01 | Failed |
| | GRMZM2G111123 | 1.138744 | 2.86E-01 | Failed |
| | GRMZM2G173321 | 1.138744 | 2.86E-01 | Failed |
| | GRMZM2G481163 | 0.72456 | 3.95E-01 | Failed |
| | GRMZM2G018414 | 0.709836 | 3.99E-01 | Failed |
| | GRMZM2G147880 | 0.5614 | 4.54E-01 | Failed |

| | | | | |
|---|---|---|---|---|
| | GRMZM2G018487 | 0.5522 | 4.57E-01 | Failed |
| | GRMZM2G104078 | 0.498952 | 4.80E-01 | Failed |
| | GRMZM2G002280 | 0.387776 | 5.33E-01 | Failed |
| | GRMZM2G069365 | 0.373408 | 5.41E-01 | Failed |
| | GRMZM2G417229 | 0.373408 | 5.41E-01 | Failed |
| | GRMZM2G114613 | 0.236706 | 6.27E-01 | Failed |
| | GRMZM2G111906 | 0.083968 | 7.72E-01 | Failed |
| | GRMZM2G168269 | 0.078702 | 7.79E-01 | Failed |
| | GRMZM2G361659 | 0.023886 | 8.77E-01 | Failed |
| | GRMZM2G132367 | 0.001108 | 9.73E-01 | Failed |
| Control | GRMZM2G102760 | 3.58932 | 5.82E-02 | Failed |
| | GRMZM2G018375 | 2.785034 | 9.51E-02 | Failed |
| | GRMZM2G157018 | 2.779618 | 9.55E-02 | Failed |
| | GRMZM2G117146 | 2.558998 | 1.10E-01 | Failed |
| | GRMZM2G119265 | 2.452146 | 1.17E-01 | Failed |
| | GRMZM2G149182 | 2.163934 | 1.41E-01 | Failed |
| | GRMZM2G403620 | 1.933588 | 1.64E-01 | Failed |
| | GRMZM2G088396 | 1.846292 | 1.74E-01 | Failed |
| | GRMZM2G062481 | 1.622384 | 2.03E-01 | Failed |
| | GRMZM2G146502 | 1.614864 | 2.04E-01 | Failed |
| | GRMZM2G160887 | 1.560196 | 2.12E-01 | Failed |
| | GRMZM2G038950 | 1.460012 | 2.27E-01 | Failed |
| | GRMZM2G033846 | 1.42468 | 2.33E-01 | Failed |
| | GRMZM2G097059 | 1.263184 | 2.61E-01 | Failed |
| | GRMZM2G074543 | 1.16612 | 2.80E-01 | Failed |
| | GRMZM2G097316 | 1.126064 | 2.89E-01 | Failed |
| | GRMZM2G072448 | 0.928984 | 3.35E-01 | Failed |
| | GRMZM2G055970 | 0.916014 | 3.39E-01 | Failed |
| | GRMZM2G175343 | 0.54022 | 4.62E-01 | Failed |
| | GRMZM2G125072 | 0.500814 | 4.79E-01 | Failed |

| | | | | |
|---|---|---|---|---|
| | GRMZM2G006721 | 0.405914 | 5.24E-01 | Failed |
| | GRMZM2G173090 | 0.339764 | 5.60E-01 | Failed |
| | GRMZM2G375517 | 0.330576 | 5.65E-01 | Failed |
| | GRMZM2G112530 | 0.318762 | 5.72E-01 | Failed |
| | GRMZM2G479596 | 0.2921 | 5.89E-01 | Failed |
| | GRMZM2G180319 | 0.215272 | 6.43E-01 | Failed |
| | GRMZM2G088309 | 0.195584 | 6.58E-01 | Failed |
| | GRMZM2G451882 | 0.173708 | 6.77E-01 | Failed |
| | GRMZM2G417945 | 0.152754 | 6.96E-01 | Failed |
| | GRMZM2G073969 | 0.123694 | 7.25E-01 | Failed |
| | GRMZM2G158526 | 0.120616 | 7.28E-01 | Failed |
| | GRMZM2G441325 | 0.104932 | 7.46E-01 | Failed |
| | GRMZM2G073912 | 0.081252 | 7.76E-01 | Failed |
| | GRMZM2G307152 | 0.05827 | 8.09E-01 | Failed |
| | GRMZM2G046615 | 0.003814 | 9.51E-01 | Failed |
| | GRMZM2G442008 | 0.000232 | 9.88E-01 | Failed |

Supplementary Table S6:
Functional annotation of genes that passed the Likelihood Ratio Test (LRT) test.

| Gene names | Maize representative from ortholog cluster | Description | Gene Annotation of the maize representative | | | | | Mutation study |
|---|---|---|---|---|---|---|---|---|
| | | | Gene Ontology | | | Interpro terms | Mapman | |
| | | | Biological Process | Molecular Function | Cellular Component | | | |
| bhlh33 | GRMZM2G015666 / Zm00001d005939 | bHLH-transcription factor 33 | - GO:0006357 regulation of transcription by RNA polymerase II<br><br>- GO:0006355 regulation of transcription, DNA-templated<br><br>- GO:0009631 cold acclimation | - GO:0046983 protein dimerization activity<br><br>- GO:0000981 DNA-binding transcription factor activity, RNA polymerase II-specific<br><br>- GO:0000978 RNA polymerase II cis-regulatory region sequence-specific DNA binding | GO:0005634 nucleus | IPR011598 Myc-type, basic helix-loop-helix (bHLH) domain IPR036638 Helix-loop-helix DNA-binding domain superfamily | 15.5.30 RNA biosynthesis.transcriptional regulation.transcription factor bHLH) | Normal |
| bhlh105 | GRMZM2G082586 / Zm00001d020790 | bhlh105 - bHLH-transcription factor 105 | - GO:0006357 regulation of transcription by RNA polymerase II<br><br>- GO:0006355 regulation of transcription, DNA-templated | - GO:0046983 protein dimerization activity<br><br>- GO:0000981 DNA-binding transcription factor activity, RNA polymerase II-specific<br><br>- GO:0000978 RNA polymerase II cis-regulatory region sequence-specific DNA binding | GO:0005634 nucleus Os09g0468700 | IPR036638 Helix-loop-helix DNA-binding domain superfamily IPR011598 Myc-type, basic helix-loop-helix (bHLH) domain | 15.5.30 RNA biosynthesis.transcriptional regulation.transcription factor (bHLH) | Normal |

| | | | | | | | | |
|---|---|---|---|---|---|---|---|---|
| ereb160 | GRMZM2G171179/ Zm00001d045044 | ereb160 - AP2-EREB P-transcrip tion factor 160 | - GO:0071456 cellular response to hypoxia<br><br>- GO:0010104 regulation of ethylene-activated signaling pathway<br><br>- GO:0071369 cellular response to ethylene stimulus<br><br>- GO:0070483 detection of hypoxia<br><br>- GO:0045893 positive regulation of transcription, DNA-templated<br><br>- GO:0071454 cellular response to anoxia<br><br>- GO:0009733 response to auxin<br><br>- GO:0009723 response to ethylene<br><br>- GO:2000280 regulation of root development | - GO:0005515 protein binding<br><br>- GO:0003677 DNA binding<br><br>- GO:0000976 transcription cis-regulatory region binding<br><br>- GO:0003700 DNA-binding transcription factor activity | GO:0005622<br><br>intracellular anatomical structure GO:0005886<br><br>plasma membrane GO:0005634 nucleus | IPR016177 DNA-binding domain superfamily IPR001471 AP2/ERF domain IPR036955 AP2/ERF domain superfamily IPR044808 Ethylene-responsiv e transcription factor | 15.5.7.1 RNA biosynthesis.trans criptional regulation.AP2/E RF transcription factor superfamily.transc ription factor (ERF) | - |

| | | | | | | | | |
|---|---|---|---|---|---|---|---|---|
| cadtfr3 | GRMZM2G147712 / Zm00001d050242 | cadtfr3 - CCAAT-DR 1 transcription factor | GO:0006355 regulation of transcription, DNA-templated | - GO:0046982 protein heterodimerization activity<br><br>- GO:0001228 DNA-binding transcription activator activity, RNA polymerase II-specific<br><br>- GO:0005515 protein binding<br><br>- GO:0000976 transcription cis-regulatory region binding | GO:0016602 CCAAT-binding factor complex GO:0005634 nucleus | IPR009072 Histone-fold IPR003958 Transcription factor CBF/NF-Y/archaeal histone domain IPR044255 Protein Dr1 homolog | 15.3.3.6.1.2 RNA biosynthesis.RNA polymerase II-dependent transcription.transcription initiation.TATA box-binding protein (TBP) regulation.NC2 regulator heterodimer.component beta | - |
| thx8 | GRMZM2G379179 / Zm00001d031266 | thx8 - Trihelix-transcription factor 8 | - GO:0019760 glucosinolate metabolic process<br><br>- GO:0006355 regulation of transcription, DNA-templated | - GO:0003677 DNA binding<br><br>- GO:0000976 transcription cis-regulatory region binding | GO:0005634 nucleus | IPR044822 Myb/SANT-like DNA-binding domain 4 IPR006578 MADF domain IPR044823 Trihelix transcription factor ASIL1/2-like | 15.5.20 RNA biosynthesis.transcriptional regulation.transcription factor (Trihelix) | - |
| mads9 | GRMZM2G005155/ Zm00001d002332 | mads9 - MADS-transcription factor 9 | - GO:0006357 regulation of transcription by RNA polymerase II<br><br>- GO:0006355 regulation of transcription, DNA-templated | - GO:0003677 DNA binding<br><br>- GO:0000978 RNA polymerase II cis-regulatory region sequence-specific DNA binding<br><br>- GO:0046983 protein | GO:0005634 nucleus | IPR002100 Transcription factor, MADS-box IPR036879 Transcription factor, MADS-box superfamily IPR002487 Transcription factor, K-box | 15.5.14 RNA biosynthesis.transcriptional regulation.transcription factor (MADS/AGL) | |

| | | | | | | | | |
|---|---|---|---|---|---|---|---|---|
| | | | | dimerization activity | | IPR033896 MADS MEF2-like | | |
| C3H28 | GRMZM2G036837/ Zm00001d008322 | c3h28 - C3H-transcription factor 328 | - GO:0046872 metal ion binding<br><br>- GO:0003677 DNA binding<br><br>- GO:0003729 mRNA binding | | | IPR036855 Zinc finger, CCCH-type superfamily IPR000571 Zinc finger, CCCH-type | 15.5.16 RNA biosynthesis.transcriptional regulation.transcription factor (C3H-ZF) | |
| BHLH116 | GRMZM2G042895/ Zm00001d024522 | bhlh116 - bHLH-transcription factor 116 | - GO:0006355 regulation of transcription, DNA-templated<br><br>- GO:0009867 jasmonic acid mediated signaling pathway<br><br>- GO:0048831 regulation of shoot system development<br><br>- GO:0055062 phosphate ion homeostasis<br><br>- GO:0080040 positive regulation of cellular response to phosphate starvation | - GO:0005515 protein binding<br><br>- GO:0042803 protein homodimerization activity<br><br>- GO:0003700 DNA-binding transcription factor activity<br><br>- GO:0043565 sequence-specific DNA binding | GO:0005737<br><br>cytoplasm GO:0005634<br><br>nucleus | IPR045084 Transcription factor AIB/MYC-like IPR036638 Helix-loop-helix DNA-binding domain superfamily IPR011598 Myc-type, basic helix-loop-helix (bHLH) domain | 15.5.30 RNA biosynthesis.transcriptional regulation.transcription factor (bHLH) | Putative regulator Wang et al. 2014. Comparative analyses of C4 and C3 photosynthesis in developing leaves of maize and rice |

Supplementary Table S7:

Enriched motifs matching the previously identified motifs driving $C_4$-specific expression.

| Gene name | Gene id | Known proximal motifs (ref. ) | Matching motifs to known motifs |
|---|---|---|---|
| PPC1 | GRMZM2G083841 | 5 conserved nucleotide sequences (CNSs), putative TATA box (Gupta et al., 2020) | CNS 3a,3b conserved |
| RBC-SSU1 | GRMZM2G098520 | Ibox (Giuliano et al., 1988),<br>Homo motif, (Xu et al., 2001) | Ibox was observed in one species. In others partially conserved. |
| NADP-ME | GRMZM2G085019 | bHLH129 motif, AP2-EREBP, MYB motifs (Burgess et al.,2019) | none |
| Transketolase | GRMZM2G033208 | Digital genomic footprint co-opted from Ancestral $C_3$ genomes (Burgess et al.,2019) | none |

Supplementary Table S8:

Transcription factor families binding to motifs enriched in control and test sets

| Family | Test | Control | p_value | odds_ratio | Enriched | sig |
|---|---|---|---|---|---|---|
| AP2 | 2 | 0 | 0.1984732824 | 0 | Control | |
| ARF | 2 | 0 | 0.358778626 | 0.24 | Control | |
| C2H2 | 11 | 1 | 0.001189172765 | 0.22 | Control | Significant |
| DOF | 9 | 2 | 0.4880170424 | 0.49 | Control | |
| E2F | 5 | 0 | 1 | 0.99 | Control | |
| ERF/DREB | 31 | 17 | 1 | 1.12 | Test | |
| GARP_G2-like | 1 | 0 | 1 | 1.40 | Test | |
| Group A | 1 | 0 | 0.4582080817 | 2.91 | Test | |
| Group G | 0 | 1 | 1 | Inf | Test | |
| Group S | 1 | 1 | 1 | Inf | Test | |
| HD-ZIP | 1 | 0 | 1 | Inf | Test | |
| IDD | 2 | 0 | 1 | Inf | Test | |
| LAV | 3 | 0 | 1 | Inf | Test | |
| LBD | 2 | 0 | 1 | Inf | Test | |
| MYB | 4 | 0 | 1 | Inf | Test | |
| MYB-related | 3 | 0 | 1 | Inf | Test | |
| NAC | 8 | 0 | 1 | Inf | Test | |
| SBP | 2 | 1 | 1 | Inf | Test | |
| Type II | 2 | 0 | 0.584013847 | Inf | Test | |
| WRKY | 4 | 1 | 0.5823761062 | Inf | Test | |
| bHLH | 11 | 2 | 0.3555291367 | Inf | Test | |

Supplementary Table S9:

Motifs that are Non-syntenic or $C_4$-shifted in test and control

| Dataset | Presence Class | Position_Class | Consensus Sequences | Orthogroup | Gene Name | TF binding to the motif |
|---|---|---|---|---|---|---|
| Test | PACMAD | c4_shifted | GCCYCKMTCGCTTCC | GRMZM2G003304 | ZmHB112 | NA |
| Test | PACMAD | c4_shifted | GCCAGCACGCAMMAG | GRMZM2G003304 | ZmHB112 | NA |
| Test | PACMAD | c4_shifted | CCTAGATCCCAAAAT | GRMZM2G003304 | ZmHB112 | NA |
| Test | PACMAD | c4_shifted | KCAAGGTTSTTTGTG | GRMZM2G005155 | ZmMADS9 | NA |
| Test | PACMAD | c4_shifted | AAATKAACAGACAGT | GRMZM2G005155 | ZmMADS9 | NA |
| Test | PACMAD | c4_shifted | CAKTGCCATTTTCAG | GRMZM2G005155 | ZmMADS9 | NA |
| Test | PACMAD | c4_shifted | ACCGAACGTGTCRAA | GRMZM2G005353 | ZmYAB14 | NA |
| Test | PACMAD | c4_shifted | ACGCCCCTATATAAA | GRMZM2G005353 | ZmYAB14 | NA |
| Test | PACMAD | c4_shifted | GATTGGAACTAACAT | GRMZM2G005353 | ZmYAB14 | NA |
| Test | PACMAD | c4_shifted | CTGCCTCTCTCGGCT | GRMZM2G005353 | ZmYAB14 | NA |
| Test | PACMAD | c4_shifted | CRTCAAATCCAACGC | GRMZM2G005353 | ZmYAB14 | NA |
| Test | PACMAD | c4_shifted | YTCTGATAGGACCGC | GRMZM2G005353 | ZmYAB14 | NA |
| Test | PACMAD | c4_shifted | GCTGTRATCGCCTGC | GRMZM2G005353 | ZmYAB14 | NA |
| Test | PACMAD | c4_shifted | TCAATCAAATCCGYC | GRMZM2G005353 | ZmYAB14 | WOX13 |
| Test | PACMAD | c4_shifted | GCTGCYGCCACTGTT | GRMZM2G005353 | ZmYAB14 | NA |
| Test | PACMAD | c4_shifted | GGACAGGACCAGACC | GRMZM2G005353 | ZmYAB14 | NA |
| Test | PACMAD | c4_shifted | GCGGTCGCACGCTAG | GRMZM2G006493 | ZmC3H54 | NA |
| Test | PACMAD | c4_shifted | GAAGCATCCAAGRAA | GRMZM2G006493 | ZmC3H54 | NA |
| Test | PACMAD | c4_shifted | TYCACAGCYGTWGCC | GRMZM2G009530 | ZmGATA9 | NA |
| Test | PACMAD | c4_shifted | AGCYSTGGCACRTCC | GRMZM2G013391 | ZmWRKY111 | NA |
| Test | PACMAD | c4_shifted | GTACTTTCBCGGCCG | GRMZM2G014653 | ZmNAC109 | NA |
| Test | PACMAD | c4_shifted | ACGCVGGCGCGCGTC | GRMZM2G014653 | ZmNAC109 | NA |
| Test | PACMAD | c4_shifted | TAGRGAGCGYGYCYT | GRMZM2G015666 | ZmbHLH33 | NA |
| Test | PACMAD | c4_shifted | GCCCGAGGAAGGAAT | GRMZM2G018487 | ZmWRKY107 | NA |
| Test | PACMAD | c4_shifted | GAAAAAGTCCAAGCG | GRMZM2G018487 | ZmWRKY107 | GLYMA-14G028900 |
| Test | PACMAD | c4_shifted | TCRCDCACAGCGCAC | GRMZM2G018487 | ZmWRKY107 | NA |
| Test | PACMAD | c4_shifted | CKTCSWCAKGAACAG | GRMZM2G028980 | ZmARF16 | NA |
| Test | PACMAD | c4_shifted | CKTCSWCAKGAACAG | GRMZM2G035405 | ZmARF18 | NA |
| Test | PACMAD | c4_shifted | GCGTGGTCCWGCAGC | GRMZM2G036837 | ZmC3H28 | NA |
| Test | PACMAD | c4_shifted | ACCCTCAAGCGCCAT | GRMZM2G036837 | ZmC3H28 | NA |
| Test | PACMAD | c4_shifted | GGCAGGTGCGYGTRC | GRMZM2G045883 | ZmbHLH161 | NA |
| Test | PACMAD | c4_shifted | TCAGTTYCTTGCGAG | GRMZM2G045883 | ZmbHLH161 | NA |

| | | | | | | |
|---|---|---|---|---|---|---|
| Test | PACMAD | c4_shifted | ACRCCACCAACCATA | GRMZM2G045883 | ZmbHLH161 | MYB13 |
| Test | PACMAD | c4_shifted | AASAATCAAGCTAGC | GRMZM2G045883 | ZmbHLH161 | NA |
| Test | PACMAD | c4_shifted | YTGCACGCCTAAYTG | GRMZM2G045883 | ZmbHLH161 | NA |
| Test | PACMAD | c4_shifted | ACCGAACGTGTCRAA | GRMZM2G054795 | ZmYAB1 | NA |
| Test | PACMAD | c4_shifted | ACGCCCCTATATAAA | GRMZM2G054795 | ZmYAB1 | NA |
| Test | PACMAD | c4_shifted | GATTGGAACTAACAT | GRMZM2G054795 | ZmYAB1 | NA |
| Test | PACMAD | c4_shifted | CTGCCTCTCTCGGCT | GRMZM2G054795 | ZmYAB1 | NA |
| Test | PACMAD | c4_shifted | CRTCAAATCCAACGC | GRMZM2G054795 | ZmYAB1 | NA |
| Test | PACMAD | c4_shifted | YTCTGATAGGACCGC | GRMZM2G054795 | ZmYAB1 | NA |
| Test | PACMAD | c4_shifted | GCTGTRATCGCCTGC | GRMZM2G054795 | ZmYAB1 | NA |
| Test | PACMAD | c4_shifted | TCAATCAAATCCGYC | GRMZM2G054795 | ZmYAB1 | WOX13 |
| Test | PACMAD | c4_shifted | GCTGCYGCCACTGTT | GRMZM2G054795 | ZmYAB1 | NA |
| Test | PACMAD | c4_shifted | GGACAGGACCAGACC | GRMZM2G054795 | ZmYAB1 | NA |
| Test | PACMAD | c4_shifted | GCTCAACGGCTGGAA | GRMZM2G081892 | ZmEREB118 | NA |
| Test | PACMAD | c4_shifted | CCTCGTCGATGCTCA | GRMZM2G081892 | ZmEREB118 | NA |
| Test | PACMAD | c4_shifted | CCGATCTVGGAGGTT | GRMZM2G081892 | ZmEREB118 | NA |
| Test | PACMAD | c4_shifted | CATAAGGCCGTGTGG | GRMZM2G081892 | ZmEREB118 | NA |
| Test | PACMAD | c4_shifted | ACTGAYGGATTCGGT | GRMZM2G081892 | ZmEREB118 | NA |
| Test | PACMAD | c4_shifted | TCSTCCAGATCTGG | GRMZM2G081892 | ZmEREB118 | NA |
| Test | PACMAD | c4_shifted | GTCCAATTTTTGCAC | GRMZM2G081892 | ZmEREB118 | NA |
| Test | PACMAD | c4_shifted | AAADTTTCWTCTTTT | GRMZM2G081892 | ZmEREB118 | NA |
| Test | PACMAD | c4_shifted | GGAGGCGACGACGAC | GRMZM2G081892 | ZmEREB118 | NA |
| Test | PACMAD | c4_shifted | TAGRGAGCGYGYCYT | GRMZM2G082586 | ZmbHLH105 | NA |
| Test | PACMAD | c4_shifted | CAAAAGATCCTCGAG | GRMZM2G088242 | ZmHSF2 | NA |
| Test | PACMAD | c4_shifted | AAATGGCGGAYGGGT | GRMZM2G088242 | ZmHSF2 | NA |
| Test | PACMAD | c4_shifted | GGAAACCTGCACGTA | GRMZM2G088524 | ZmGLK62 | NA |
| Test | PACMAD | c4_shifted | CCCCACCVKCCAG | GRMZM2G095727 | ZmOrphan208 | NA |
| Test | PACMAD | c4_shifted | CCACCGACGTGCGGG | GRMZM2G095727 | ZmOrphan208 | NA |
| Test | PACMAD | c4_shifted | GATCTGCGGATCCGG | GRMZM2G095727 | ZmOrphan208 | NA |
| Test | PACMAD | c4_shifted | CGAGCYTGTTCWTGC | GRMZM2G095899 | ZmbHLH114 | NA |
| Test | PACMAD | c4_shifted | GCCCAAAATCCAGCG | GRMZM2G096709 | ZmGRF3 | NA |
| Test | PACMAD | c4_shifted | GGCTCGCCGTKTCTT | GRMZM2G098813 | ZmLFY2 | NA |
| Test | PACMAD | c4_shifted | GGGGGCGTAGAATCT | GRMZM2G098813 | ZmLFY2 | HHO6 |
| Test | PACMAD | c4_shifted | TAGGATWAGYTAWTT | GRMZM2G098988 | NA | NA |
| Test | PACMAD | c4_shifted | CGGCGCCDCSGSCS | GRMZM2G104078 | ZmNAC101 | NA |

| | | | | | | |
|---|---|---|---|---|---|---|
| Test | PACMAD | c4_shifted | CSSGCGKCSGCGGCG | GRMZM2G114613 | ZmLSD4 | NA |
| Test | PACMAD | c4_shifted | CTAGGGTTAGGKYTT | GRMZM2G114613 | ZmLSD4 | NA |
| Test | PACMAD | c4_shifted | TCGTCCTSCTCCTCC | GRMZM2G114613 | ZmLSD4 | NA |
| Test | PACMAD | c4_shifted | CCGTCCCYGCKKCGC | GRMZM2G114613 | ZmLSD4 | NA |
| Test | PACMAD | c4_shifted | GGBCCCCACCRGCT | GRMZM2G116557 | ZmARF25 | GLYMA-20G154400 |
| Test | PACMAD | c4_shifted | GYTGYGGCGTAGGWG | GRMZM2G116557 | ZmARF25 | NA |
| Test | PACMAD | c4_shifted | YGTCGSCRCTWTAAK | GRMZM2G123909 | ZmGATA35 | NA |
| Test | PACMAD | c4_shifted | CCACACCKCTCTTTK | GRMZM2G123909 | ZmGATA35 | NA |
| Test | PACMAD | c4_shifted | GTCTTACAMTAAGAA | GRMZM2G126018 | ZmSBP23 | NA |
| Test | PACMAD | c4_shifted | CTGCTGGCGCCGGCC | GRMZM2G126018 | ZmSBP23 | ereb18 |
| Test | PACMAD | c4_shifted | CTGTTCRTGYCACAG | GRMZM2G126018 | ZmSBP23 | NA |
| Test | PACMAD | c4_shifted | GTKTCYCTSTGCATG | GRMZM2G133398 | ZmABI1 | FUS3 |
| Test | PACMAD | c4_shifted | GAAGTARGGTGGCCG | GRMZM2G133398 | ZmABI1 | NA |
| Test | PACMAD | c4_shifted | CTACGGCGMCGAAAC | GRMZM2G133398 | ZmABI1 | Os05g0497200 |
| Test | PACMAD | c4_shifted | CKYCCTCKYTAMGCT | GRMZM2G137541 | ZmbHLH151 | NA |
| Test | PACMAD | c4_shifted | GGVCGGCCCWCCACA | GRMZM2G140669 | ZmGATA3 | NA |
| Test | PACMAD | c4_shifted | TTCCACCGCCATGGC | GRMZM2G146020 | ZmbZIP8 | NA |
| Test | PACMAD | c4_shifted | CCGGCAAACAGYCGC | GRMZM2G146020 | ZmbZIP8 | NA |
| Test | PACMAD | c4_shifted | CCCGCBTCCDCCGTM | GRMZM2G146020 | ZmbZIP8 | NA |
| Test | PACMAD | c4_shifted | AGTMCAHCAAAAGTG | GRMZM2G146688 | ZmEREB41 | NA |
| Test | PACMAD | c4_shifted | TCCAAGTCMAASCAT | GRMZM2G146688 | ZmEREB41 | NA |
| Test | PACMAD | c4_shifted | GCCCGAGGAAGGAAT | GRMZM2G147880 | ZmWRKY113 | NA |
| Test | PACMAD | c4_shifted | GAAAAAGTCCAAGCG | GRMZM2G147880 | ZmWRKY113 | NA |
| Test | PACMAD | c4_shifted | TCRCDCACAGCGCAC | GRMZM2G147880 | ZmWRKY113 | NA |
| Test | PACMAD | c4_shifted | ATTGAACTCATCAAT | GRMZM2G148467 | ZmSBP21 | NA |
| Test | PACMAD | c4_shifted | GTGAGGGAAGGAGCG | GRMZM2G148467 | ZmSBP21 | NA |
| Test | PACMAD | c4_shifted | CSACTGGACCGGGCC | GRMZM2G148467 | ZmSBP21 | NA |
| Test | PACMAD | c4_shifted | CCCGTACGTATCYCC | GRMZM2G148467 | ZmSBP21 | NA |
| Test | PACMAD | c4_shifted | CCCRCGGTCACGAGC | GRMZM2G148467 | ZmSBP21 | NA |
| Test | PACMAD | c4_shifted | AAATCTTTTGTGGCG | GRMZM2G148467 | ZmSBP21 | NA |
| Test | PACMAD | c4_shifted | GMKCKCGCGCASCAA | GRMZM2G156006 | ZmEREB207 | NA |
| Test | PACMAD | c4_shifted | GGGCCCACYYGGWAG | GRMZM2G156006 | ZmEREB207 | MEME-2 |
| Test | PACMAD | c4_shifted | GGCYCACAKKCAVYC | GRMZM2G156006 | ZmEREB207 | NA |
| Test | PACMAD | c4_shifted | TKGASAMAAGCSCAC | GRMZM2G163813 | ZmSBP19 | NA |
| Test | PACMAD | c4_shifted | CGTGAAYAAAATGGG | GRMZM2G163813 | ZmSBP19 | NA |

| | | | | | | |
|---|---|---|---|---|---|---|
| Test | PACMAD | c4_shifted | SMGGRGTCGGAGTCA | GRMZM2G163813 | ZmSBP19 | NA |
| Test | PACMAD | c4_shifted | GGSCMABTAKCAGCT | GRMZM2G163813 | ZmSBP19 | NA |
| Test | PACMAD | c4_shifted | AGACCRTTTTCTTTT | GRMZM2G163975 | NA | Zm00001d027846 |
| Test | PACMAD | c4_shifted | CAGACAGACAAATAC | GRMZM2G163975 | NA | NA |
| Test | PACMAD | c4_shifted | CTACTTCYATTTGGT | GRMZM2G163975 | NA | NA |
| Test | PACMAD | c4_shifted | GAGAACAATTTGTAC | GRMZM2G163975 | NA | NA |
| Test | PACMAD | c4_shifted | CAGTGACTGATCGAT | GRMZM2G163975 | NA | NA |
| Test | PACMAD | c4_shifted | GCTGTTTAAAGCTGC | GRMZM2G178102 | ZmHB25 | NA |
| Test | PACMAD | c4_shifted | GAAAGAGCCGAAATC | GRMZM2G178102 | ZmHB25 | NA |
| Test | PACMAD | c4_shifted | CTCGCTTTGACAGAG | GRMZM2G178102 | ZmHB25 | NA |
| Test | PACMAD | c4_shifted | CTTGAYGCCAAAAGA | GRMZM2G178102 | ZmHB25 | NA |
| Test | PACMAD | c4_shifted | AAAGCGTGMTGTAAA | GRMZM2G178102 | ZmHB25 | NA |
| Test | PACMAD | c4_shifted | CAAACGGWAAATATC | GRMZM2G178261 | ZmGRF1 | CCA1 |
| Test | PACMAD | c4_shifted | TGTCCCCTTGTGGCC | GRMZM2G180406 | ZmbHLH32 | NA |
| Test | PACMAD | c4_shifted | GCCCCCTCCCCRATT | GRMZM2G180406 | ZmbHLH32 | NA |
| Test | PACMAD | c4_shifted | GTATACGATCTAAMC | GRMZM2G180406 | ZmbHLH32 | NA |
| Test | PACMAD | c4_shifted | GGCACCAACATCRTC | GRMZM2G180406 | ZmbHLH32 | NA |
| Test | PACMAD | c4_shifted | GTGGTTGGAAGGAGG | GRMZM2G312419 | ZmMYB60 | NA |
| Test | PACMAD | c4_shifted | GTTGGACTCCCCAAA | GRMZM2G312419 | ZmMYB60 | NA |
| Test | PACMAD | c4_shifted | TTTCTWGGYTGTYTT | GRMZM2G374986 | ZmGLK30 | NAC078 |
| Test | PACMAD | c4_shifted | GDCGCAATTAGCTTG | GRMZM2G377217 | ZmWRKY98 | NA |
| Test | PACMAD | c4_shifted | SCCCKGCRAWYTCGC | GRMZM2G398506 | ZmWRKY1 | NA |
| Test | PACMAD | c4_shifted | TCACCBCTCACTGCC | GRMZM2G398506 | ZmWRKY1 | NA |
| Test | PACMAD | c4_shifted | CTTTTGTTTTGCATG | GRMZM2G399072 | ZmEREB130 | NA |
| Test | PACMAD | c4_shifted | AKACATGAGGATCTR | GRMZM2G469304 | NA | NA |
| Test | PACMAD | c4_shifted | GRACGRAGTAAAATT | GRMZM2G469304 | NA | NA |
| Test | PACMAD | c4_shifted | CCATGTGAGCGTAGC | GRMZM5G880069 | ZmWRKY109 | NA |
| Test | PACMAD | c4_shifted | TCTGCATBCGGTGGT | GRMZM5G880069 | ZmWRKY109 | NA |
| Test | PACMAD | c4_shifted | TAGGTTTGAAGTCWG | GRMZM5G880069 | ZmWRKY109 | NA |
| Test | PACMAD | c4_shifted | CCYCAACAGGTCATC | GRMZM5G880069 | ZmWRKY109 | NA |
| Test | PACMAD | c4_shifted | GCGGGAGGGACCCGG | GRMZM5G880069 | ZmWRKY109 | BAD1 |
| Test | PACMAD | c4_shifted | ACTCATGTCTTTGAC | GRMZM5G880069 | ZmWRKY109 | NA |
| Test | PACMAD | c4_shifted | TACTCCGMGGGCGTC | GRMZM5G880069 | ZmWRKY109 | NA |
| Test | PACMAD | c4_shifted | TTTCTWGGYTGTYTT | GRMZM5G887276 | ZmGLK48 | NA |
| Test | PACMAD | not_syntenic | TGTGTGTAAATGTGG | GRMZM2G008234 | ZmEREB114 | NA |

| | | | | | | |
|---|---|---|---|---|---|---|
| Test | PACMAD | not_syntenic | TGAAGAGAATGTTAT | GRMZM2G008234 | ZmEREB114 | NA |
| Test | PACMAD | not_syntenic | TCCAAGTCCAASCAT | GRMZM2G008234 | ZmEREB114 | NA |
| Test | PACMAD | not_syntenic | GCGAGGGVGAGAGAG | GRMZM2G013391 | ZmWRKY111 | RAMOSA1 |
| Test | PACMAD | not_syntenic | CGGAGTGGTTAGGGS | GRMZM2G013391 | ZmWRKY111 | NA |
| Test | PACMAD | not_syntenic | GCGTATCCGCACATG | GRMZM2G013391 | ZmWRKY111 | NA |
| Test | PACMAD | not_syntenic | RTGTSCATGATTGGC | GRMZM2G013391 | ZmWRKY111 | ATHB-7 |
| Test | PACMAD | not_syntenic | GYGTGTGCACGGGCG | GRMZM2G013391 | ZmWRKY111 | NA |
| Test | PACMAD | not_syntenic | CCSCRCCGCYGGCCG | GRMZM2G014653 | ZmNAC109 | NA |
| Test | PACMAD | not_syntenic | CGYGTCGCCGTCGCS | GRMZM2G014653 | ZmNAC109 | ereb17 |
| Test | PACMAD | not_syntenic | CCTCCCVTTCCGWGC | GRMZM2G015666 | ZmbHLH33 | NA |
| Test | PACMAD | not_syntenic | AGVCGAWGCTRAAWA | GRMZM2G015666 | ZmbHLH33 | NA |
| Test | PACMAD | not_syntenic | TCTCCATTTCACCVC | GRMZM2G015666 | ZmbHLH33 | NA |
| Test | PACMAD | not_syntenic | CARACSGGYYTCCAC | GRMZM2G028980 | ZmARF16 | NA |
| Test | PACMAD | not_syntenic | CARACSGGYYTCCAC | GRMZM2G035405 | ZmARF18 | NA |
| Test | PACMAD | not_syntenic | GTAACCACGCGGCGG | GRMZM2G060544 | ZmLBD5 | NA |
| Test | PACMAD | not_syntenic | CAAGGGTTCCATGCT | GRMZM2G061734 | ZmSBP5 | NA |
| Test | PACMAD | not_syntenic | TGAACATGGCSSTGG | GRMZM2G061734 | ZmSBP5 | NA |
| Test | PACMAD | not_syntenic | CGCAYCKTCCATTTC | GRMZM2G061734 | ZmSBP5 | NA |
| Test | PACMAD | not_syntenic | CCSCCKCCCCC | GRMZM2G061734 | ZmSBP5 | GLYMA-03G162700 |
| Test | PACMAD | not_syntenic | CCTCCCVTTCCGWGC | GRMZM2G082586 | ZmbHLH105 | NA |
| Test | PACMAD | not_syntenic | AGVCGAWGCTRAAWA | GRMZM2G082586 | ZmbHLH105 | NA |
| Test | PACMAD | not_syntenic | TCTCCATTTCACCVC | GRMZM2G082586 | ZmbHLH105 | NA |
| Test | PACMAD | not_syntenic | GAGACGCGGCTGGGC | GRMZM2G102845 | ZmARF20 | NA |
| Test | PACMAD | not_syntenic | AAVADGACGAGGMAA | GRMZM2G102845 | ZmARF20 | NA |
| Test | PACMAD | not_syntenic | GGKTCSCGTGCGCGG | GRMZM2G104078 | ZmNAC101 | NA |
| Test | PACMAD | not_syntenic | SACCCCGTAGATGGC | GRMZM2G114613 | ZmLSD4 | NA |
| Test | PACMAD | not_syntenic | CGGGTCTGCGGCGCC | GRMZM2G114613 | ZmLSD4 | NA |
| Test | PACMAD | not_syntenic | CCCCGTCSCSGCCGA | GRMZM2G114613 | ZmLSD4 | NA |
| Test | PACMAD | not_syntenic | CKCCGGCTCCGGTAG | GRMZM2G114613 | ZmLSD4 | ereb17 |
| Test | PACMAD | not_syntenic | CMCCVCCRTGGCCGG | GRMZM2G137541 | ZmbHLH151 | NA |
| Test | PACMAD | not_syntenic | AATGCCTCGATBGGC | GRMZM2G137541 | ZmbHLH151 | NA |
| Test | PACMAD | not_syntenic | TGGCASTGTCWGGGA | GRMZM2G137541 | ZmbHLH151 | NA |
| Test | PACMAD | not_syntenic | GGTTTGCTTTGCYGT | GRMZM2G137541 | ZmbHLH151 | NA |
| Test | PACMAD | not_syntenic | GCRGGCAGGGAATAA | GRMZM2G137541 | ZmbHLH151 | NA |
| Test | PACMAD | not_syntenic | CACACACAMACACAM | GRMZM2G146688 | ZmEREB41 | NA |

| | | | | | | |
|---|---|---|---|---|---|---|
| Test | PACMAD | not_syntenic | GCTTGTGCAGTGAGT | GRMZM2G146688 | ZmEREB41 | NA |
| Test | PACMAD | not_syntenic | TCSGTTGGTGDGSGC | GRMZM2G151689 | ZmC3H18 | NA |
| Test | PACMAD | not_syntenic | GATGCTSWMYCTAGG | GRMZM2G156006 | ZmEREB207 | NA |
| Test | PACMAD | not_syntenic | CCGGCATCTSATGTG | GRMZM2G156006 | ZmEREB207 | NA |
| Test | PACMAD | not_syntenic | GTCAAGGCCGGGAGG | GRMZM2G312419 | ZmMYB60 | NA |
| Test | PACMAD | not_syntenic | CCADCCAAACCMATC | GRMZM2G312419 | ZmMYB60 | NA |
| Test | PACMAD | not_syntenic | GGVAAGGGAAGSCTG | GRMZM2G312419 | ZmMYB60 | NA |
| Test | PACMAD | not_syntenic | YTYTCTCYCTCCTCY | GRMZM2G312419 | ZmMYB60 | NA |
| Test | PACMAD | not_syntenic | AAAAACAAATC | GRMZM2G312419 | ZmMYB60 | NA |
| Test | PACMAD | not_syntenic | CYGGTAGTGGTTGGS | GRMZM2G312419 | ZmMYB60 | NA |
| Test | PACMAD | not_syntenic | CCTTTTGGTDYTCCC | GRMZM2G374986 | ZmGLK30 | NA |
| Test | PACMAD | not_syntenic | TGYCATCATGC | GRMZM2G374986 | ZmGLK30 | NA |
| Test | PACMAD | not_syntenic | CCTCCTMCGGCARGC | GRMZM2G374986 | ZmGLK30 | NA |
| Test | PACMAD | not_syntenic | GCATGCCAAGCTTCC | GRMZM2G374986 | ZmGLK30 | NA |
| Test | PACMAD | not_syntenic | TTATCACTTTCAGTT | GRMZM2G374986 | ZmGLK30 | NA |
| Test | PACMAD | not_syntenic | GCAATCGATCGCAGG | GRMZM2G377217 | ZmWRKY98 | NA |
| Test | PACMAD | not_syntenic | AGACGGTGATGATTC | GRMZM2G377217 | ZmWRKY98 | NA |
| Test | PACMAD | not_syntenic | AGTGCTCTTTTTGCG | GRMZM2G377217 | ZmWRKY98 | NA |
| Test | PACMAD | not_syntenic | GCAAAACCTGCCCCG | GRMZM2G377217 | ZmWRKY98 | NA |
| Test | PACMAD | not_syntenic | TGTRCACACGGCACA | GRMZM2G377217 | ZmWRKY98 | NA |
| Test | PACMAD | not_syntenic | GAGVGCATCCTRAAA | GRMZM2G377217 | ZmWRKY98 | NA |
| Test | PACMAD | not_syntenic | CTGAAAACGAGTGAA | GRMZM2G377217 | ZmWRKY98 | NA |
| Test | PACMAD | not_syntenic | AACRAAACGTGATYA | GRMZM2G377217 | ZmWRKY98 | NA |
| Test | PACMAD | not_syntenic | CTCCAGTGTCYAYGC | GRMZM2G377217 | ZmWRKY98 | NA |
| Test | PACMAD | not_syntenic | AGGCAGATCTCTGAC | GRMZM2G377217 | ZmWRKY98 | NA |
| Test | PACMAD | not_syntenic | AGATCGGTCGATTGG | GRMZM2G377217 | ZmWRKY98 | NA |
| Test | PACMAD | not_syntenic | TGCCCCGYCTGAAAA | GRMZM2G377217 | ZmWRKY98 | NA |
| Test | PACMAD | not_syntenic | CCTGGCYCCTGCCYC | GRMZM2G529859 | ZmYAB8 | NA |
| Test | PACMAD | not_syntenic | CSTGCCTGCCTGCGC | GRMZM2G529859 | ZmYAB8 | NA |
| Test | PACMAD | not_syntenic | GGTTCCACCWTGCCC | GRMZM2G529859 | ZmYAB8 | NA |
| Test | PACMAD | not_syntenic | SAAGAYCCRCAYGCA | GRMZM2G529859 | ZmYAB8 | NA |
| Test | PACMAD | not_syntenic | ABAGTCTCGAGATGC | GRMZM2G529859 | ZmYAB8 | NA |
| Test | PACMAD | not_syntenic | TCCGAGGACCCGGCG | GRMZM5G813892 | ZmGLK61 | NA |
| Test | PACMAD | not_syntenic | AAACGAAACGAGMGG | GRMZM5G813892 | ZmGLK61 | NA |
| Test | PACMAD | not_syntenic | CCTTTTGGTDYTCCC | GRMZM5G887276 | ZmGLK48 | NA |

| | | | | | | |
|---|---|---|---|---|---|---|
| Test | PACMAD | not_syntenic | TGYCATCATGC | GRMZM5G887276 | ZmGLK48 | NA |
| Test | PACMAD | not_syntenic | CCTCCTMCGGCARGC | GRMZM5G887276 | ZmGLK48 | NA |
| Test | PACMAD | not_syntenic | GCATGCCAAGCTTCC | GRMZM5G887276 | ZmGLK48 | NA |
| Test | PACMAD | not_syntenic | TTATCACTTTCAGTT | GRMZM5G887276 | ZmGLK48 | NA |
| Test | PACMAD_BEP | c4_shifted | ACAGTGCCACAACTG | GRMZM2G008234 | ZmEREB114 | NA |
| Test | PACMAD_BEP | c4_shifted | GTCGCGATCGCAGCC | GRMZM2G013391 | ZmWRKY111 | NA |
| Test | PACMAD_BEP | c4_shifted | GYATACGATCACGGG | GRMZM2G013391 | ZmWRKY111 | NA |
| Test | PACMAD_BEP | c4_shifted | TRGCCTAATCACRCC | GRMZM2G015666 | ZmbHLH33 | NA |
| Test | PACMAD_BEP | c4_shifted | CSGCGGGGCCCASSA | GRMZM2G018487 | ZmWRKY107 | PK19717.1 |
| Test | PACMAD_BEP | c4_shifted | CGATGGACGGCCGCG | GRMZM2G018487 | ZmWRKY107 | NA |
| Test | PACMAD_BEP | c4_shifted | GGGCCCACCKCCCCC | GRMZM2G018487 | ZmWRKY107 | NA |
| Test | PACMAD_BEP | c4_shifted | GCCCCAAGCGCAAAG | GRMZM2G024247 | ZmHMG4 | NA |
| Test | PACMAD_BEP | c4_shifted | ACCTCACTAGCTGCG | GRMZM2G024247 | ZmHMG4 | NA |
| Test | PACMAD_BEP | c4_shifted | GAGRGAGAGRGKGGG | GRMZM2G060544 | ZmLBD5 | NA |
| Test | PACMAD_BEP | c4_shifted | GTCGCTGTACGATTT | GRMZM2G060544 | ZmLBD5 | NA |
| Test | PACMAD_BEP | c4_shifted | GTCTCTCTCGAGACA | GRMZM2G062244 | ZmHB107 | NA |
| Test | PACMAD_BEP | c4_shifted | TRGCCTAATCACRCC | GRMZM2G082586 | ZmbHLH105 | NA |
| Test | PACMAD_BEP | c4_shifted | CGACGTGAAAACGAC | GRMZM2G098813 | ZmLFY2 | NA |
| Test | PACMAD_BEP | c4_shifted | GRGCACGCTCGCKCG | GRMZM2G098813 | ZmLFY2 | NA |
| Test | PACMAD_BEP | c4_shifted | TGCACGCCTAATTAT | GRMZM2G121309 | ZmIAA16 | NA |
| Test | PACMAD_BEP | c4_shifted | WTGCATGGMAKGSAW | GRMZM2G123909 | ZmGATA35 | NA |
| Test | PACMAD_BEP | c4_shifted | AYTGCACCCACCTCM | GRMZM2G133398 | ZmABI1 | NA |
| Test | PACMAD_BEP | c4_shifted | CGSSCGCCACGTGTG | GRMZM2G133398 | ZmABI1 | NA |
| Test | PACMAD_BEP | c4_shifted | GGCTGGCGTGGGCGG | GRMZM2G133398 | ZmABI1 | NA |
| Test | PACMAD_BEP | c4_shifted | CRCTCCCCACAGCCC | GRMZM2G133398 | ZmABI1 | NA |
| Test | PACMAD_BEP | c4_shifted | GAAGCAGGAGGTGGA | GRMZM2G133398 | ZmABI1 | NA |
| Test | PACMAD_BEP | c4_shifted | CCTCAGTCGGCGCGC | GRMZM2G133398 | ZmABI1 | NA |
| Test | PACMAD_BEP | c4_shifted | AAAGAAAGATTGAGA | GRMZM2G137541 | ZmbHLH151 | BPC1 |
| Test | PACMAD_BEP | c4_shifted | SCGGGACCACGCGGT | GRMZM2G146020 | ZmbZIP8 | NA |
| Test | PACMAD_BEP | c4_shifted | TTCATCTTYTRCTWT | GRMZM2G146688 | ZmEREB41 | NA |
| Test | PACMAD_BEP | c4_shifted | CSGCGGGGCCCASSA | GRMZM2G147880 | ZmWRKY113 | PK19717.1 |
| Test | PACMAD_BEP | c4_shifted | CGATGGACGGCCGCG | GRMZM2G147880 | ZmWRKY113 | NA |
| Test | PACMAD_BEP | c4_shifted | GGGCCCACCKCCCCC | GRMZM2G147880 | ZmWRKY113 | TCP23 |
| Test | PACMAD_BEP | c4_shifted | AACCGGATTTGATAG | GRMZM2G155123 | ZmPHD17 | NA |
| Test | PACMAD_BEP | c4_shifted | GGCTGGGTSGGTCGC | GRMZM2G155123 | ZmPHD17 | NA |

| | | | | | | |
|---|---|---|---|---|---|---|
| Test | PACMAD_BEP | c4_shifted | GACTACCAAGCATCC | GRMZM2G180406 | ZmbHLH32 | NA |
| Test | PACMAD_BEP | c4_shifted | AATGATTGGTTGATC | GRMZM2G180406 | ZmbHLH32 | ZAT6 |
| Test | PACMAD_BEP | c4_shifted | TGCCGCBGCCGCACG | GRMZM2G320827 | ZmTHX32 | NA |
| Test | PACMAD_BEP | c4_shifted | CGCCGCATGCCGCCG | GRMZM2G320827 | ZmTHX32 | NA |
| Test | PACMAD_BEP | c4_shifted | GCTGTCTTCATCTYC | GRMZM2G374986 | ZmGLK30 | NA |
| Test | PACMAD_BEP | c4_shifted | GGGTCAACAGCTCTG | GRMZM2G374986 | ZmGLK30 | WRKY63 |
| Test | PACMAD_BEP | c4_shifted | TGCTCTGACCGCATC | GRMZM2G374986 | ZmGLK30 | NA |
| Test | PACMAD_BEP | c4_shifted | CCGGTTTCGTGCCGC | GRMZM2G398506 | ZmWRKY1 | NA |
| Test | PACMAD_BEP | c4_shifted | KTGAYKTAATYAATC | GRMZM2G398506 | ZmWRKY1 | NA |
| Test | PACMAD_BEP | c4_shifted | CCCCCGGATCCCACC | GRMZM2G398506 | ZmWRKY1 | NA |
| Test | PACMAD_BEP | c4_shifted | GCCATATATAGSTYG | GRMZM2G399072 | ZmEREB130 | NA |
| Test | PACMAD_BEP | c4_shifted | ATCACCAAATCACCT | GRMZM2G399072 | ZmEREB130 | NA |
| Test | PACMAD_BEP | c4_shifted | AGTCGCRCACCGAAA | GRMZM2G399072 | ZmEREB130 | NA |
| Test | PACMAD_BEP | c4_shifted | GGCACGGCCMCCAAG | GRMZM2G399072 | ZmEREB130 | NA |
| Test | PACMAD_BEP | c4_shifted | TGCCGCBGCCGCACG | GRMZM2G427087 | ZmTHX31 | NA |
| Test | PACMAD_BEP | c4_shifted | CGCCGCATGCCGCCG | GRMZM2G427087 | ZmTHX31 | NA |
| Test | PACMAD_BEP | c4_shifted | GCATGTGACCTGCTC | GRMZM2G469304 | NA | NA |
| Test | PACMAD_BEP | c4_shifted | GCATGCCACTGTCCT | GRMZM2G469304 | NA | ABI3 |
| Test | PACMAD_BEP | c4_shifted | GCGCGGCCCACGCCC | GRMZM2G481163 | ZmTHX4 | NA |
| Test | PACMAD_BEP | c4_shifted | MGGRACMACGCGGCR | GRMZM5G821024 | ZmbZIP23 | NA |
| Test | PACMAD_BEP | c4_shifted | CCTCGHGATYCGTGY | GRMZM5G821024 | ZmbZIP23 | NA |
| Test | PACMAD_BEP | c4_shifted | GCTGTCTTCATCTYC | GRMZM5G887276 | ZmGLK48 | NA |
| Test | PACMAD_BEP | c4_shifted | GGGTCAACAGCTCTG | GRMZM5G887276 | ZmGLK48 | WRKY63 |
| Test | PACMAD_BEP | c4_shifted | TGCTCTGACCGCATC | GRMZM5G887276 | ZmGLK48 | NA |
| Test | PACMAD_BEP | not_syntenic | TCGCTAGCGGCTACT | GRMZM2G005353 | ZmYAB14 | NA |
| Test | PACMAD_BEP | not_syntenic | ACAGTGCCACAACTG | GRMZM2G008234 | ZmEREB114 | NA |
| Test | PACMAD_BEP | not_syntenic | GTCGCGATCGCAGCC | GRMZM2G013391 | ZmWRKY111 | NA |
| Test | PACMAD_BEP | not_syntenic | GYATACGATCACGGG | GRMZM2G013391 | ZmWRKY111 | NA |
| Test | PACMAD_BEP | not_syntenic | GMGGCCGGCTTCTTT | GRMZM2G013391 | ZmWRKY111 | NA |
| Test | PACMAD_BEP | not_syntenic | GBGRGGGATCTGGCC | GRMZM2G013391 | ZmWRKY111 | NA |
| Test | PACMAD_BEP | not_syntenic | TCGCTAGCGGCTACT | GRMZM2G054795 | ZmYAB1 | NA |
| Test | PACMAD_BEP | not_syntenic | GCGGCGYCSAATCCG | GRMZM2G060544 | ZmLBD5 | RAMOSA1 |
| Test | PACMAD_BEP | not_syntenic | GTCTCTCTCGAGACA | GRMZM2G062244 | ZmHB107 | NA |
| Test | PACMAD_BEP | not_syntenic | CTTTYGCTCCTCTCG | GRMZM2G081892 | ZmEREB118 | NA |
| Test | PACMAD_BEP | not_syntenic | CCTTGAAAGCTGCAT | GRMZM2G088242 | ZmHSF2 | NA |

| | | | | | | |
|---|---|---|---|---|---|---|
| Test | PACMAD_BEP | not_syntenic | CGGAAGTGTCACGGG | GRMZM2G088242 | ZmEREB118 | NA |
| Test | PACMAD_BEP | not_syntenic | CGGAAGTGTCACGGG | GRMZM2G088242 | ZmHSF2 | NA |
| Test | PACMAD_BEP | not_syntenic | ATTTGTCAAGTCAAC | GRMZM2G098813 | ZmLFY2 | NA |
| Test | PACMAD_BEP | not_syntenic | GRGCACGCTCGCKCG | GRMZM2G098813 | ZmHSF2 | NA |
| Test | PACMAD_BEP | not_syntenic | GRGCACGCTCGCKCG | GRMZM2G098813 | ZmLFY2 | NA |
| Test | PACMAD_BEP | not_syntenic | TGGGAGACTGYTCCC | GRMZM2G121309 | ZmIAA16 | NA |
| Test | PACMAD_BEP | not_syntenic | TCGTTGTTGCTAGCT | GRMZM2G121309 | ZmIAA16 | NA |
| Test | PACMAD_BEP | not_syntenic | GRACAGATCGRACT | GRMZM2G121309 | ZmIAA16 | NA |
| Test | PACMAD_BEP | not_syntenic | CAGGCAGACATSCGC | GRMZM2G121309 | ZmIAA16 | NA |
| Test | PACMAD_BEP | not_syntenic | CTGCYTTCYTTTGSA | GRMZM2G121309 | ZmIAA16 | NA |
| Test | PACMAD_BEP | not_syntenic | ABGCATGCATGCAGC | GRMZM2G121309 | ZmIAA16 | NA |
| Test | PACMAD_BEP | not_syntenic | ATMATGTRGCMAAAC | GRMZM2G121309 | ZmIAA16 | NA |
| Test | PACMAD_BEP | not_syntenic | GTGCSCCTCGTGATC | GRMZM2G121309 | ZmIAA16 | NA |
| Test | PACMAD_BEP | not_syntenic | CMACRCCACGCCAGC | GRMZM2G121309 | ZmIAA16 | NA |
| Test | PACMAD_BEP | not_syntenic | TTCACGTGGCCKCGC | GRMZM2G121309 | ZmIAA16 | NA |
| Test | PACMAD_BEP | not_syntenic | TGCACGCCTAATTAT | GRMZM2G121309 | ZmIAA16 | NA |
| Test | PACMAD_BEP | not_syntenic | TGCACGCCTAATTAT | GRMZM2G121309 | ZmLFY2 | NA |
| Test | PACMAD_BEP | not_syntenic | CYCTTTTGCTYCTTT | GRMZM2G121309 | ZmIAA16 | NA |
| Test | PACMAD_BEP | not_syntenic | CATCGCKTSYCGCGC | GRMZM2G121309 | ZmIAA16 | NA |
| Test | PACMAD_BEP | not_syntenic | TGGGAGACTGYTCCC | GRMZM2G123909 | ZmGATA35 | NA |
| Test | PACMAD_BEP | not_syntenic | TCGTTGTTGCTAGCT | GRMZM2G123909 | ZmGATA35 | NA |
| Test | PACMAD_BEP | not_syntenic | GRACAGATCGRACT | GRMZM2G123909 | ZmGATA35 | NA |
| Test | PACMAD_BEP | not_syntenic | CAGGCAGACATSCGC | GRMZM2G123909 | ZmGATA35 | ARF27 |
| Test | PACMAD_BEP | not_syntenic | CTGCYTTCYTTTGSA | GRMZM2G123909 | ZmGATA35 | NA |
| Test | PACMAD_BEP | not_syntenic | ABGCATGCATGCAGC | GRMZM2G123909 | ZmGATA35 | GLYMA-08G357600 |
| Test | PACMAD_BEP | not_syntenic | ATMATGTRGCMAAAC | GRMZM2G123909 | ZmGATA35 | NA |
| Test | PACMAD_BEP | not_syntenic | GTGCSCCTCGTGATC | GRMZM2G123909 | ZmGATA35 | NA |
| Test | PACMAD_BEP | not_syntenic | CMACRCCACGCCAGC | GRMZM2G123909 | ZmGATA35 | NA |
| Test | PACMAD_BEP | not_syntenic | TTCACGTGGCCKCGC | GRMZM2G123909 | ZmGATA35 | bHLH74 |
| Test | PACMAD_BEP | not_syntenic | CYCTTTTGCTYCTTT | GRMZM2G123909 | ZmGATA35 | NA |
| Test | PACMAD_BEP | not_syntenic | CATCGCKTSYCGCGC | GRMZM2G123909 | ZmGATA35 | NA |
| Test | PACMAD_BEP | not_syntenic | AYTGCACCCACCTCM | GRMZM2G133398 | ZmABI1 | NA |
| Test | PACMAD_BEP | not_syntenic | TGACAARTGGGCCCM | GRMZM2G133398 | ZmABI1 | NA |
| Test | PACMAD_BEP | not_syntenic | CCCGAGGCTAACGAA | GRMZM2G137541 | ZmbHLH151 | NA |
| Test | PACMAD_BEP | not_syntenic | CTTCAACAAAACTAA | GRMZM2G137541 | ZmbHLH151 | NA |

| | | | | | | |
|---|---|---|---|---|---|---|
| Test | PACMAD_BEP | not_syntenic | GSGCGWGTARATAAT | GRMZM2G137541 | ZmbHLH151 | NA |
| Test | PACMAD_BEP | not_syntenic | MTGTTGACYACCATT | GRMZM2G137541 | ZmbHLH151 | NA |
| Test | PACMAD_BEP | not_syntenic | TTTAAAAGCTTAATG | GRMZM2G137541 | ZmbHLH151 | NA |
| Test | PACMAD_BEP | not_syntenic | CTTGCAGGCCACTTC | GRMZM2G137541 | ZmbHLH151 | NA |
| Test | PACMAD_BEP | not_syntenic | AAGCTAGATTCGAGC | GRMZM2G137541 | ZmbHLH151 | NA |
| Test | PACMAD_BEP | not_syntenic | GGGGCAAAAGCGTGT | GRMZM2G137541 | ZmbHLH151 | NA |
| Test | PACMAD_BEP | not_syntenic | TATTGCATGGGYGGG | GRMZM2G137541 | ZmbHLH151 | NA |
| Test | PACMAD_BEP | not_syntenic | CASCTCBCGYCCACA | GRMZM2G146020 | ZmbZIP8 | NA |
| Test | PACMAD_BEP | not_syntenic | SCGGGACCACGCGGT | GRMZM2G146020 | ZmbZIP8 | NA |
| Test | PACMAD_BEP | not_syntenic | ACACAGATCCCYATG | GRMZM2G146688 | ZmEREB41 | ANT |
| Test | PACMAD_BEP | not_syntenic | GTGTGGGTCAAMTGC | GRMZM2G146688 | ZmEREB41 | WRKY63 |
| Test | PACMAD_BEP | not_syntenic | GACACCCCTCCATGT | GRMZM2G148467 | ZmSBP21 | NA |
| Test | PACMAD_BEP | not_syntenic | AATAGGAGAGAGAAA | GRMZM2G155123 | ZmPHD17 | NA |
| Test | PACMAD_BEP | not_syntenic | CAGCYAGTACTGAAG | GRMZM2G377217 | ZmWRKY98 | NA |
| Test | PACMAD_BEP | not_syntenic | CCGGTTTCGTGCCGC | GRMZM2G398506 | ZmWRKY1 | NA |
| Test | PACMAD_BEP | not_syntenic | CCCCCGGATCCCACC | GRMZM2G398506 | ZmWRKY1 | NA |
| Test | PACMAD_BEP | not_syntenic | CCTKGCATGCATGYG | GRMZM2G399072 | ZmEREB130 | NA |
| Test | PACMAD_BEP | not_syntenic | GGCACGGCCMCCAAG | GRMZM2G399072 | ZmEREB130 | NA |
| Test | PACMAD_BEP | not_syntenic | GCGCGGCCCACGCCC | GRMZM2G481163 | ZmTHX4 | NA |
| Test | PACMAD_BEP | not_syntenic | GTTCAAATTTCGAAT | GRMZM5G803308 | ZmMYB112 | NA |
| Test | PACMAD_BEP | not_syntenic | CACTKCCCCACGWAC | GRMZM5G803308 | ZmMYB112 | NA |
| Test | PACMAD_BEP | not_syntenic | TCCGTTGGCCGTTAC | GRMZM5G803308 | ZmMYB112 | NA |
| Test | PACMAD_BEP | not_syntenic | TGRCAYGTGGG | GRMZM5G821024 | ZmbZIP23 | NA |
| Test | PACMAD_BEP | not_syntenic | CTGACCGAAGTGACC | GRMZM5G850129 | ZmGRF11 | NA |
| Control | PACMAD | c4_shifted | CGCACGGGAACGGTC | GRMZM2G033846 | NA | NA |
| Control | PACMAD | c4_shifted | CCGCGTTGACATGTC | GRMZM2G033846 | NA | NA |
| Control | PACMAD | c4_shifted | CCYTGCACCGGACTC | GRMZM2G033846 | NA | NA |
| Control | PACMAD | c4_shifted | CGTCCTCGATGACGG | GRMZM2G033846 | NA | NA |
| Control | PACMAD | c4_shifted | GCCTATAARTASCRG | GRMZM2G073969 | NA | NA |
| Control | PACMAD | c4_shifted | CCWTTGAAACAGTCY | GRMZM2G162505 | NA | NA |
| Control | PACMAD | c4_shifted | TAAAGATCGGMTTCG | GRMZM2G442008 | NA | NA |
| Control | PACMAD | c4_shifted | GTAGTGTGTRATGTT | GRMZM2G442008 | NA | NA |
| Control | PACMAD | c4_shifted | TGTACSACDTAGTTT | GRMZM2G442008 | NA | NA |
| Control | PACMAD | c4_shifted | CYGTGATYRTACCAT | GRMZM2G442008 | NA | NA |
| Control | PACMAD | c4_shifted | TCATTTGAAAGTGGA | GRMZM2G442008 | NA | NA |

| | | | | | | |
|---|---|---|---|---|---|---|
| Control | PACMAD | c4_shifted | CATGGAGTTCATYRG | GRMZM2G479596 | NA | NA |
| Control | PACMAD | c4_shifted | RCCTCTTGATC | GRMZM2G479596 | NA | NA |
| Control | PACMAD | c4_shifted | TCTTKGKTCCAGGTG | GRMZM2G479596 | NA | NA |
| Control | PACMAD | c4_shifted | CCCCTCGRGTCGGCC | GRMZM2G479596 | NA | NA |
| Control | PACMAD | c4_shifted | SGGGTCAGAGAGGAS | GRMZM2G479596 | NA | NA |
| Control | PACMAD | c4_shifted | GACGCCATGAACTGG | GRMZM2G479596 | NA | NA |
| Control | PACMAD | not_syntenic | TCAGSTGCKGCSGBG | GRMZM2G145027 | NA | NA |
| Control | PACMAD | not_syntenic | TCAAAGTTGGTAGCC | GRMZM2G162505 | NA | NA |
| Control | PACMAD | not_syntenic | TTTTCWGTTTGAAAA | GRMZM2G162505 | NA | NA |
| Control | PACMAD | not_syntenic | ATCTTCTGGAAGTG | GRMZM2G162505 | NA | NA |
| Control | PACMAD | not_syntenic | GCAAGATTTGTTTGC | GRMZM2G162505 | NA | NA |
| Control | PACMAD | not_syntenic | CWCYCTCTCTCTCAT | GRMZM2G479596 | NA | RAMOSA1 |
| Control | PACMAD | not_syntenic | ACTGCCAAACCGTAC | GRMZM2G479596 | NA | NA |
| Control | PACMAD_BEP | c4_shifted | ATCMATGCAGCASYR | GRMZM2G073969 | NA | NA |
| Control | PACMAD_BEP | c4_shifted | GCYTCTAGGGTTCCG | GRMZM2G097059 | ZmMADS7 | NA |
| Control | PACMAD_BEP | c4_shifted | GAGGTTCCATCCGGA | GRMZM2G097059 | ZmMADS7 | NA |
| Control | PACMAD_BEP | c4_shifted | SCCTCGACCTGTCGA | GRMZM2G097059 | ZmMADS7 | NA |
| Control | PACMAD_BEP | c4_shifted | GAAGAAGTGGTTCAA | GRMZM2G442008 | NA | NA |
| Control | PACMAD_BEP | c4_shifted | GAATCCAATTCAGAT | GRMZM2G442008 | NA | NA |
| Control | PACMAD_BEP | c4_shifted | GAAGGCCCTGCTGAG | GRMZM2G442008 | NA | NA |
| Control | PACMAD_BEP | c4_shifted | AAATGRTGTAAAGAA | GRMZM2G442008 | NA | NA |
| Control | PACMAD_BEP | c4_shifted | TTCCTGAAAAGTCAG | GRMZM2G442008 | NA | NA |
| Control | PACMAD_BEP | not_syntenic | GCYTCTAGGGTTCCG | GRMZM2G097059 | ZmMADS7 | NA |
| Control | PACMAD_BEP | not_syntenic | CTCTCTCTCTCTCTC | GRMZM2G097059 | ZmMADS7 | BPC5 |
| Control | PACMAD_BEP | not_syntenic | CGRTCCTCGCYCCCC | GRMZM2G173341 | NA | NA |
| Control | PACMAD_BEP | not_syntenic | CGCCGCGGCGAGCGC | GRMZM2G366434 | ZmEREB206 | NA |
| Control | PACMAD_BEP | not_syntenic | GAAGAAGTGGTTCAA | GRMZM2G442008 | NA | NA |
| Control | PACMAD_BEP | not_syntenic | GAAGGCCCTGCTGAG | GRMZM2G442008 | NA | NA |
